# Unveiling dynamics behind Glioblastoma Multiforme single-cell data heterogeneity

**DOI:** 10.1101/2023.04.25.538198

**Authors:** Marcos Guilherme Vieira Junior, Adriano Maurício de Almeida Côrtes, Flávia Raquel Gonçalves Carneiro, Nicolas Carels, Fabrício Alves Barbosa da Silva

**Affiliations:** Graduate Program in Computational and Systems Biology, Instituto Oswaldo Cruz (IOC), Oswaldo Cruz Foundation (FIOCRUZ), Rio de Janeiro, Brazil; Department of Applied Mathematics, Institute of Mathematics, Federal University of Rio de Janeiro, Rio de Janeiro, Brazil; Systems Engineering and Computer Science Program, COPPE, Federal University of Rio de Janeiro, Rio de Janeiro, Brazil; Center of Technological Development in Health (CDTS), FIOCRUZ, Rio de Janeiro, Brazil; Laboratório Interdisciplinar de Pesquisas Médicas, IOC, FIOCRUZ, Rio de Janeiro, Brazil; Program of Immunology and Tumor Biology, Brazilian National Cancer Institute(INCA), Rio de Janeiro, Brazil; Laboratory of Biological System Modeling, CDTS, FIOCRUZ, Rio de Janeiro, Brazil; Scientific Computing Program, FIOCRUZ, Rio de Janeiro, Brazil

## Abstract

Glioblastoma Multiforme (GBM), a brain tumor distinguished for its aggressive nature, presents heterogeneity that stems from multifaceted mechanisms, such as genetic mutations, epigenetic modifications, genomic instability, and selective pressures. We reason that the aggressiveness of GBM enables even a limited dataset to serve as a representative sample within the domain of cancer attractors, allowing our sample to reflect a representative trajectory within the attractor’s domain. Utilizing single-cell RNA sequencing data, we proceed with a detailed analysis of GBM’s cellular landscape. Rooted in characteristics observed in stochastic systems, we considered factors like genomic instability to introduce a level of noise or unpredictability, thereby characterizing the cancer dynamics through stochastic fixed points. These fixed points, derived from centroids obtained through various clustering methods, were rigorously verified for method sensitivity. This methodological foundation, wherein sample and time averages are equivalent, assigns paramount interpretative value to the data cluster centroids, aiding both in parameter fitting and subsequent stochastic simulations. This scenario is supported by the compelling correlation found between centroids of experimental and simulated datasets. The use of stochastic modeling to compute the Waddington landscape enriched our analysis of GBM’s cellular heterogeneity and provided a visual framework for validating the centroids and standard deviations as accurate characterizations of potential cancer attractors. Specifically, this approach allowed us to assess the compatibility between data dispersion and the corresponding basin of attraction, thereby bridging molecular-level variations and systems-level dynamics. We also examined the stability and transitions between these attractors, revealing a potential interplay between subtypes and potentially uncovering factors that drive cancer recurrence and progression. By connecting molecular mechanisms related to cancer heterogeneity with statistical properties of gene expression dynamics we expect to set the stage for the development of potential diagnostic tools and pave the way for personalized therapeutic interventions.

## Introduction

Notwithstanding the significant advancements in understanding and therapeutics, cancer continues to be a predominant global cause of mortality (***Who, 2022***). Glioblastoma multiforme (GBM; Glioblastoma Multiforme) stands as the most common and aggressive brain tumor, characterized by an average survival time of 15 months and a roughly 10% probability of achieving a 5-year overall survival (***Sasmita et al., 2017***; ***Gallego, 2015***). Single-cell RNA sequencing (scRNA-seq) has spotlighted the pronounced heterogeneity inherent in GBM (***Neftel et al., 2019***; ***Patel et al., 2014***). Intriguingly, this level of complexity in scRNA-seq data is not exclusive to GBM. Many cancers manifest similar intricate patterns in their sequencing data (***Tirosh et al., 2016***; ***Kim et al., 2018***), underpinning a broader challenge in oncology research. With such heterogeneity possibly driving the aggressiveness of these malignancies (***McGranahan and Swanton, 2017***; ***Marusyk et al., 2014***), pressing questions emerge regarding the underlying dynamics. Foremost among these are the interpretations of observed data distributions, their ensuing consequences, and the patterns encapsulated within the data. Do we behold stable fixed points, limit cycles, or might there be a deeper, more convoluted narrative at play?

Extensive research has delved into the complexities of carcinogenesis, advancing discourses not only on driver and passenger mutations but also on the profound influences of epigenetics (***Vogelstein et al., 2013***; ***Shen and Laird, 2013***; ***Esteller, 2008***). In the intricate landscape of gene regulatory networks (GRN) dynamics, pivotal studies have elucidated the alignment between cell types or subtypes and stable states, often termed ‘attractors’ (***Huang, 2011***; ***Li et al., 2016***). Concurrently, certain oscillatory cellular processes — integral to diverse functions such as circadian rhythms (***Takahashi, 2016***), cell cycle progression (***Pomerening et al., 2005***), and NF-*κ*B signaling in response to inflammation (***Nelson et al., 2004***) — are closely tied to limit cycles in GRNs. Yet, the foundational principles guiding transitions between these states, as well as the mechanisms by which a system traverses within an attractor’s domain — a notion sometimes framed as a ‘cancer attractor’ (***Huang et al., 2009***) — remain areas of vigorous exploration and research.

The intricacies of tumor evolutionary trajectories further underscore a pressing need for understanding. As cancer progresses or counteracts therapeutic measures, the dynamic shifts in tumor subclonal architectures come to the forefront (***Álvarez-Arenas et al., 2019***). Traditional linear evolution models may inadequately capture the complexities of tumor evolution. In certain scenarios, a dominant clone proliferates, leading to a predominantly homogeneous tumor composition. Conversely, other situations present coexisting subclonal populations, suggesting a more branched evolutionary pattern than a purely linear trajectory (***Burrell et al., 2013***). In light of these challenges, the increasing availability of omics data presents an opportunity for deeper investigations. For instance, studies have identified subtypes of GBM, a fundamental leap forward in our understanding of the disease and a critical step for choosing the applying treatment (***Sasmita et al., 2017***; ***Verhaak et al., 2010***; ***Phillips et al., 2006***; ***Jiao et al., 2012***). Yet, as the horizon of our knowledge expands, questions about the evolutionary pathways of these subtypes and the dynamic interplay underpinning their classifications persist, beckoning deeper investigations (***Wang et al., 2017***; ***Sidaway, 2017***; ***Rajapakse et al., 2020***).

This evolving scenario necessitates a shift in perspective. Instead of remaining anchored in reductionist perspectives, there’s a pressing call to embrace a more systemic approach. This perspective not only provides a comprehensive view of cell type and functionality dynamics but is also reinforced by studies like (***Strauss et al., 2021***). This systemic thinking is present in contemporary paradigms that depict cancer as a nuanced series of events leading to a ‘state disease’ (***Hanahan, 2022***), rather than being merely the fallout of isolated mutations. In this context, we define the *state* as the biochemical milieu of a cell, signifying the evolutionary trajectory of states within a complex system. Contrasting starkly with earlier models that portrayed cancer progression as linear, this perspective revels in understanding the multifaceted dynamics of the disease within a broader, multidimensional context (***Huang, 2011***). A vivid analogy for this conceptual shift can be traced back to C. Waddington’s 1957 metaphor, where cellular differentiation is visualized as a sphere traversing an (epi)genetic^1^ landscape of peaks and valleys (***Waddington, 1957***), with the zeniths representing the undifferentiated phenotype.

In the Waddington (epi)genetic landscape, a differentiated state is achieved through developmental paths influenced by both intrinsic cellular factors — as the cell’s historical events, such as lineage, gene expression patterns, and epigenetic modifications (***Reik, 2007***) — and by extrinsic factors like tissue-level perturbations or environmental influences. According to this perspective, cancer might be associated with one or more (defective or malignant) attractor states in the (epi)genetic landscape. These malignant attractors can either pre-exist hidden within the landscape or emerge due to genetic and epigenetic alterations. In both scenarios, they are undesirably reached due to the inherent stochastic fluctuations of biological systems at the biomolecular scale (***Huang et al., 2009***). Using the Waddington (epi)genetic landscape as a conceptual framework allows us to more clearly understand the extent of heterogeneity within GBM. Cellular differentiation, as envisioned by Waddington, is deeply intertwined with the nuances of cancer progression. If we extend this framework, it’s clear that the various paths and trajectories through which a cell might journey — and eventually culminate in a malignant phenotype — are likely shaped by a combination of genetic, epigenetic, and environmental forces. This understanding reinforces the need to decipher the intricate details behind single-cell data, especially since heterogeneity is both an outcome and an influencer of tumor dynamics. As we delve deeper, the landscape metaphor becomes more than just a conceptual tool; it provides a practical framework that guides our investigation of GBM. By examining how and why certain trajectories or states become prominent in GBM, we lay the groundwork for exploring the underlying factors and mechanisms that drive cellular heterogeneity and what this might signify for our broader understanding of cancer evolution.

From this exploration, it becomes evident that cellular heterogeneity in single-cell data raises crucial questions. These questions concern the interplay between epigenetic regulation, genomic stability, selective pressures, and the inherent GRN governing cancer cell behavior. A thorough investigation into these aspects offers fresh perspectives into the evolving landscape of GBM. One of the primary sources of heterogeneity is genomic instability (***Burrell et al., 2013***). This instability, marked by a high mutation rate at the DNA level, results in a cellular milieu teeming with diversity (***Guo et al., 2023***). While genomic instability is a significant contributor to heterogeneity, it represents only one facet of the complexity. Genetic alterations, combined with epigenetic mechanisms, lead to a diverse range of cellular responses, amplifying the heterogeneity and adding layers to our understanding. This way, the dynamics underlying heterogeneity are molded by additional factors that add to its complexity.

Selective pressures exert a significant influence on cancer heterogeneity. Rather than being merely passive, these pressures actively shape the outcome of the diversity introduced by genomic instability (***Greaves and Maley, 2012***). They curate the cellular environment, favoring the emergence of stable cellular states and modulating the dynamics within the (epi)genetic landscape. Such stable states, also termed ‘attractors’, represent characteristic cellular phenotypes. Trajectories within the phase space tend to gravitate towards these attractors, shaped by the so-called ‘basins of attraction’. In light of this, the interplay between the chaos of genomic instability and the order introduced by selective pressures, as revealed through the lens of clonal evolution, offers a detailed understanding (***Greaves and Maley, 2012***). It’s not merely about heterogeneity; it underscores a structured heterogeneity, reflecting a complex dynamic infused with order.

This narrative implies a representation that captures a cell’s transformative journey toward a malignant state. In this work, we propose a modeling of this representation that aligns with the snapshot-like data from scRNA-seq (figure 1). Stemming from a structured stochastic limit cycle (***Biswas et al., 2021***) (figure 1-I), the cellular trajectory is subject to perturbations from genetic mutations, epigenetic regulations, and selective pressures. It’s noteworthy that even in this initial state, some dimensions, potentially associated with marker genes, may already exhibit a fixed-point dynamic. As these forces act, the initial trajectory undergoes alterations, manifested as an increase in stochastic noise that broadens the oscillatory boundary, or ‘stochastic tube’ (figure 1-II). This amplification in noise effectively populates the state space, thereby increasing the relevance of mean values as accurate representations of the cellular dynamics. Finally, the cell adopts more erratic behavior, eventually taking on characteristics resembling a random wandering around a fixed point (figure 1-III). This transformation might occur for multiple genes, culminating in a tumor’s genetic and phenotypic variability, termed intratumor heterogeneity. Importantly, this evolving subclonal architecture is dynamic, undergoing continual shifts throughout disease progression and thus presenting challenges for both diagnostics and therapeutic strategies.

**Figure 1.**
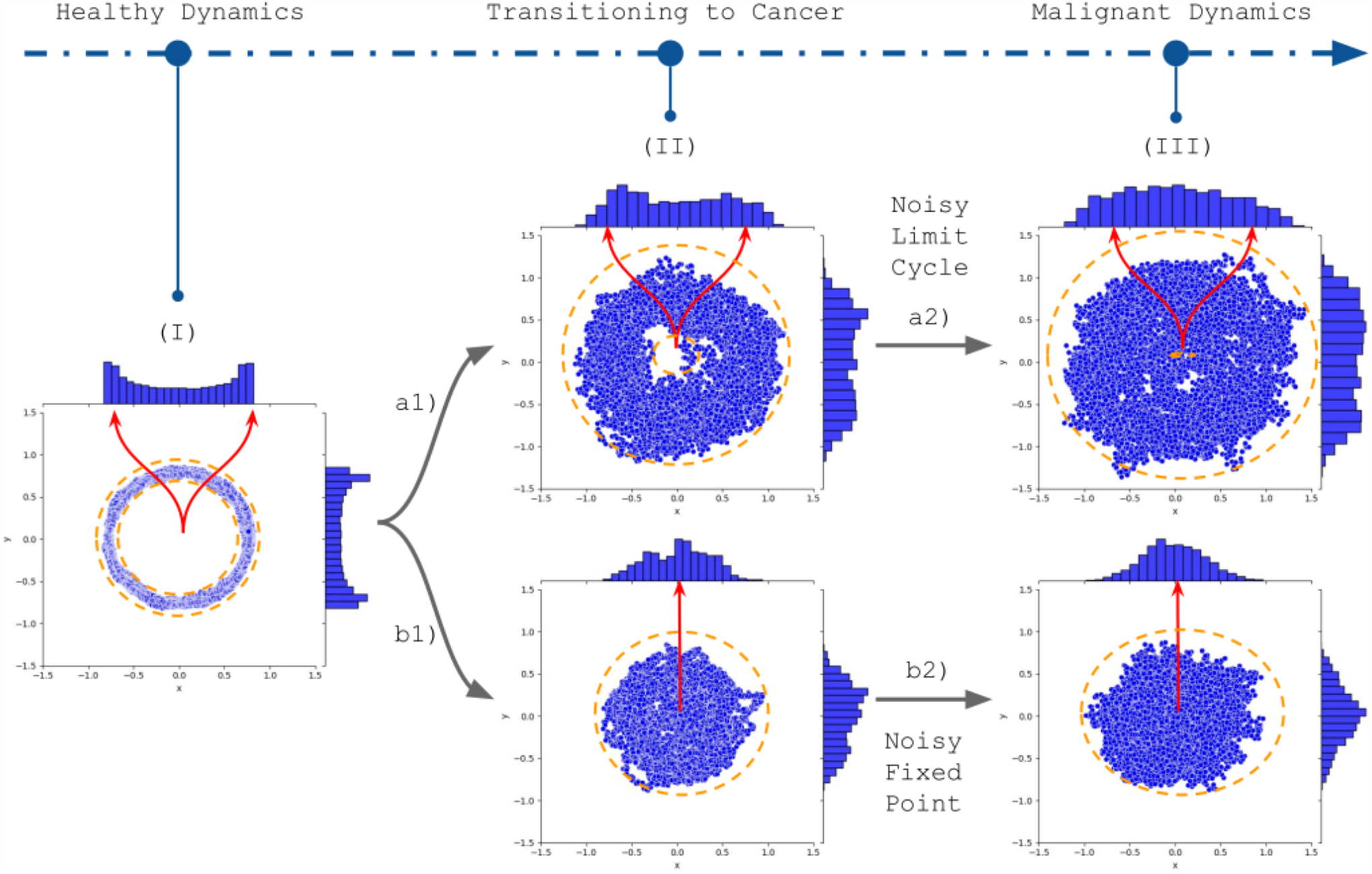
The image depicts our hypothesis about a cellular transition from a ‘healthy’ limit cycle to a malignant state, a transformation that would enlighten the dynamics underlying the single-cell RNA sequencing data heterogeneity. The ‘initial state’ (I) represents a stable cellular trajectory. However, it’s important to note certain dimensions—potentially related to marker genes— might already display a fixed-point dynamic. The outcomes of genetic, epigenetic, and microenvironmental alterations are depicted through directional arrows (a1 to a2, and b1 to b2). The upward trajectory (a1 to a2) underscores an expanded oscillatory boundary, typically referred to as a stochastic tube. Notably, the initial limit cycle distribution (I) is characterized by two non-central peaks, which witness a reduced prominence due to augmented stochastic fluctuations (II), resulting in an irregular distribution resembling oscillations around a stable point (III). In contrast, the descending trajectory (b1 to b2) emphasizes an amplified fluctuation envelope coupled with parameter shifts associated with a Hopf bifurcation, consolidating the dynamics of a malignancy state around a fixed point. The fixed point histogram displays a peak aligned with the fixed points, highlighting the relevance of features such as clustering centroids in capturing the nuanced hallmarks of malignant transformation.

Models aiming to represent a cell’s transformative journey, especially through the lens of the (epi)genetic landscape, have emerged in the scientific community (***Wang et al., 2010, 2011***; ***Ferrell, 2012***; ***Li and Wang, 2013a, 2014, 2015***; ***Li et al., 2016***; ***Verd et al., 2014***). These models provide insights into the temporal flow and stability of cellular states, enhancing our understanding of cancer’s evolutionary pathways. However, they face challenges owing to their reliance on time series data for parameter estimation—a requirement that’s often elusive given the intricacies of capturing real-time biological processes. Recognizing these limitations, techniques have emerged that infer temporality (the ordered succession of events) through pseudotime (***Saelens et al., 2019***). Though pseudotime provides a promising avenue to infer transitions between attractors using scRNA-seq data, it’s not without its challenges. Events like mutations, deregulation of the cell cycle (***Witkiewicz et al., 2022***), and cellular heterogeneity can potentially obscure the interpretation of pseudotime trajectories, further complicating the characterization of cancer dynamics. Yet, such complexities underscore the relevance of a stochastic modeling approach.

Temporal data acquisition presents inherent challenges, giving rise to numerous theoretical constructs. Among these, a central theme is a system’s statistical behavior, tracking its evolution in phase space over time. A distinguishing feature of some of these perspectives is the convergence of time averages to ensemble averages (***Moore, 2015***), which is particularly transformative for snapshot-centric data like scRNA-seq. This approach offers an avenue to bypass the intricacies of temporal sampling. Expanding on this, some theories emphasize examining specific components within a system, such as basins of attraction (***Palmer, 1982***; ***Mauro and Smedskjaer, 2014***). Translating this to the GBM context, the distinct aggressiveness levels of each subtype can be interpreted as different moments in a broader cellular narrative. We propose that the ensemble averages derived from the distinct GBM subtypes might be akin to time averages of a singular representative trajectory (1). This correlation might be especially evident in specific gene space dimensions, likely marker genes. In combination with the fixed points modeling, such intertwining of theoretical insights with observed biological phenomena has inspired our hypothesis, driving us toward a comprehensive modeling perspective.

Given these challenges and insights, our hypothesis is grounded in the idea of a system that over an extended timeframe will navigate through all accessible states within the bounds of a cancer attractor with a consistent likelihood. While this assumption might simplify certain complexities, it establishes a robust foundation to probe the stochastic essence of single-cell data. Such a model gathers the concepts of an equilibrium state with inherent variability, offers a streamlined approach to estimate parameters, and interprets the variability presented in data. However, should there be complications in estimating these parameters, we may need to reassess the foundation of our hypothesis. Validity would largely depend on comparing simulated outputs with experimental findings, a step that could significantly enhance our hypothesis’s reliability.

In validating our hypothesis, we employ experimental GBM scRNA-seq data to generate a computational model, incorporating principles from (epi)genetic landscape modeling, Langevin’s dynamics (***Gillespie, 2000***), and Hill Functions ((***Santillán, 2008***)) with modifications for the GRN dynamics. Our methodology centers on dynamic modeling, integrating raw data with insights derived from the underlying biological processes and mechanisms. In the proposed context, the cancer attractor concept suggests a propensity for specific cancer subtypes, recognizable as distinctive regions in phase space. Additionally, while exploring the gene expression space it is expected that some areas might be inaccessible due to biological constraints. Our assumptions include metric transitivity, which means that two points in phase space can be connected by a shortest path in the gene expression space. This phenomenon aligns with the idea that intrinsic cellular noise enhances phase space exploration in cancer cells, diminishing the barriers between different basins of attraction (***Huang et al., 2009***). Building upon these insights into the gene expression space and its constraints, we propose a stochastic model in silico, aiming to quantify the (epi)genetic landscapes derived from scRNA-seq data of four GBM subtypes: Classical, Mesenchymal, Proneural, and Neural. This model also contemplates interactions inherent to a GBM-specific GRN.

Our modeling efforts promise more than theoretical research. For instance, they hint at tangible avenues for interpreting the intricacies of genomic instability related to cancer heterogeneity. As various mechanisms that contribute to genomic instability imprint distinct molecular signatures (***Guo et al., 2023***; ***Burrell et al., 2013***), by exploring the statistical behaviors explained in our approach, we could potentially correlate mechanisms underlying these signatures, offering a chance to identify novel therapeutic targets (***da Costa et al., 2022***). In other words, we aimed to associate alterations in statistical properties observed in scRNA-seq data with molecular-level events that modify the system’s exploration of possible states. Distinguishing these unique molecular imprints would allow us to forge stronger connections between the ‘geometry of heterogeneity’ seen in single-cell data and distinct instability mechanisms, offering a more enriched understanding of tumor biology. Consequently, this statistical property could be viewed as a consequence of the progression of the malignant state.

In light of these insights and the challenges they present, this report seeks to integrate the theoretical foundations of the Waddington (epi)genetic landscape with the wealth of data emerging from single-cell technologies. By leveraging a stochastic dynamics model, our aim was to unravel the intricate mechanisms that sculpt the heterogeneity inherent to the GBM landscape. Our methodology provides a data-driven quantification of the (epi)genetic landscape specific to GBM and its respective subtypes. Additionally, we probed the statistical dynamics of our in silico model, establishing a framework for subsequent inquiries and potential practical applications. Furthermore, it contributes to developing studies on a biological system’s possible long-term behavior and stability.

## Methods and Materials

Historically, biology has utilized models to interpret complex biological phenomena. Traditional approaches often employed model organisms or cell lines for in vitro studies. However, with recent advancements in computational methods and mathematics, there has been a notable shift towards abstract mathematical models. These models, acting as approximations, allow researchers to navigate and hypothesize within controlled digital environments, simulating the complexities of biological systems. While in silico experiments may not conclusively validate general biological principles, they offer insights into hypothesis outcomes and afford preliminary validation via induction (***Voit, 2019***).

One of the most intricate and fundamental processes in biology that benefits from such computational modeling is gene regulation. Given its inherent complexity and dynamic nature, mathematical modeling has emerged as an indispensable tool to elucidate the nuances of this process.

### Model background

The regulation of gene expression is a complex process that involves multiple layers and mechanisms (***Huang, 2011***). One possible measurement of gene expression is the number of messenger RNA (mRNA) molecules that effectively translate into proteins. The expression profile is a dynamic feature, changing in time according to cell types and characteristics. By considering a vector **X** = (*X*_1_, *X*_2_, …, *X*_*N*_) with *N* being the total number of variables and each vector component representing the quantification of mRNA molecules, the cell state can be modeled using system dynamics theory. The basis of the gene regulation dynamics modeling is its associated deterministic differential equations system, an autonomous system of ordinary differential equations 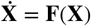, containing information about the temporal trajectory driven by the interaction forces between each of its components (***Meister et al., 2014***).

There are several possible functions to parameterize the interactions of a nonlinear model, but the common choice is sigmoidal functions. Among them, the Hill function is the most frequent as it has many experimentally observed required characteristics (***Santillán, 2008***). An example of a driving force *F* using Hill functions can be seen in ***Wang et al***. (***2010***), with a more general form described by:

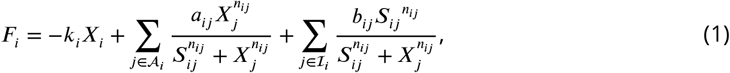

where, for each gene *i*, represented by the component *X*_*i*_, the index sets 𝒜_*i*_ and ℐ_*i*_ represent the genes that interact with gene *i* through activation and inhibition, respectively. The value *j* represents the edge that bridges the regulation of transcription factors interacting with their target gene promoters. Note that in the case of self-activation or self-inhibition, one has *i* ∈ 𝒜_*i*_ or *i* ∈ ℐ_*i*_, respectively. The parameter *S* denotes the value where the Hill function reaches its maximum inclination, *n* represents the intensity of the transition, *a* is the activation coefficient, *b* is the inhibition coefficient, and *k* is the self-degradation constant. When *a* is a self-activation, or *b* is a self-inhibition parameter, they will be denoted by *sa* and *sb*, respectively. The parameters *k, a*, and *b* have units of time^−1^, while the remaining parameters are dimensionless.

As seen in the equation (1), in the most general form, the gene activation (*a*) and inhibition (*b*) parameters may vary for each interaction or even as a function of time (non-autonomous system). In addition, sigmoid coefficients may (i) be constant, (ii) vary according to interactions or some proposed functions, or (iii) present a time dependence. In this model, the gene inhibition is given by constraining its basal expression, as can be seen in the positive sign of the inhibition term, with the higher inhibitions obtained by lower values of *b*.

Although using deterministic differential equations to study general behavior is adequate, biological systems are inherently stochastic (***Meister et al., 2014***). Thermal fluctuations and varying conditions affect the likelihood of interactions and give these systems a probabilistic nature. Consequently, the number of molecules over time follows a fluctuating, noisy pattern. A common way to model this stochasticity is through the Chemical Master Equation (CME), a Markovian model that captures the probabilistic nature of molecular interactions (***Gillespie, 2000***). However, solving the CME can be computationally challenging, especially for large systems. An alternative is to use Langevin dynamics (***Gillespie, 2000***), which serves as an approximation of the CME, described by a deterministic term *F* (*x*) and a stochastic term *ξ*(*t*). In Langevin dynamics, we can treat *ξ*(*t*) as random fluctuations (without memory) due to its much smaller timescale compared to *F* (*x*). The dynamics became:

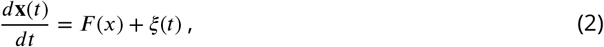

where *x*(*t*) is the expression level as a function of time (implicit dependence) relative to random variables of *X, F* (*x*) is the deterministic term representing regulation due to network interactions, and *ξ*(*t*) is the stochastic term with average ⟨*ξ*(*t*) ⟩ = 0 and amplitude given by its autocorrelation function ⟨*ξ*_*i*_(*t*)*ξ*_*j*_(*t*^′^)⟩ = 2*D*_*ij*_ *δ*_*ij*_ *δ*(*t* − *t*^′^) (***Li and Wang, 2013a***), with *D* being the diffusion coefficient and representing a fluctuation scale factor.

With the presence of fluctuations, probability distributions model gene expression levels, and the temporal evolution *p*(*x, t*) is described by the Fokker-Planck equation (eq 35). This equation provides a continuous approximation to the CME (***Meister et al., 2014***), capturing molecular diffusion kinetics across an epigenetic landscape. However, since the Fokker-Planck equation’s driving force and noise components are unknown, an alternative approach to studying the system is to consider a stochastic differential equation (SDE) with possible deterministic and stochastic components. In this context, we consider a system described by (***Li et al., 2016***):

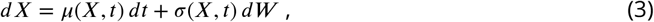

with drift *μ*(*X, t*), noise parameter *σ*(*X, t*) and Wiener standard process *dW*. We assumed *μ*(*X, t*) = *F* (*X*) by considering the drift due to the driven force. The noise *σ*(*X, t*) can be divided into two major contributions: (i) intrinsic, which is related to the system’s internal dynamics, and (ii) extrinsic, which is due to the effects of the environment/microenvironment. We considered a multiplicative noise *σ*(*X, t*) = *g*(*X, t*), so that the fluctuations may be described by different timescales and constrained by the defined regulation function. The complete definition of the system is given both by the parameters of the deterministic and the stochastic components, with the specifics of the multiplicative noise and the regulation function to be detailed later in this report.

### scRNA-seq data

While theoretical models provide a conceptual procedure to understand gene regulation, capturing accurate data remains paramount. In recent years, scRNA-seq has emerged as a powerful tool to study gene expression profiles at the individual cell level, enabling the investigation of cellular heterogeneity and the identification of distinct cell subpopulations. This technology has been particularly valuable for studying GRNs. It provides insights into the complex interactions between genes and the possible regulatory mechanisms that drive cell-type-specific gene expression patterns.

#### GBM and single-cell data

Transitioning from the broader picture of scRNA-seq to its specialized utility, GBM stands out as a compelling case study. GBM, renowned for its profound cellular heterogeneity, exemplifies the challenges researchers grapple with when studying complex disease landscapes (***Patel et al., 2014***). Considering the nuances of cellular evolution in tumor environments, a deeper dive into the roles of selective pressures and genomic instability in shaping GBM’s intricate heterogeneity became imperative. Yet, it is precisely this complexity that makes GBM a fertile ground for scRNA-seq explorations. Single-cell datasets, in this context, serve as “temporal snapshots,” chronicling the multifaceted expression patterns of GBM’s cellular ensemble at distinct timeline intervals. Although these snapshots might appear isolated, a deeper dive reveals they often resonate with the broader dynamism governing cellular behavior. Figure 1 encapsulates this idea of the richness of information each “snapshot” brings to the table. It suggests that while each scRNA-seq dataset offers a temporally distinct perspective, collectively, they can traverse the entire phase space, capturing the essence of GBM’s intricate dynamics over time. Such insights emphasize scRNA-seq’s transformative potential in unveiling the dynamics underpinning tumor heterogeneity.

#### GBM dataset

Our analytical endeavors are anchored on data curated and analyzed by ***Darmanis et al***. (***2017***). This comprehensive dataset encompasses single-cell resolution RNA sequencing outputs from patients diagnosed with diverse GBM subtypes. The study scrutinized tumor heterogeneity, contrasting the tumor core with its periphery. This dataset aggregates samples from four patients, all diagnosed with primary GBM and characterized by a negative *IDH1* signature (indicating an absence of mutations in the *IDH* gene). Following stringent quality control measures, the dataset retained information from 3589 cells, including various cell types from the central nervous system such as vascular, immune, neuronal, and glial cells.

The analytical framework employed by ***Darmanis et al***. (***2017***) identified cellular clusters from the dimensionality reduction with tSNE, layered over a dissimilarity matrix. Subsequent clustering via the k-means algorithm refined cellular groupings. A meticulous gene expression audit identified the signature genes of each cluster, the results of which were cross-referenced against healthy tissue data to chart cellular identities. Residual clusters, predominantly localized to the tumor core and marked by heightened expression of genes like *EGFR* and *SOX9*, were cataloged as neoplastic. Further validation against independent datasets from both healthy brain tissue and GBM bulk RNA-Seq reinforced the study’s findings. An intriguing observation was the conspicuous absence of astrocytes within the tumor core. Furthermore, a consistent expression profile for tumor cells residing in the peripheral zones was documented across all patient samples (***Darmanis et al., 2017***).

### GRN construction and implementation

Utilizing the scRNA-seq data from ***Darmanis et al***. (***2017***), our research transitions into its computational modeling phase. The central goal is to create a representative model of the GRNs to understand the cellular nuances of GBM’s various subtypes and the inherent heterogeneity they exhibit. In the subsequent section, we detail the methodology that forms the foundation of this computational framework.

#### Biological criteria and methodological approach

The challenge we embraced was formulating a GRN that captures the essential dynamics behind GBM subtypes and heterogeneity. The GRN needs to be composed of regulatory interactions (edges) between the genes (vertices) of the system supposed to be representative of the case under analysis (GBM). To begin this endeavor, the first step involved setting up clear biological criteria to guide the curation of these interactions. Our primary focus revolved around molecular mechanisms that directly or indirectly affect the number of mRNA molecules. Our approach embarked on an initial survey of interactions, both direct and indirect. *Direct interactions*, highlighted at the transcriptional level, are characterized by the binding of transcription factors (TF) to their target gene promoters. These interactions directly affect the amount of mRNA and are represented by a direct connection between the transcription factor vertex and the vertex representing the targeted gene. Conversely, *indirect interactions* encompass mechanisms that modulate the TF’s ability to bind to a gene promoter, like the ubiquitination of a TF culminating in its degradation. Unlike direct interactions, the biological effects of indirect interactions are not immediate, and their consequences on the number of mRNAs need to be evaluated before they can be adequately represented within the model. As we delve into the complexities of these interactions, it becomes evident that a structured approach is needed to assemble the GRN.

To construct this survey of interactions, we began by pinpointing genes and markers pivotal for GBM subtypes (***Sasmita et al., 2017***). The subtype classification was anchored in the schema presented by ***Verhaak et al***. (***2010***). We then employed the MetaCore (***Analytics, 2019***) platform to search for interactions among these genes and markers. Specifically, we used the MetaCore transcription regulation network construction algorithm to build the network, identifying new vertices that bridge the initial gene list (provided in Supplementary Table rede_GenesList.xlsx). However, the output generated consisted of several disjoint subnetworks, with the initial genes scattered among them. We extracted the initial and additional genes from these subnetworks to address this issue and created a new input list. This new list was then used to generate a connected network, ensuring that all the genes of interest were included in a single structure (Figure S3). With the GRN in place, the next step was to cross-reference it with the scRNA-seq expression data.

To achieve this alignment, an algorithm (***Vieira, 2023a***) in R (***R Development Core Team, 2008***) was developed to match the scRNA-seq expression data with the network generated by MetaCore. The data preprocessing was performed using the Seurat package (***Satija et al., 2015***), from which a *sctransform* normalization was applied to reduce technical bias and recover biologically significant distributions (***Hafemeister and Satija, 2019***; ***Lab, 2022***). Cell cycle effects were not removed, as such information may lose its accuracy in tumor cells (***Witkiewicz et al., 2022***). The algorithm then performed the following operations: (i) selected interactions classified as *Transcription Regulation*, (ii) intersected the genes of the network with those present in the data, and (iii) removed the genes that were associated with null values, even if they were present in the network. As a result, the network may change with variations in data, whether due to the selected cell types or patient IDs. With this adaptive network established, the subsequent step involved refining our working model to more accurately represent the cell types and their spatial localization within the GBM landscape.

Building upon our initial network, we adopted the cell type classifications proposed in ***Darmanis et al***. (***2017***). These authors identified various cell types present in GBM samples, such as astrocytes, oligodendrocytes, and neurons, among others, through clustering and other methods. Centering our study on neoplastic cells nested within the tumor core, we delved into the dispersion of expression value for every patient consolidated in our dataset (Figure S1). To increase the probability of sampling over the entire phase space of GBM, we decided to obtain the landscape for all patients simultaneously instead of analyzing individual patients separately. This approach allows for better characterization of GBM by observing attractors related to the four subtypes of GBM, while still reflecting the specific characteristics of each individual. By filtering for neoplastic cells located in the tumor core, we avoided incorporating the different features observed for neoplastic cells present in the periphery of the tumor. The filtration process culminated in an aggregate of 1027 cells, offering a robust foundation for downstream analyses.

To enable the automatic integration of the network with mathematical and computational models, we developed an algorithm, available in the provided code repository (***Vieira, 2023b***), which simplifies the process. The algorithm takes the network as input in a tabular format, representing the connectivity list between each vertex. It then processes the input and converts the tabular data structure into two directional graphs (digraphs), one for activation interactions and another for inhibition interactions, each represented by its adjacency matrix. With these transformed data structures, equation (1) can be written as:

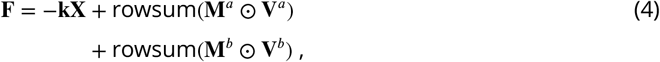

with **k** = diag(*k*_1_, …, *k*_*N*_) a diagonal matrix, **M**^*a*^ the activation matrix with entries (**M**^*a*^)_*ij*_ = *a*_*ij*_, **M**^*b*^ the inhibition matrix with entries (**M**^*b*^)_*ij*_ = *b*_*ij*_, **V**^*a*^ the activation Hill functions matrix with entries

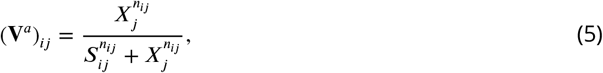

**V**^*b*^ the inhibition Hill functions matrix with entries

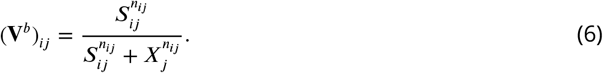

The ⊙ denotes the Hadamard product (element-wise matrix product), and rowsum(⋅) returns the vector with the row-wise sums of the matrix.

To observe the effects of perturbations, we represented the original adjacency matrices positions corresponding to activation (*a*), self-activation (*sa*), inhibition (*b*), and self-inhibition (*sb*) parameters as arbitrary values using symbolic computation. This method allowed further replacement of these symbols with numerical values. For example, *a* = *sa* = *b* = *sb* = 1 during the parameter’s estimation, and a broader parameter space exploration for studying the basins’ stability. We could achieve the same by using multiplicative factors for each matrix element when *i* = *j* or *i* ≠ *j*.

### Data analysis: Dynamics underlying heterogeneity

Upon the construction of our GRN, we have delineated the requisite genes and established the targeted scenario suitable for an in-depth investigation into GBM dynamics. The diverse clusters within our dataset underscore the heterogeneity of tumor evolution. This heterogeneity, especially when viewed through single-cell data, hints at the complex evolutionary trajectories of gene expressions.

At the heart of this exploration is the concept of the cancer attractor (***Huang et al., 2009***). This paradigm posits that, despite genetic differences and the evident heterogeneity, cancer cells often gravitate towards a common state. It is this theoretical backbone that justifies our application of dynamic systems theory. Cellular states, acting as non-linear evolving trajectories in a multi-dimensional space (***Huang, 2011***), are not merely random paths but rather can be viewed as potential courses directed towards these cancer attractors, converging at *basins of attraction*.

Given the challenge of real-time tracking, our focus shifts to snapshot-like data, which reflects cellular states at distinct temporal intervals. Such snapshots can be construed as potential distributions surrounding these cancer attractors for each GBM subtype. This interpretation allows studying system dynamics indirectly by observing the data variability (***Weinreb et al., 2018***). Representations akin to Figure 1, then, not only enhance our understanding of the underlying dynamics but also equip us with refined methodologies for data interpretation and parameter estimation. This integrated approach allows for a more nuanced perspective on the observed heterogeneity, paving the way for an optimized analytical framework.

In light of the cancer attractor concept, which underpins the dynamics evident in our Figure 1 hypothesis, the premise that ensemble and time averages converge becomes central. For example, for aggressive malignancies like GBM, mutation accumulation and disease progression lead to a swifter exploration of phase space. It’s imperative to note that not all cells are destined to traverse every available state over extended time intervals. Our hypothesis applies to the system’s intrinsic components (‘basins of attraction’), as delineated in ***Palmer*** (***1982***) and expanded upon in later studies such as (***Mauro et al., 2007***; ***Mauro and Smedskjaer, 2014***). Given GBM’s inherent aggressiveness, it’s posited that the heterogeneity discerned from our spatial samples provides a panoramic view of this attractor’s topography. The cellular subsets within the spatial samples are believed to span a substantial segment of the attractor.

To initiate, we discern clusters within the data (refer to Section scRNA-seq data), considering the centroids of these clusters as approximations of equilibrium states. With this presumption, the driving forces (i.e., expression level change rates) near cluster centroids are assumed negligible. Consequently, the system can be characterized by:

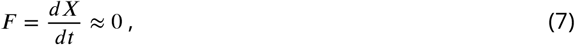

which enables the system’s stability characterization through parameter estimation (see Section GRN edges and parameter estimation).

Our characterization of clusters relies on dimensions associated with four pivotal GBM subtype markers: *EGFR* (classic), *IDH1* (proneural), *NEFL* (neural), and *CD44* (mesenchymal) (***Sasmita et al., 2017***). Unlike genes linked with the cell cycle or prone to high variability, these markers typically exhibit constrained expression ranges, ensuring reliability for clustering and landscape visualization. While many clusters likely represent distinct cell types and subtypes, others may be spurious due to artifacts, noise, or external factors. Some clusters could also reflect metastable states, short-lived configurations within the phase space. This necessitates careful analysis and validation in interpreting clustering outcomes within the GRN framework.

To operationalize the aforementioned considerations, we employed Mathematica (***Inc., 2018***) to analyze the gene expression data from ***Darmanis et al***. (***2017***). We applied a dimensionality reduction (t-SNE) to identify the clusters and then two clustering methods, k-means and Neighborhood Contraction (NbC). K-means is a popular and computationally efficient algorithm partitioning data into spherical groups (***Inc., 2023d***). NbC is a density-based method that identifies clusters of varying shapes and densities without a prior cluster number definition (***Inc., 2023e***).

### Mapping the landscape: Subtypes distributions

Given GBM’s complex landscape, it’s crucial to define the framework for our study. This landscape can be visualized as a hierarchical structure, where each layer or ‘envelope’ captures different cellular dynamics. At a high level, the landscape encapsulates cell-type attractors. Delving deeper, it reveals the intricacies of dynamics within cell types, highlighting subtypes. An even more granular approach would shed light on processes within these subtypes, such as metabolism.

Our analysis targets the intermediary level — that of dynamics among subtypes, situated within the broader GBM ‘basin of attraction’. This choice enables us to provide detailed insights while maintaining a manageable computational scope. Through our modeling, distinct basins of attraction have been identified, emphasizing GBM’s inherent heterogeneity. These basins reflect factors such as genomic instability, epigenetic regulations, and selective pressures driving the observed diversity.

We applied the central limit theorem and the law of large numbers, considering the centroids of the clusters as means of Gaussian distributions and representing gene expression levels associated with potential GBM subtypes. Additionally, we computed and stored each cluster’s standard deviation based on the respective averages, alongside the proportions of cells they contain.

Therefore, we modeled gene expression as a Gaussian distribution given by the equation:

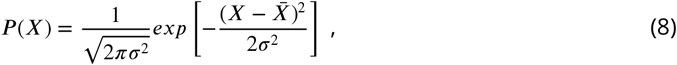

with 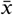 representing the sample mean and *σ* the (unbiased) sample standard deviation.

Gene interactions exhibit intricate dynamics, with alterations in one gene influencing others. Nevertheless, in steady-state systems, these interactions are reflected in the statistical distribution, enabling independent consideration of each gene (mean-field approximation). By focusing on the moments that depict the resultant distributions from the system’s collective interactions, we can manage a more tractable analysis, capturing the essential behavior of the stochastic dynamics.

The mean-field approximation helps to compute the Waddington (epi)genetic landscape, enabling the study of state transitions and the probability of observing specific gene expression distributions. This approximation allows each attractor *α* probability to be approximated by (***Li and Wang, 2013a***):

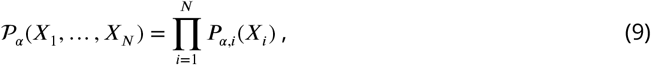

with *i* the index of genes, *N* the number of genes, and *P*_*α,i*_ defined as (8), each one with sample mean 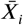 and sample standard deviation *σ*_*i*_.

The probability of a cell being in some state will be given by the steady-state probability ℙ_*ss*_ (***Li and Wang, 2013a***):

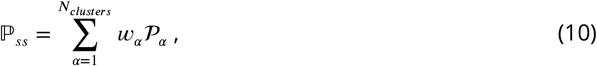

where *w*_*α*_ is the percentage of cells in each attractor, and *N*_*clusters*_ is the maximum number of attractors (clusters) found by the method.

The total steady-state probability (equation (10)) can be estimated for both experimental and simulated data. For the experimental data, by clustering the scRNA-seq data and for the simulated data by allowing sufficient time for the in silico system to evolve from sampled initial conditions towards steady states. Once we obtain the steady-states, the process is the same as for scRNA-seq data.

Computing steady-state probabilities enables studying the system’s global behavior using (epi)genetic landscapes. This theory is based on the flow theory of nonequilibrium dynamical systems, as discussed in ***Wang*** (***2015***). Obtaining steady-state probability ℙ_*ss*_ through sampling rather than an analytical solution of the Fokker-Planck equations leads to the *populational landscape*. This landscape *U* is defined as derived from the negative logarithm of the total probability of the system (***Li and Wang, 2013a***):

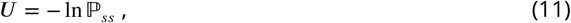

with *U* a dimensionless potential that quantifies the probabilities of states and their transitions. Higher probability states correspond to lower potentials (greater depths), while the barriers between the basins of attraction are related to the time spent in each state.

In cases where the system is in equilibrium, with detailed balance preserved, the Boltzmann relation holds. For these cases, the landscape corresponds to the equilibrium probability. However, in cases out of equilibrium, as considered here, rates of transitions between states don’t balance each other. This causes trajectories not to follow the gradient but to be characterized by the presence of probability flux, as discussed in ***Wang et al***. (***2010***).

Despite these complexities, the (epi)genetic landscape can still provide valuable insights into the behavior of systems out of equilibrium. By investigating the nonequilibrium (epi)genetic landscape, we expect to gain insights into the impact of stochastic fluctuations and transient states, which can help guide further investigations and develop more accurate models for the system’s behavior.

### in silico dynamics and simulated landscape

Having delineated the intricate (epi)genetic landscape of GBM data, we transition to the construction of an in silico dynamical model. This endeavor seeks to simulate a compatible landscape and evaluate the emergent properties of the in silico system in comparison to our initial observations. Central to this modeling process is the accurate representation of gene interactions within the GRN. To achieve this, we employ regulation functions, with a particular emphasis on Hill functions.

#### Specifying dynamics: Regulation functions

Hill functions are widely used for modeling GRN interactions, accounting for the influence of activators and inhibitors on regulation strength. Their steepness reflects the gene expression’s sensitivity to variations in regulatory molecule concentrations. They are typically characterized by constant coefficients for each interaction, resulting in a uniform regulation strength when applied to all target genes of the same TF. As monotonic functions, they exhibit increasing regulation strength in response to increasing TF concentrations. These features can lead to obstacles in modeling cancer heterogeneity.

Cancer heterogeneity implies cells might present different regulations. In other words, the same TF concentration might lead to different regulation strengths for different cells. For example, one gene may exhibit high expression for a specific number of TF molecules while another presents low expression. Typical features of Hill functions cannot model this dependence on the target gene.

Another important aspect is that they might face challenges modeling complex regulatory mechanisms when the network is incomplete. For example, gene expression levels are typically biologically constrained. It could be due to internal regulatory interaction or even microenvironment responses. In any case, both constraints are typically intricate to implement (***Strogatz, 2001***). One reason is the need to know a priori all interactions that could constrain gene expression levels. Another is that even knowing these constraints, it would still need to implement a very complex model with complicated network dynamics.

To overcome these limitations, we introduce modified Hill functions, which provide a flexible framework for modeling the relationships between genes and regulators, allowing experimental data to constrain the dynamics. We propose the modified Hill functions given by:

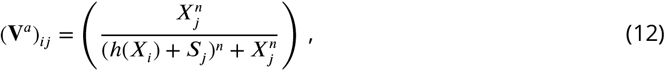

for activation, in place of (5), and by:

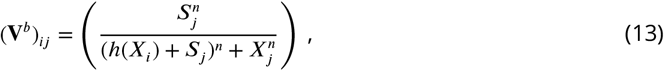

for inhibition, in place of (6). The index *i* is for the vertex related to the target, *j* for the vertex affecting the target, and *h* is a modifier function used to obtain shapes with desired properties.

To determine the best model, we tested several options for *h*, including the original model with *h*(*x*) = 0. The proposed possibilities were:

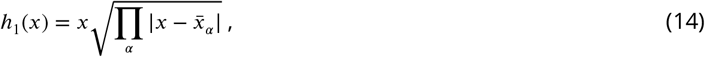

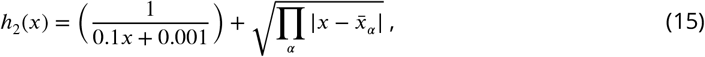

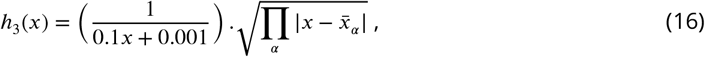

where 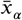 is the gene average considering each attractor *α*. The first term of equations (15) and (16) was chosen to ensure appropriate behavior for values close to zero with the constants in the denominator empirically verified to optimize the results.

In addition, we considered:

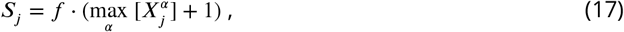

where *f* is a proportionality constant and max_*α*_ 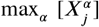 represents the transcription factor with the highest expression value among all attractors. Adding a unit to the base value of *S* ensures that the proportionality constant can adjust cases with only zeroes and avoids null denominators. This modification sets the value as a fraction of the maximum gene expression for all attractors found. This assumption is a biological simplification, proposing that regulation intensity is proportional to the highest equilibrium value of the transcription factor. This approach helps capture the effects of weaker transcription factor levels when using a single sigmoid function to represent regulation intensity. Utilizing lower levels could lead to a rapid increase in the sigmoid function from zero, impeding accurate modeling of regulation intensity across various transcription factor concentrations.

These modifications were proposed to impose constraints that are difficult to explicitly add to the system using a graph structure alone. By introducing these changes, we aim to make the model less sensitive to the incompleteness of the gene regulatory network and better integrate hidden information within the data to improve the biological description of the system dynamics. Additionally, these modifications allow us to consider different regulation intensities for each network interaction, which is a more realistic representation than assuming the same value for all interactions.

#### GRN edges and parameter estimation

Within the context of the GRN and cancer, it’s observed that parameter values do more than modulate gene interactions. When certain values approach near-zero levels, they can effectively alter the GRN topology, removing specific network connections altogether. This reflects the potential for cancer to fundamentally reconfigure cellular regulatory landscapes and provides the model with a capacity to indirectly capture GRN topology from scRNA-seq data in the process of parameter estimation.

Dynamic model parameters can be categorized into deterministic and noise-related parameters. These parameters must be able to retrieve expression values compatible with the data and known biological behaviors. Since the expression values obtained from scRNA-seq data result from genetic and epigenetic regulation, using these data for parameter estimation requires implicitly considering all contributions, including those from the microenvironment.

It is important to note that the parameters are estimated for the system at values close to equilibrium. There is no guarantee that they will remain consistent in regions far from equilibrium. States far from equilibrium might exhibit genetic and epigenetic differences, deviating from the behavior predicted by the parameter tuning. Nevertheless, the dispersion of experimental data described by the model may sufficiently reflect the statistical dynamics of GBM for characterizing its subtypes.

We proposed different ways of addressing the challenges associated with estimating the deterministic parameters in our model. These challenges arise from a large number of parameters, the selection of an appropriate estimation method, and the need to ensure biological interpretability. One concern when dealing with a large number of parameters is overfitting; thus, it is important to be aware of this issue and to approach it cautiously.

Additionally, identifiability is a crucial aspect to consider, as it can be difficult to uniquely estimate parameters from the available data in models with many parameters. This challenge may lead to multiple sets of parameter values that produce similar model outputs, making it difficult to draw meaningful conclusions. Furthermore, estimating many parameters often requires significant computational resources and time due to the increasing search space for the parameter values. The complexity of models with many parameters can make them harder to interpret and understand, as the relationships between variables may be obscured.

Keeping these factors in mind, we explored various parameter estimation strategies that strike a balance between model complexity, computational efficiency, and biological interpretability. We considered the following three scenarios: (i) with 2 parameters per equation (one for activation and one for inhibition); (ii) with one parameter related to each input vertex (divided between activation and inhibition), that is, 2*n* parameters per equation (including null values); and (iii) with a combination of (i) and (ii), i.e., *n* × *n* parameters (including null values). Departing from Equation (7), equation (18) illustrates the first case:

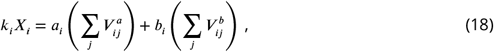

where a possible biological interpretation is an activation and inhibition intensity proportional to the target gene, for example, due to epigenetic regulations. The equation (19) brings the second case,

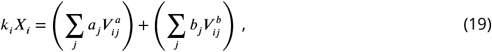

where the consideration of 2*n* parameters would be due to intermediate factors affecting the interactions preceding the binding to the promoter region and the resulting gene transcription. The equation (20) brings the last case,

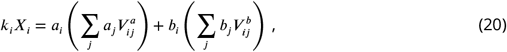

where the idea was (i) to obtain a different parameter for each edge and (ii) to capture as much information as possible. Each activation coefficient would be *a*_*ij*_ = *a*_*i*_*a*_*j*_, with an equivalent procedure for the inhibition coefficients *b*_*ij*_.

To estimate the parameters of cases (i) and (ii), we used the *L*_1_-norm robust regression that can be solved as a linear programming problem (***Inc., 2023a***). We used the Simplex algorithm in the Mathematica environment (***Inc., 2018***). Assuming uniform and constant degradation coefficients for all mRNA molecules, we have for all gene *i, k*_*i*_ = *k*, and equations (18) and (19) can be rewritten in the form of the following equation:

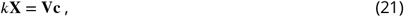

with **V** = (**V**^*a*^ |**V**^*b*^), **c** = (**c**^*a*^ |**c**^*b*^), for (**c**^*a*^)_*i*_ = *a*_*i*_ and (**c**^*b*^)_*i*_ = *b*.

The parameter estimation was conducted by simultaneously incorporating all centroids of all basins of attraction, meaning that the parameters were chosen to capture the contributions of all possible equilibrium states of the system. This approach was taken to avoid overfitting individual clusters, which could potentially hinder the representation of other attractors. Mathematically, for each centroid vector **X**_*α*_, we build the matrices **V**_*α*_ and the vectors ***β***_*α*_ = *k***X**_*α*_, and stack them as

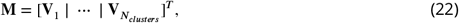

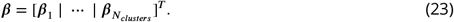

By doing so, we solve the *L*_1_-norm minimization problem

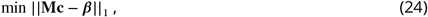

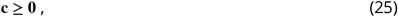

then we compute and store the maximum residual *R*_∞_ = ||**Mc** − ***β***|| _∞_ and the total *R*_1_ = ||**Mc** − ***β*** ||_1_ for each fit as a regression quality measurement.

We obtain the solutions for cases (i) and (ii) for various combinations of *n* and *S*, the regulation functions coefficients. The values of *n* ranged from 1 to 4 in increments of 1, while the proportionality constants *f*_*a*_ and *f*_*b*_ ranged from 0.1 to 1.3 in increments of 0.2. These constants were applied separately for activation and inhibition regulation functions. We used each of the proposed regulation functions for each parameter combination. After estimating the parameters, the absolute residuals list, *R*_∞_, and *R*_1_ were saved. We performed another estimation for each set of parameters obtained using equation (19). This additional estimation allowed us to combine the newly estimated parameters with the initial set, as in equation (20).

#### Noise characterization and stochastic simulation

We subsequently solved equation (3) numerically using the Euler-Maruyama method with an Itô interpretation, where noise is added before increasing expression levels. A Stratonovich interpretation is also possible, as discussed in (***Coomer et al., 2022***). We employed a multiplicative noise function *g*(*X, t*), which accounts for possible mean and standard deviation dependence after a logarithm transformation (***Bioconductor*.*org, 2023***; ***Hafemeister and Satija, 2019***), as shown in equation (26):

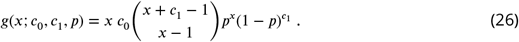

where *c*_0_ is the noise amplitude that scales a negative binomial distribution, with *c*_1_ and *p* empirically determined to fit the data best. *p* is any positive real number less than or equal to 1, and *x* represents the expression level as any positive real number.

For initial conditions such as experimental data points or attractor coordinates, the system is expected to evolve towards the centroid of the cluster it belongs to or maintain trajectories around its average value. To investigate the system dynamics, we performed stochastic simulations with a time interval of 50 a.u. (time steps of Δ*t* = 0.1) to ensure the system reaches equilibrium states. We varied the noise amplitude *c*_0_ (3.5 and 7.0) and explored the impact of different activation and inhibition levels (*a, sa, b*, and *sb*) from 0.6 to 1.4 (0.1 by 0.1). These settings were determined through extensive preliminary simulations to understand the system dynamics better, reproduce the observed variability in the experimental data, and maintain computational feasibility.

We then stored the parameter configurations that exhibited an average of 15 or more genes remaining within two standard deviations from the respective attractors’ values (as identified by clustering and fitted with the parameters to be stable points) as described in equation (27). In essence, given:

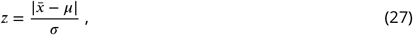

with *μ* representing the cluster centroid of each gene, 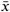 denoting the time average within an interval from time 30 to 50 (allowing the system to fluctuate around the centroid), and *σ* as the standard deviation of the simulated attractor. If *z* ≤ 2 for at least 15 genes out of the 40-gene vector, we stored the parameter set. This approach allowed us to test the hypothesis that the clusters found in the data are statistically significant compared to those obtained by the model with the determined set of parameters. For computing the landscapes, the initial condition was sampled with a uniform spatial distribution. This sampling method was employed on the notion that the trajectories of the system, given sufficient time, would populate regions of the phase space in a manner consistent with their statistical significance.

#### Clusters comparison and simulated landscape

At the culmination of our previous section, we underscored our strategy of analyzing the in silico data akin to the experimental scRNA-seq data. This involved utilizing cluster centroids as a close approximation of fixed points to delineate the attractors and their respective basins. Such an approach not only serves as an analytical framework but also paves the way for the critical validation step: a juxtaposition of in silico and experimental data distributions. To what extent we could reproduce data centroids and heterogeneity considering stochastic fixed point dynamics? Given the necessity of this validation, an essential preliminary step was undertaken—endeavoring to align simulated clusters with experimental ones.

To facilitate the clustering comparison, we rearranged each simulated data cluster to match its closest experimental data cluster. The rearranging starts with (i) computing the centroids of experimental and simulated clustered data. Both cases contain grouped cells linked to each cluster stored in a list of lists manner, with each sub-list representing the cells of an *α* cluster. This way, the centroids for each case are stored in an *α*-ordered list. With two lists of centroids, we then (ii) calculated the Euclidean distances between each element using the experimental centroids as a reference. This step ends up creating a distance matrix. (iii) The reordering starts with an algorithm that identifies the simulated centroid corresponding to the smallest distance to one of the experimental ones. (iv) These indices are mapped together and removed from the distance matrix. This step ensures that each element in the simulated set is associated with a unique experimental cluster. (v) This process is repeated until mapping all experimental centroids to one of the simulated data. (vi) The code returns the reordered simulated clusters in a list of lists format. (vii) For differing numbers of clusters, the ones not matched are appended at the end. This code pseudocode is present below. This process aids in identifying similarities and differences between the two sets and helps to understand the underlying structure of the data.

##### Algorithm 1

Reorder Clusters

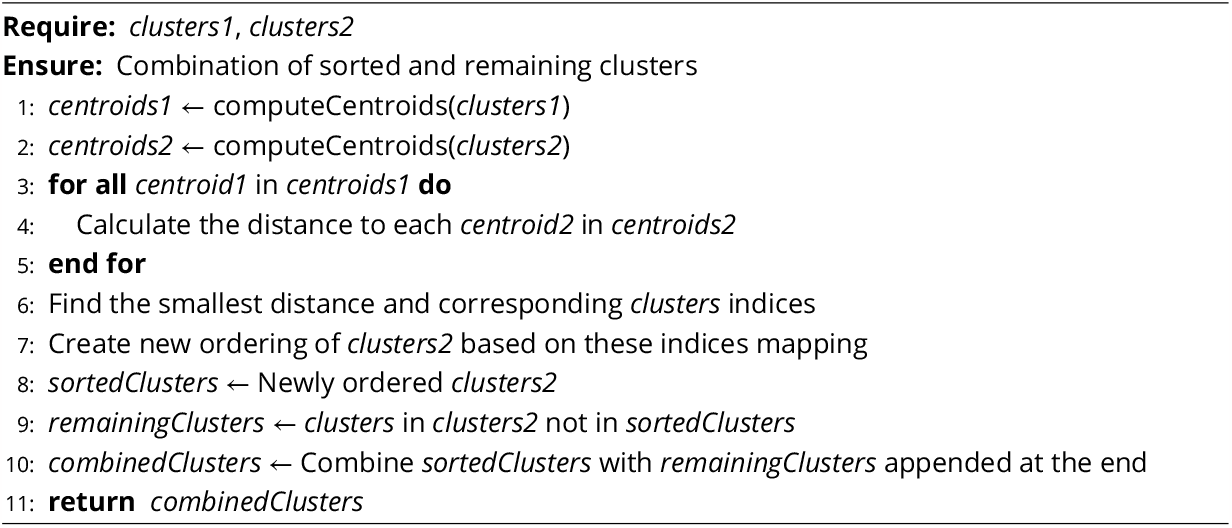

Finally, we use different ways to assess the compatibility between experimental and simulated distributions, including their respective landscapes, computed using equations (9) to (11).

### Dynamics inside basins of attraction

Within the framework of our model, we sought to understand the depth of GBM’s heterogeneity by examining the basins of attraction. Our primary focus was to assess the model’s capability to capture data features, notably the heterogeneity evident in GBM. We aimed to ascertain the stability and scope of the cancer subtype attractors, as these signify the regions significantly influenced by them.

Additionally, we ventured into the statistical dynamics inherent within each attractor. This involved evaluating the consistency between sample and time averages, a vital aspect of the used framework. We employed autocorrelation functions, which facilitated the determination of the system’s inherent timescales and the propensity for transitions between attractors.

### Gene expression dispersion in Cancer

The dispersion observed in cancer data prompts inquiries about the underlying dynamics causing such patterns (***Uthamacumaran, 2021***). In this study, we postulate that the observed dispersion might be influenced by stochasticity, chaotic dynamics, or a combination of both. Regardless of the underlying reason, we assume that these dynamics unfold within specific basins of attraction (see section Basins of attraction: deterministic vs stochastic modeling).

We view gene expression dispersion as a consequence of cancer progression. Factors like genetic mutations, epigenetic shifts, and changing cellular microenvironments amplify the system’s diversity, resulting in more intricate and varied expression distributions. Such diversity mirrors the myriad ways in which cancerous cells can organize and interact, mirroring the increasing complexity and heterogeneity of the tumor environment (***Chen and Wang, 2016***).

Past research has delved into the rising entropy associated with these phenomena (***Nijman, 2020***; ***Tarabichi et al., 2013***). This increase in entropy not only points to varied cellular states but also possibly fluctuating energy levels. Given that differences in energy levels between cell types closely intertwine with genetics and epigenetics, our aim is to examine the stability of cancer subtypes through the lens of these varied states’ attraction basins. We utilize the Waddington landscape concept to study the system’s behavior upon settling into these configurations, as opposed to focusing on the journey leading to them.

Our approach centers on observing the system’s properties over time to ensure a comprehensive capture of its dynamics and fluctuations. Moreover, sampling must span a broad spectrum of potential states. We conceive each basin of attraction as a distinct subsystem, wherein the movement between them is minimal yet possible. Such a perspective aligns with the notion of attractors and the definition of subtypes; frequent transitions between subtypes would undermine their distinct categorization.

Furthermore, rapidly evolving malignancies, like GBM, are expected to explore the phase space more extensively compared to the constrained dynamics of healthy cells or less aggressive tumors. Our analysis, thus, focuses primarily on these basins of attraction, adding rigidity to our suppositions.

#### Fixed Points: Sample vs time averages

In dynamical systems modeling GBM, attractors represent stable cellular states the system naturally gravitates towards. These cancer attractors, identified through cluster centroids, elucidate the system’s tendencies. When analyzing variations around these attractors, the time average and sample average become crucial. Ideally, for a system closely orbiting an attractor, the time average should align with the sample average from various trajectories. This alignment sheds light on GBM’s inherent dynamics and the stability of its states.

To elucidate this phenomenon mathematically, let’s contextualize our variables. The expression level of gene *i* at a particular time point within the interval *T* is denoted by *X*_*i*_(*t*). The spatial domain or basin, in which these expression levels predominantly lie and which is closely associated with the specific cancer attractor ***A***, is termed ℬ (***A***) (refer to section 1 for a comprehensive definition). Within this schema, we propose a model suggesting that the alignment between the time and sample averages becomes increasingly robust as both the temporal subinterval *T*_12_ = *t*_2_−*t*_1_ enlarges and the sample size, *N*, increases. This alignment resonates with our understanding of GBM’s aggressive progression and its manifestation over different timescales. The underlying principle is straightforward: as we gather more data samples and witness increased cellular aggression (indicating shorter times to span the associated basins), our mathematical approximation inches closer to empirical reality. This time average expression for gene *i* can then be expressed as 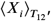, given by:

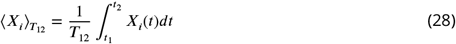

with the time average of *x*_*i*_ in the subinterval [*t*_1_, *t*_2_], and the sample average:

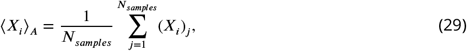

where *N*_*samples*_ is the sample size in the basin of attraction ℬ(***A***), and (*X*_*i*_)_*j*_ represents the *j*-th sample of the gene expression level *X*_*i*_ within the basin of attraction ℬ(***A***).

Our foundational assumption rests on the idea that:

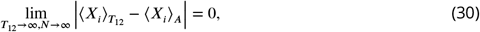

with the length of the subinterval *T*_12_ and the sample size *N* approaching infinity, the difference between the time and space average approaches zero.

#### Timescales and the propensity for transitions

Finally, timescale separation helps assess the relaxation time of *X*_*i*_ within ℬ (***A***) and outside of it. We propose this by examining the decay rates of the autocorrelation function for *X*_*i*_(*t*). The auto-correlation function measures the similarity between a variable and its lagged version, with a faster decay implying a shorter relaxation time. This behavior would suggest a propensity for reaching equilibrium. The autocorrelation function of *X*_*i*_ is given by (***Inc., 2023b***):

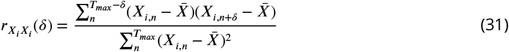

where 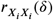 is the autocorrelation at lag *δ, X*_*i,n*_ ≈ *X*_*i*_(*t*_*n*_) is the value of the time series at time step *n* (*t*_*n*_ = *n*Δ*t*, where Δ*t* is the time step interval), *X*_*i,n*+*δ*_ ≈ *X*_*i*_(*t*_*n*_ + *δ*) is the value of the time series at time step *n* + *δ*, 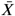 is the mean of the time series, and *T*_*max*_ is the total number time series points. The autocorrelation ratio normalizes the measure to stay between -1 and 1.

The autocorrelation function provides valuable insights into the system dynamic guided by a driving force. Derivatives greater than zero imply a system driven by network interactions. In this case, the autocorrelation values at different lags capture a pattern and may not oscillate around zero. As the lag increases, the autocorrelation values might decrease, but still suggesting the influence of the driving force persisting over time.

On the other hand, the driving force’s influence becomes constant or negligible for a system in a steady state. Being mainly affected by stochastic fluctuations, the autocorrelation at different lags likely oscillates around zero. It indicates little or no correlation between points and more unpredictable behavior, less dependent on past values. We expect to achieve this behavior fast once using the centroids (i.e., attractors) as starting points. In this case, discrepancies would possibly relate to non-stability or transitions.

To proceed with the autocorrelation analysis, we defined a vector 𝒦 with each element representing the autocorrelation at a specific lag *δ*. So 𝒦 can be computed as follows:

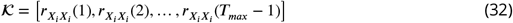

with each value 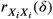 computed using equation (31). By computing the autocorrelation values for increasing lags, we measure if time series values that are further apart are linearly related or correlated.

To quantify the relaxation time, we defined the timescale by:

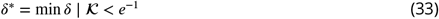

with *δ*^∗^ being the lowest lag *δ*, related to a characteristic timescale, at which the autocorrelation value falls below the threshold *e*^−1^.

In conclusion, these methods offer a strategy to validate the aforementioned criteria, delving into the consistent patterns of gene expression within the simulated dataset. This examination underscores how such assumptions can significantly enhance our understanding of GBM GRN dynamics derived from scRNA-seq data.

## Results

The results section is organized as follows: (i) We present the gene regulatory network, which defines the vertices (genes/TFs) and edges (regulation interactions). (ii) We discuss the initial challenges of choosing a clustering method, which will subsequently influence parameter estimation and optimization. One of the outputs of this optimization process is the selection of regulation functions, which we present in the following. (iii) We compare experimental data with various simulated data obtained using different clustering methods. This comparison serves not only to assess the accuracy of the parameter estimates but also to provide a valuable perspective on the simulation outcomes. (iv) We examine a chosen case from the simulations to verify the dynamics inside the basins of attraction and the hypothesis of equivalence between the sample and time average. The results may also serve as a preliminary study for analyzing experimental data as it becomes more available. Finally, we aim to comprehensively understand the relationships between data analysis, parameter estimation, regulation functions, and simulations by presenting the results in this order.

### Glioblastoma GRN

Our initial step was constructing a GRN, with the methodology outlined in figure S3. This network, visualized in Figure 2, captures the intricate interplay among key genes and markers related to GBM subtypes. Primarily based on an initial list of GBM markers, the MetaCore platform autonomously expanded the network, ensuring an objective, bias-free augmentation. The resulting structure comprises 40 vertices and 242 edges: 187 activations, 11 self-activations, 41 inhibitions, and 3 self-inhibitions. Supplementary Table rede_BT_ALL_seurat.xlsx provides details on the vertices, including those initially selected and those added by MetaCore.

**Figure 2.**
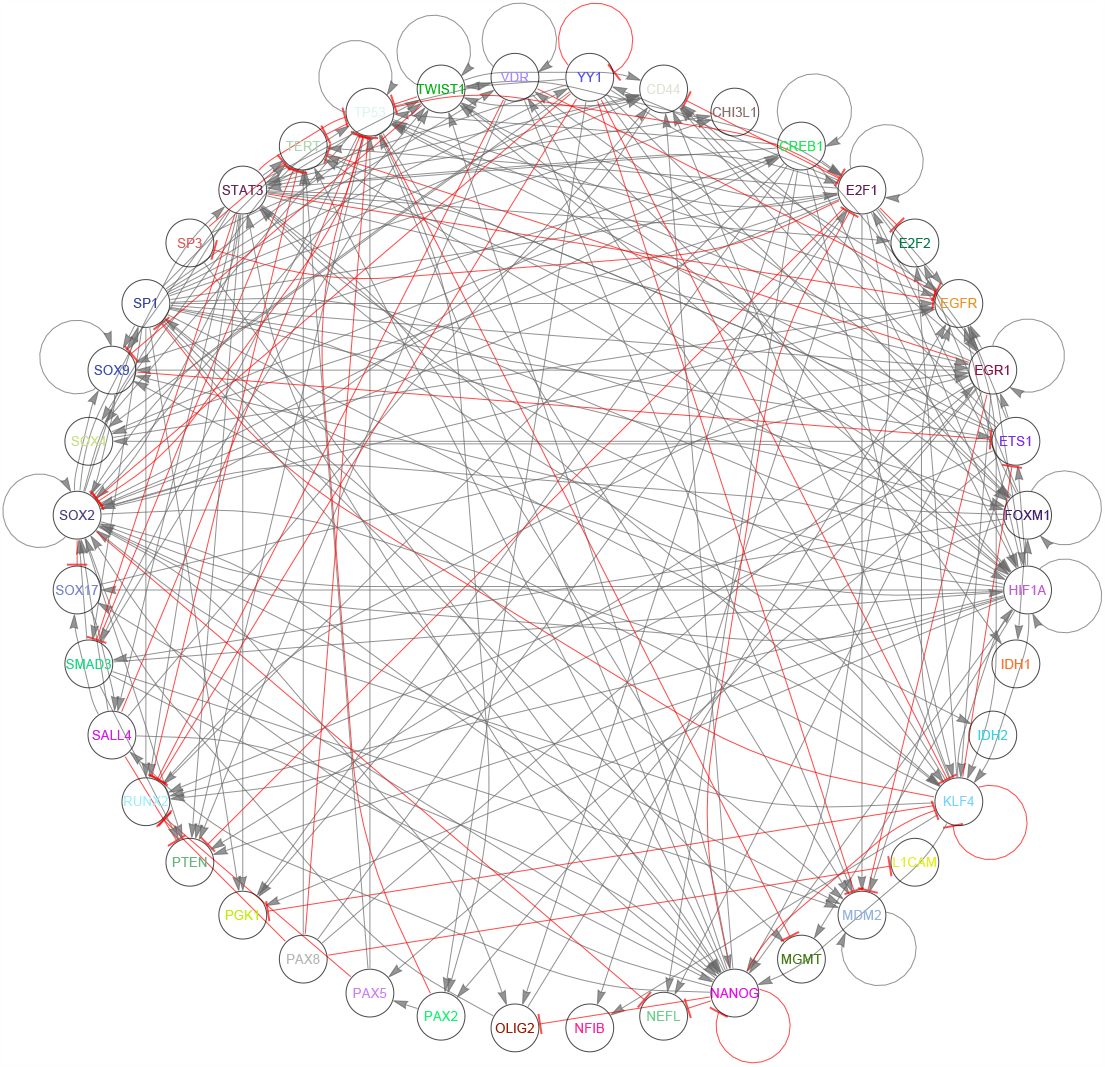
Gene regulatory network for single-cell RNA sequencing of the tumor core of 4 patients with GBM. Black lines with flat arrows represent activations, and red lines with arrowheads represent inhibitions. It contains 40 vertices and 242 edges, with 187 activations, 11 self-activations, 41 inhibitions, and 3 self-inhibitions.

### Clustering methods and parameters estimation

A central question in the scRNA-seq analysis is how to interpret gene expression variations across individual cells. Initial data analysis revealed a group of genes with apparently multimodal distributions (Figure S1). Such patterns, inherent in complex systems like GBM, possibly hint at various cellular states, emphasizing the tumor’s heterogeneous nature. This observation led us to investigate the extent to which this pattern represented the presence of multiple clusters (Figure S2). Focusing on the four pivotal marker genes that delineate GBM subtypes (as described in section Data analysis: Dynamics underlying heterogeneity), our analytical approach employed dimensionality reduction using t-SNE in Wolfram Mathematica. With a perplexity of 60, this method facilitated an optimal visualization of potential clusters, as depicted in figure S4.

To perform the clustering analysis, we utilized two different clustering methods (k-means and NbC). We configured the built-in Mathematica functions for both clustering methods with the ‘PerformanceGoal’ set to quality, the ‘CriterionFunction’ set to standard deviation, and the ‘Distance-Function’ set to Euclidean distance. By comparing the results of these two clustering methods, we aimed to understand the underlying data structure and identify the optimal number of clusters for the given gene expression data. All clusters’ statistics are available in the supplementary table BeforeSim_combinedClusteringResultsNoTidyDataset.xlsx.

Parameter estimation was conducted using the centroid coordinates of all genes within each cluster in an oversized fit, considering *k* = 1. All auxiliary parameter values are available in the supplementary table parAuxData.xlsx. An interval ranging from 0.01 to 10 was used for adjusting the parameters. These values were initially established through manual checks, ensuring that changes in the interval would not lead to substantial variations in the quality estimator values^2^. The first estimate was performed using equation (18), obtaining 784 sets of parameters. The distribution of values for each parameter and both clustering methods can be seen in figure S5. As shown in the figure, some parameters have distributions that vary within the interval limits, such as a[*EGR1*] and a[*SOX4*]. In contrast, others display values constrained to smaller intervals, such as a[*E2F1*] and a[*TP53*]. This pattern is evident in both clustering methods, signifying a strong relationship with the network structure. The larger variations in specific parameters can demonstrate how these parameters are susceptible to changes within the established network. In contrast, others may require considerable compensation, without which it could lead to undesirable changes. The second parameter estimation using equation (19) was performed on top of the first to adjust the values for each interaction. This step’s distribution can be seen in figure S6. The distribution of most parameters is around the unitary value, suggesting a possible correction of the previous estimate. This observation implies that the second estimation step fine-tunes the parameters to enhance the model’s accuracy.

Figures S7 and S8 show the residual for each gene for the two clustering methods superimposed, one for the first parameters estimation and the other for the second parameter estimation. Both figures demonstrate that the two consecutive parameter estimations did not significantly affect the residuals of the first parameter estimation and indicate that it did not result in overfitting. Additionally, the model showed good compatibility for some genes based on the centroids of the experimental data clusters. For instance, genes such as *HIF1A* and *SOX2* may have presented low residuals due to their close centroid values. In contrast, genes like *CD44* and *EGFR* exhibited high residuals and variations due to their distant centroid values. While these residuals may indicate a need for adjustments in network interactions or the model itself, they might also reflect the influence of the clustering method. Therefore, analyzing the relationship between residuals, network structure, and the clustering method is crucial for drawing more accurate conclusions. Furthermore, this analysis could provide insights into the model’s limitations and help identify potential improvement areas. We present all residual values and statistics in the supplementary tables parResidualsTidyData.xlsx and parResidualsStatisticsData.xlsx.

The final step of the parameter estimation process aimed to assess the compatibility between the stability of cluster centroids derived from experimental data and those obtained from parameter adjustments. To accomplish this, we selected the set of parameters with the smallest *T* and *R* values (as displayed in Table 1). Figure S9 (a) and (b) illustrate the activation (left) and inhibition (right) matrices, with the color gradient signifying the logarithm of the parameter values for both clustering methods. The ‘o’ and ‘x’ symbols denote the regulations and self-regulations present in the network, respectively. Parameter values are acquired by multiplying parameters from the first and second fits (columns and rows weighting). The simulation proceeds by multiplying each matrix element by either 0 or 1 (1 if the element corresponds to an ‘x’ or ‘o’ position, and 0 otherwise) to account for only the combinations present in the network. Figure S10 displays the ceiling function (ceil()) of all parameter values for k-means clustering and NbC clustering, respectively. The ceiling function, ceil(), rounds up a given number to the nearest integer. In this case, it helps visualize the order of magnitude of the parameter values. The left images correspond to activation parameters, while the right images depict inhibition parameters. All parameter values are available in the supplementary table parsRegressionData.xlsx.

**Table 1.** Table with optimized parameters. The first and second lines correspond to parameters after k-means and Neighborhood Contraction clustering, respectively. The left side parameters correspond to the optimization using deterministic dynamics, and the superscript 0 and 1 in *R*_∞_ and *R*_1_ inform if it is for the first or second estimation. The right side parameters correspond to the optimization considering stochastic dynamics, with *G*/*C* the mean gene number per cluster when considering the parameters optimization.

| $n$ | $h(x)$ | $f_a$ | $f_b$ | $R_\infty^0$ | $R_\infty^1$ | $R_1^0$ | $R_1^1$ | $c_0$ | $a$ | $sa$ | $b$ | $sb$ | $G/C$ |
| --- | --- | --- | --- | --- | --- | --- | --- | --- | --- | --- | --- | --- | --- |
| 1 | 2 | 0.1 | 1.3 | 1.62 | 1.15 | 24.85 | 20.89 | 3.5 | 1.4 | 0.7 | 0.9 | 1.3 | 20.40 |
| 1 | 2 | 0.1 | 1.1 | 1.72 | 1.69 | 35.43 | 28.48 | 5.6 | 1.4 | 0.6 | 1.3 | 1.2 | 16.71 |

The global optimization process aimed to maximize the number of simulated gene distributions compatible with the experimental data by optimizing the global strength of activation, self-activation, inhibition, and self-inhibition (‘x’ and ‘o’ positions) using multiplicative factors. The factors were chosen to maximize the average number of genes that stay within two sigmas of their cluster centroids. In the first and second parameter estimation steps, 784 sets were generated for both k-means and Neighborhood Contraction clustering methods. The best sets, number 435 for k-means clustering and number 428 for Neighborhood Contraction clustering, were then used in the optimization process. Figure S9 (c) and (d) illustrate the distribution of genes compatible with each attractor during the stochastic parameter analysis for both clustering methods.

For the k-means clustering, the optimization process yielded a list of 191 values, with the best set being number 150 (out of 191). The experimental attractors were defined as A, B, C, D, and E. Varying the regulation weights in each stochastic simulation, attractors A and C had an average of about 15 genes within two sigmas in relation to the observed data, attractor B had an average greater than 20, attractor D had an average close to 20, and attractor E had an average of only 5 genes compatible with the data. Moreover, the following compatibility with data values was observed: attractor A (15), B (24), C (30), D (29), and E (4) when using factors 1.4, 0.7, 0.9, and 1.3, respectively. In the case of Neighborhood Contraction clustering, the optimization process generated a list of 13 values. The best set was number 11 (out of 13), and the clustering ranged from A to G. The compatibility with data values for attractors A to G were 19, 20, 11, 20, 17, 17, and 13 when using factors 1.4, 0.6, 1.3, and 1.2, respectively. Table 1 presents the values of two parameter sets: one for k-means clustering (5 clusters) and another for Neighborhood Contraction (7 clusters).

When comparing the k-means and Neighborhood Contraction clustering methods, we observed differences in the number of genes compatible with each attractor and the quality of the estimated parameters. For k-means clustering, the total number of parameters that lead to genes compatible with attractors A, B, C, D, and E was 191, while 13 with attractors A to G for Neighborhood Contraction clustering. These results suggest that as the number of clusters increases, the set of parameters that matches the data for the established conditions decreases.

### Regulation functions

One of the results of the first and second estimations was the selection of the regulation function. The modified regulation function depends on the transcription factor and the regulated genes. The total number of regulatory functions equals the number of interactions in the network (242 interactions). After the parameters optimization, the best compatibility between experimental and simulated data was obtained with the regulation function of the equation (15). For example, figure S11 shows the activation and inhibition regulation functions for some gene/transcription factor combinations. Each case presents a different relationship between the variables representing the amount of transcription factor mRNA and the activity of the gene promoter. Figures S11 (a) and (b) correspond to the regulation functions of the first clustering choice, while figures S11 (c) and (d) to the second choice. Figure 3 displays some of these cases, such as the activation of *CHI3L1* by *SP1* and the inhibition of *EGFR* by *STAT3*. All surfaces were obtained with *n* = 1 and *h*_2_(*x*), with (a) and (b) using *f*_*a*_ = 0.1 and *f*_*b*_ = 1.3 and (c) and (d) using *f*_*a*_ = 0.1 and *f*_*b*_ = 1.1.

**Figure 3.**
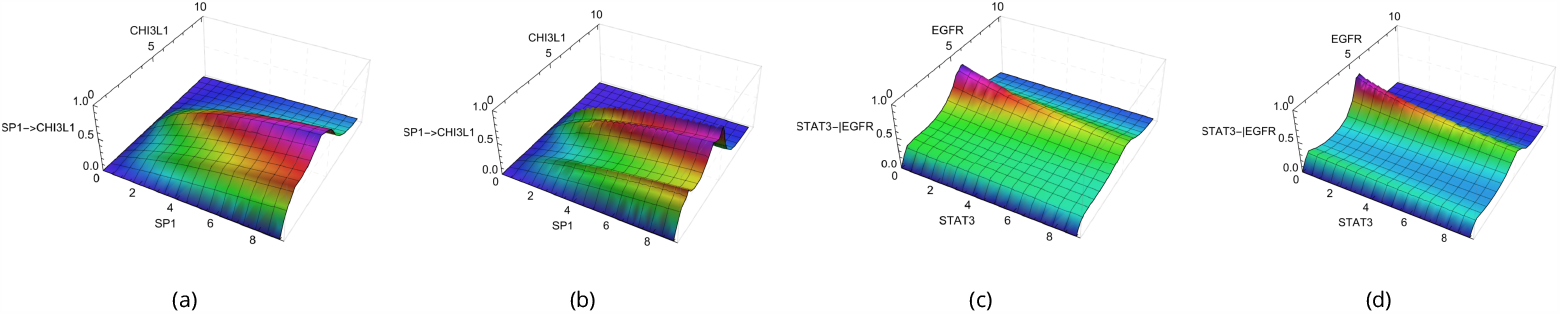
New regulation function *V* with *n* = 1 and *h*_2_(*x*) for different combinations of genes and/or transcription factors. The horizontal axis represents the transcription factor and gene quantification using the normalized amount of single-cell RNA sequencing of experimental data. The vertical axis represents the quantification of the interaction regulations. (a) and (b) Activation values using the k-mean and NbC clusters, respectively (*f*_*a*_ = 0.1); (c) and (d) Inhibitory interactions using the k-mean and NbC clusters, respectively (*f*_*b*_ = 1.3 and *f*_*b*_ = 1.1).

The dependence of the regulation function on the transcription factor can be seen along its axis, as affected by the contribution of the parameter *f* (*f*_*a*_ for activation and *f*_*b*_ for inhibition) and the values 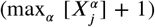. The dependence on the target gene is represented on the axis transverse to the transcription factor axis, where the maximum values for constant transcription factor concentration correspond to the centroid values observed in the experimental data. It is possible to observe how the peaks occur around the average values of *CD44, CHI3L1, EGFR* and *IDH1* for activation functions and *CD44, EGFR, MDM2*, and *PGK1* for inhibition (figure S11).

In constant target gene conditions, regulation functions simplify to one-dimensional curves, similar to standard Hill functions with *n* = 1. At high target levels, these functions approach zero, mirroring trends in our experimental data. This pattern mimics potential biological mechanisms not explicitly detailed in the network, like missing environmental signals or missing interactions. Notably, lower inactivation values mean greater inhibitions; a smaller *V* ^*b*^ value indicates increased inhibition and decreased basal activation. Furthermore, the modified regulation function does not exhibit peaks at zero target concentrations, a deliberate change to prevent unwanted activation peaks that might affect observed null data values. Beyond these specific cases, the system’s progression will be guided by the interplay of network interactions, parameter values, and noise.

### Comparing experimental and simulated data

#### Noise and distributions compatibility

Data transformation often results in a dependence of the standard deviation on the mean (***Bioconductor*.*org, 2023***). Part of this dependency can be reduced depending on the normalization process (***Hafemeister and Satija, 2019***). However, to achieve a better fit between the simulation outcomes and the sequencing data, the function for the multiplicative noise was defined as equation (26), with *p* = 0.23 and *c*_1_ = 8, which were found empirically to fit the data best. Figure S12 presents the graphic for these parameter values. The optimized set of parameters was evaluated by comparing the histograms of expression levels for data at different time points to assess whether the system had reached a steady state. Figure S13 compares experimental and simulated data. Figure S13 (a) shows the simulation outcome for time 50 (500 steps), (b) time 25 (250 steps), (c) time 5 (50 steps), and (d) the initial conditions. The means of the simulated distributions are mostly within two standard deviations of the means of the observed multimodal distributions in the experimental data, demonstrating good compatibility for most genes. However, mainly for higher expression values, the simulated outcomes exhibit smaller standard deviations than those observed experimentally. Gene expression values for each of the three times are available in the supplementary tables AfterSim_Method1_combinedHistogramNoTidy.xlsx and After-Sim_Method2_combinedHistogramNoTidy.xlsx.

#### Clusters compatibility

To evaluate the congruence between patterns in simulated and experimental data, we employed k-means (***Inc., 2023d***) and Neighborhood Contraction (NbC) (***Inc., 2023e***) on the experimental data and extended the analysis to include Gaussian Mixture (***Inc., 2023c***) for the simulated datasets. The clustering outcomes are depicted in Figure S14, which represents data simulated from parameters estimated after the k-means and NbC clustering of experimental samples. Clusters were obtained in two reduced dimensions instead of the original four marker gene dimensions, facilitating a more tractable visual assessment. Preliminary analyses suggest NbC’s superior performance in the reduced dimensional space for both simulation cases.

To further our comparison, we assessed centroids of simulated data juxtaposed with data distributions. Figures S15 and S16 illustrate these for k-means and NbC methods. The centroids, regardless of method, particularly between NbC and Gaussian Mixture, displayed notable similarities. These centroids from the simulated data appeared to more precisely represent the centers of the experimental data’s multi-modal distributions than direct clustering. Such findings could underscore biologically pertinent insights if these distributions reflect core biological activities.

To further elucidate our findings, we analyzed the proportion of data points in each cluster using pie charts from both experimental and simulated data. Figures S17 and S18 illustrate these distributions. Clusters A and B demonstrate consistent proportions across most scenarios, except when using the Gaussian Mixture on NbC simulated data. Other clusters displayed method-specific variations, possibly influenced by metastable clusters. Such clusters can challenge clustering techniques, leading to misclassification or indistinct cluster boundaries. Since clustering methods may address metastable clusters differently, discrepancies arise in both qualitative and quantitative outcomes. Consequently, the chosen clustering method is critical in determining clustering results.

We also assessed the distribution within each cluster. Figures S19 and S20 indicated that standard deviations in every simulated data cluster were consistently lower than in experimental data, suggesting narrow local stability influenced by estimated parameters, and potential noise-induced jumps between attractors. Through boxplots and histograms, we contrasted gene expression distributions within clusters. As evidenced by Figures S21 and S22, the most congruent gene expression distributions were observed in clusters B, C, and E for post-k-means, and A, B, and G for post-NbC. Figures S23 and S24 present the specific distributions, excluding *NEFL* due to its near-zero distribution. These findings emphasize our earlier observations about centroid alignment and standard deviation differences. Notably, parameter estimates using NbC produced notably divergent simulated clustering outcomes, indicating result instability.

In analyzing scatter plots of the combined marker genes, excluding the *NEFL* gene, transitional points between clusters were evident. As depicted in Figures S25 and S26, these points might sustain the narrow stability, but potentially challenge the sample-time average equivalence if they arise frequently. As these points denote distinct cells at the same time step, further exploration of single trajectories over time is essential. Another observation is that, despite the differences in distribution dispersion, a qualitative analysis (low and high) of the clusters would have compatible results. This suggests that the clusters may still be comparable when focusing on their overall trends rather than the specific dispersion of data points.

#### Experimental and simulated landscapes

Considering the results obtained, we now examine the corresponding (epi)genetic landscapes. Assuming that each cluster’s average and deviation characterize each cell state’s distribution, visualizing a potential surface derived directly from the experimental data can help evaluate the clustering quality and serve as a reference for a qualitative comparison with the simulation results. Figures S27 and S28 present the landscapes for both experimental data clustering methods and all simulated data clustering methods. A constant of 0.01 was added due to the presence of null values (or close to zero) for the standard deviation of some genes. This value was sufficient to avoid numerical issues and significantly smaller than non-null values.

Qualitative changes can be observed due to the greater number of attractors in the NbC method compared to the k-means one, particularly in the dimensions *CD44*×*IDH1* (figure 4 (a) and (b)), *CD44*×*NEFL*, and *EGFR*×*IDH1*, with the presence of new basins. When comparing these experimental landscapes to the simulated ones (figure 4 (c) and (d)), as expected from our previous discussion of section, there are significant differences. The landscape is a precise structure that requires a high degree of correspondence between means, standard deviations, weights, and attractors. It is important to note that the experimental landscape does not consider dynamics and takes into account data points that may be in transient or metastable states. Regardless, the previous results are relevant and may indicate avenues for further improvements and studies on the system dynamics.

**Figure 4.**
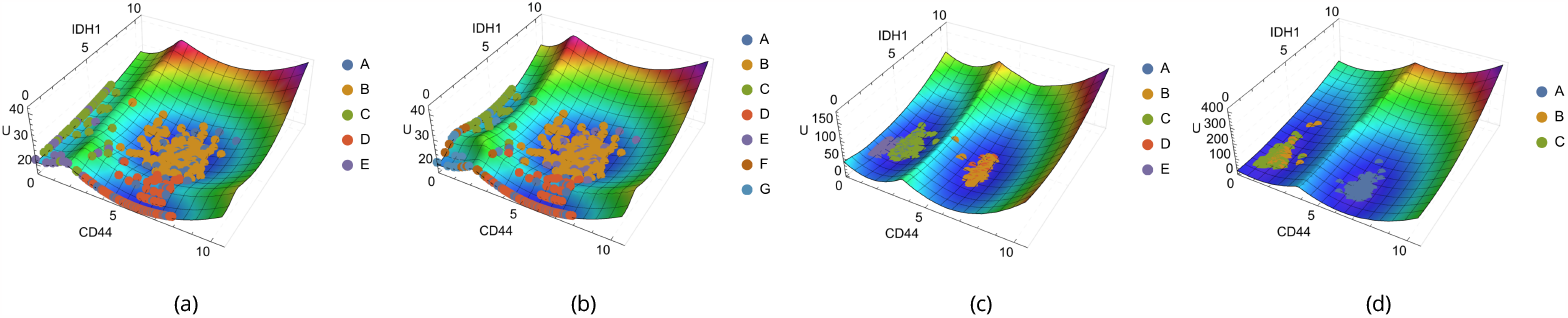
(Epi)genetic landscapes for experimental (a and b) and simulated (c and d) data, with experimental and simulated points overlaid for compatibility visualization. (a) Landscape for experimental data with k-means clusters; (b) Landscape for experimental data with NbC clusters; (c) Landscape for simulated data after parameter estimation with the k-means centroids and clustering with Gaussian Mixture; (d) Landscape for simulated data after parameter estimation with the NbC centroids and clustering with Gaussian Mixture. The horizontal axes show the expression values of each marker gene, while the vertical axis represents the values of the landscape.

### Dynamics inside basins for chosen simulated case

The results and analysis presented above lay the groundwork for further investigations into various topics related to the system’s dynamics and properties. Here, we will explore results concerning the statistical dynamics inside the basins of attraction. First, let’s examine a collection of trajectories originating from the centroids of the clustered simulated data. Then, we selected one case to investigate the dynamic properties of the trajectories. This case corresponds to the Gaussian mixture after the k-means parameters optimization. Figure S29 displays the trajectories starting from each cluster centroid for each gene, using the parameters obtained after k-means clustering and the Gaussian mixture cluster classification.

For most clusters and genes, the trajectories oscillate around their respective centroids. However, for these single trajectories of each cluster, we can observe that specific genes, such as *CD44* and *EGFR*, exhibit transitions (figure 5). Under the hypothesis of sample and time average equivalence, such transitions may not be frequent and may occur on a longer timescale than the component dynamics. This suggests that a more detailed investigation is needed to better understand the nature and frequency of these transitions in the context of the system’s statistical properties.

**Figure 5.**
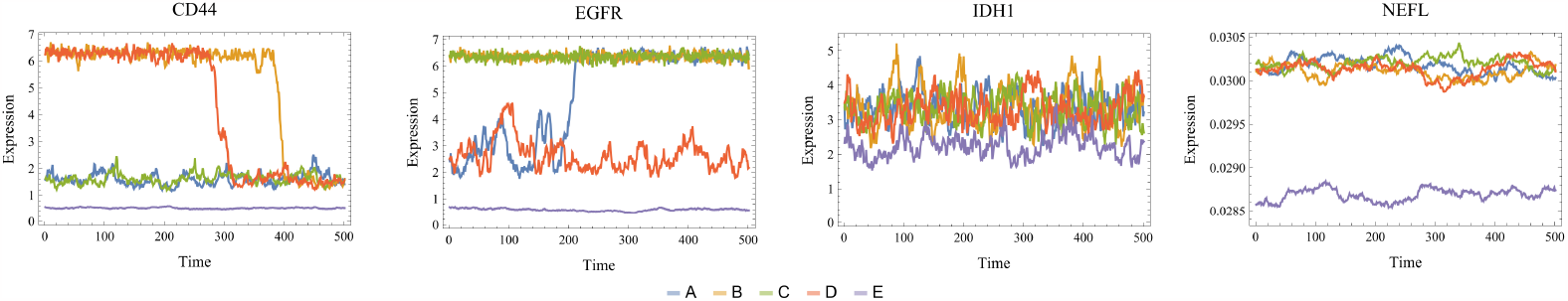
Trajectory plots for each basin (A, B, C, D, and E), showcasing the dynamics of the four marker genes (CD44, EGFR, IDH1, NEFL). The trajectories illustrate the time evolution from initial conditions as the centroid of the clusters found by Gaussian mixture after the parameters found using the k-means centroids. The horizontal axis represents time steps, and the vertical axis shows the expression level of each gene.

To further investigate the transitions between components, we focused on two main features: the identification and quantification of the transitions among the defined basins of attraction and the time spent in each basin before the occurrence of a transition. This analysis was performed by sampling the time steps of each trajectory starting from the centroid of each cluster, measuring each point’s Euclidean distances to all basins, and assigning it to the closest one. We also considered establishing a threshold to determine whether a point belongs to a basin. Still, it would lead to the challenges associated with complex dynamical systems and basins of attraction. In addition, determining the basins’ exact boundaries and neighborhood sizes can be difficult, as the basins’ shapes may be irregular and not easily described by simple measures like distances or standard deviations. Ultimately, we decided to focus on the closest basin assignment, acknowledging the limitations and complexities involved in this approach.

We sampled 100 trajectories for each cluster and tracked their paths. We stored all time series data in a *.m Wolfram format file named AfterSim_trajectories_resultsListForTrajectories.m. Due to the large file size, we wait requesting. Figure S30 (a) and (b) show the quantification of the transitions between the basins. Absent basins indicate that there were no transitions to or from them. Figure S30 (a) displays the number of transitions between each cluster, which may be affected by multiple transitions within a single trajectory. The most frequent transitions were observed between clusters A and C, followed by D to A. Both of these transitions are visible in figure S29. Figure S30 (b) quantifies the probability of a jump from one cluster to another. For example, transitions from cluster A have a probability of 0.85 to go to C and the remaining probability of going to D, and so on.

Studying the time spent by a trajectory within each cluster can help us to understand the system’s stability and the relative significance of each identified state. Long residence times within a cluster suggest a stable state, while shorter times could indicate a transient or metastable state. Figure S31 (a) displays the time spent in each cluster before it jumps to another. The absence of dispersion in clusters C and E suggests that the trajectories starting in these clusters remained there. This observation does not contradict the transition graph; it simply means that when jumping to cluster C and returning to their original cluster, we might consider that the transition was not fully completed, as the system may not have reached the narrow stability of cluster C. This could be due to significant fluctuations in other dimensions. In any case, we can observe that the time spent before transitioning is relatively high for all basins compared to the total observation time.

Additionally, figure S31 (b) shows the histogram for the number of transitions within each trajectory. In other words, considering the 500 trajectories (100 for each cluster), most did not exhibit any transition. The most frequent number of transitions was 1, followed by a decrease in frequency as the number of transitions increased. These results agree with the hypothesis of low-frequency transitions. However, quantifying a timescale within each basin and for basin transitions is still required to verify the hypothesis and further understand the system’s behavior.

Before assessing the timescale, let’s consider figures S32 to S35, illustrating the dispersion of trajectory time points for each basin. We can see the well-defined localization and even the time points possibly representing transitions between clusters. The points are all time points for three trajectories of each cluster. Figure 6 summarizes the results, with (a) one of the 3D trajectories, (b) the transition matrix, (c) the frequency of transitions per trajectory, and (d) the time spent before a transition. The 3D plots help visualize the complexity of each trajectory, and it is important to remember that they are a simplification of the 40-dimensional space. This analysis provides a qualitative intuition, given that the biological system is much more complex.

**Figure 6.**
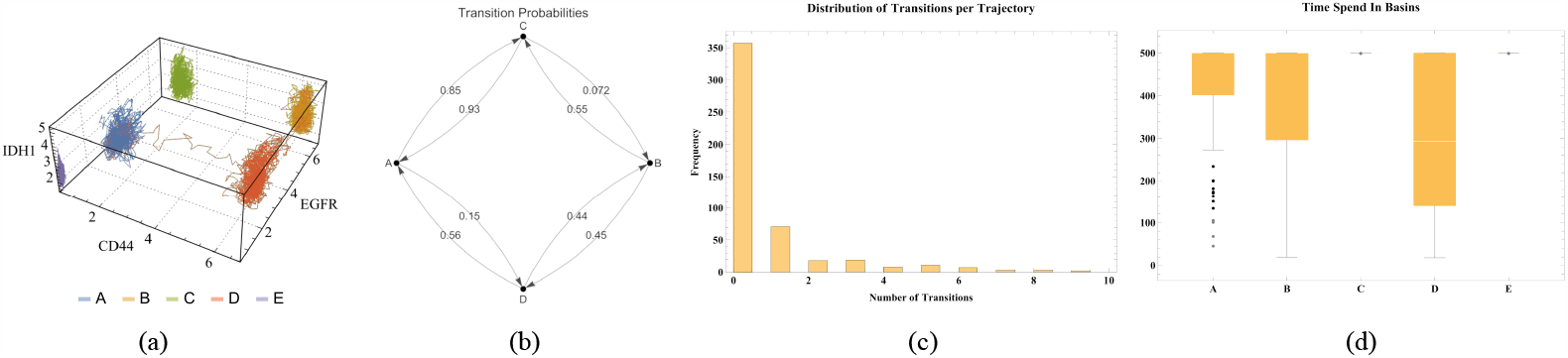
(a) Three-dimensional visualization of full trajectories in three of the markers space. All axes display the expression values of each marker gene. Each line represents an entire time of the three considered trajectories. Each color/letter indicates its respective basin. (b) Transition graph illustrating the connections between different basins. Vertices represent basins, and edge weights represent the probabilities of transitions between basins. (c) Frequency of transitions per trajectory. Histogram showing the frequency of transitions between basins in each trajectory, highlighting that most trajectories do not present any transition, and those that do tend to have a small number of transitions. The vertical axis shows the frequency of each number of transitions per trajectory, while the horizontal axis the number of transitions per trajectory; (d) Analysis of time spent in basins before a transition. Box plots reflect the distribution of time spent in each basin across all trajectories before they present a transition. The vertical axis represents the time spent in the basin, while the horizontal axis the correspondent basins.

Studying autocorrelations within the trajectories is essential to understand better the system’s dynamics and gain insights into the timescales. Autocorrelation analysis can reveal the system’s temporal structure. By examining these structures, we can verify timescales that characterize each basin’s internal dynamics or are associated with transitions between basins. Furthermore, this approach allows us to connect the qualitative intuition provided by the trajectory plots with quantitative measures that can more accurately describe the system’s statistical behavior.

Figure S36 shows the time series side by side with its corresponding autocorrelation functions for all genes and basins considering two sampled trajectories. The autocorrelation functions represent the autocorrelation for each time lag up to a specific maximum lag. The autocorrelation function can be a powerful tool for understanding the characteristic timescales of the system’s dynamics. By analyzing the autocorrelation function, we can identify the timescales at which the system exhibits significant correlations, indicating the persistence of certain behaviors or states. In addition, we can see that each basin and gene may present different timescales, with consistency for both simulations. Besides the visual inspection, we defined another way to quantify this due to the data complexity. We computed the timescale as the minimum time step to reach an autocorrelation value below *e*^−1^ for each basin, variable, and simulation.

Figures S37, S38, and S39 aim to understand the distribution of these timescales within each internal dynamic and search for possible transition behavior. Figure S37 shows that even with timescales varying between genes, they tend to present similar values for each gene. However, compared to other clusters’ timescale patterns, some discrepant cases are observed for *CD44* and *EGFR* genes and for cluster E. We can see in figure S36 (a) that these cases presented transitions, leading to increased timescales. Figure 7 displays an example of the increased timescale for the *CD44* by comparing the autocorrelation with and without the presence of a transition. These figures highlight the different characteristic timescales within the basins and inter-basins.

**Figure 7.**
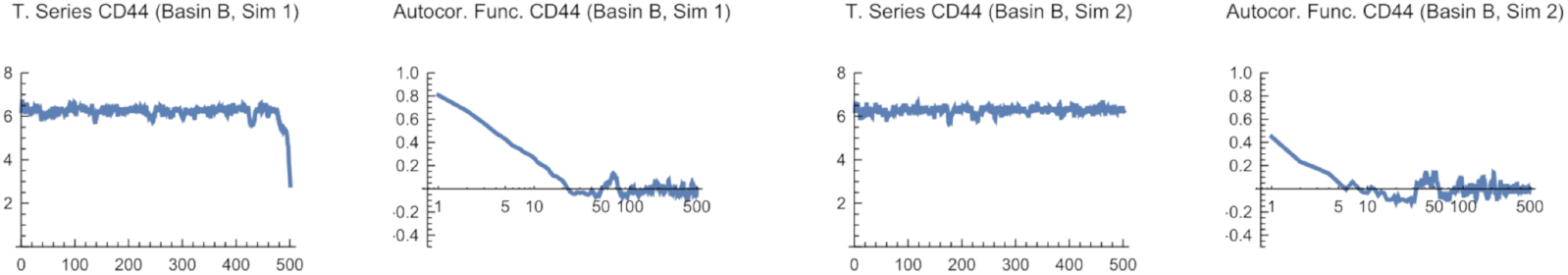
Autocorrelation analysis of time series data for CD44, basin B, and two simulations. Each pair of plots within includes a time series plot (left) with the horizontal axis representing time and the vertical axis representing expression values, and an autocorrelation plot (right) with the horizontal axis representing time lags and the vertical axis representing autocorrelation values.

Furthermore, figures S38 and S39 emphasize the values in each basin. Figure S38 shows the significant variations of timescales for the genes within each basin. Still, given the previous discussion, we note that some genes present very narrow timescale distributions while others have wide error bars. This suggests that the latter may be related to transitions and could be potential variables for further analysis.

Finally, figure S39 shows that the average over all genes and trajectories yields very close results, with only cluster E displaying a different pattern. This may suggest that the timescales are interconnected, and the system may somehow compensate for them. The underlying biological processes at play within the system may lead to interconnectedness that allows for compensation across different timescales. This observation opens up opportunities for further investigation into the mechanisms behind this behavior and how it might relate to the system’s overall function and stability.

After conducting all these analyses, we can finally assess the compatibility between time and sample averages. Figure S40 demonstrates this compatibility for nearly all clusters and genes. The left panels of figure S40 represent the sample average of 100 samples at the final time, while the right panels display the time average, considering 10 trajectories from time 30 to 50 (steps 300 to 500). Once again, the observed discrepancies, such as in *CD44* basin B and D, might be connected to transitions between clusters (figure 8). We computed these averages considering their departure states, and they may have jumped to another basin during the process.

**Figure 8.**
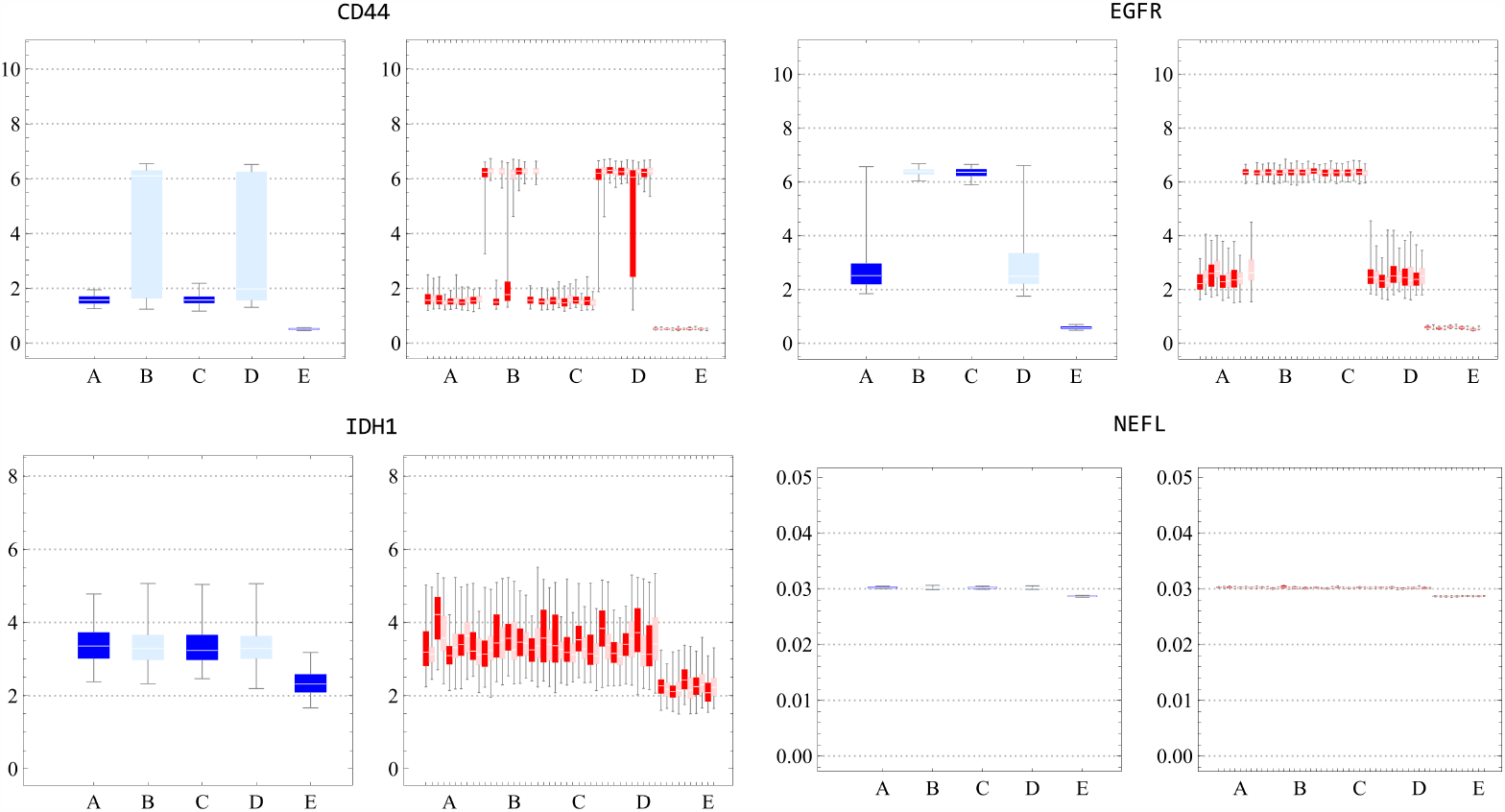
Boxplots for comparison of sample and time averages for all clusters and the four marker genes. The horizontal axis represents the cluster labels, and the vertical axis shows the expression values. The figure displays two sets of plots: the left set shows the sample average of 100 samples at the final time interval, while the right set represents the time average considering 10 trajectories from time 30 to 50 (steps 300 to 500).

In conclusion, these results support our initial assumption of the statistical dynamics behavior inside cancer attractor basins, even if only for simulations. The correspondence with the data suggests that it might also be present in the biological system to some degree. This finding encourages further investigation of these properties within biological samples. Our analyses can be compared and utilized for future applications, ultimately contributing to a better understanding of the system and developing more effective treatments.

## Discussion

In an effort to examine scRNA-seq datasets through stochastic modeling, we encountered challenges ranging from a lack of time series data to complex network construction and a vast number of parameters. Here we discuss our results and interpretation in the context of the present and possible future contributions.

### Dynamics of GBM: aggressiveness and heterogeneity

In this report, we introduced an approach to investigate the dynamics underlying tumor heterogeneity as revealed by scRNA-seq data of GBM cancer cells. We strategically focused on GBM due to its pronounced aggressiveness, which allowed us to make specific assumptions about inherent dynamics. Recognizing GBM’s aggressive nature, we assumed our sample contained a considerable portion of the cancer attractor domain. We proposed that factors like genomic instability, epigenetic changes, and selective pressures could alter inherent trajectories, yielding diverse dynamic behaviors. For instance, stochastic fixed points might become increasingly noisy, or a stochastic limit cycle could either resemble a stochastic fixed point or bifurcate towards it (figure 1).

To the best of our knowledge, this report represents the first application of an integrated framework combining clustering techniques to identify stochastic fixed points and model the cancer attractors dynamics of GBM subtypes using scRNA-seq data. This methodology provided conceptual support for the empirical application of the cancer attractor model. In essence, while our dataset provides a snapshot, this framework provides an underlying theory that shapes this snapshot. This scenario provided a context so we could consider the cluster centroid (sample mean) as being equivalent to the time average of a representative trajectory within the cancer attractor domain — each cluster symbolizing a distinct cancer subtype phenotype.

Additionally, through the examination of a reduced dimension of specific marker genes space across scRNA-seq datasets, highlighted in studies like ***Sáez et al***. (***2022***), we delineated the clusters and identified the centroids. We proposed that these reduced dimensions are particularly amenable to the manifestation of stochastic fixed point dynamics, owing to the inherent characteristics that render them suitable for subtype classification. Our analyses indicated that expression values (both experimental and simulated) are indeed constrained within subregions of these marker gene dimensions, which proved instrumental in studying the dynamics of subtypes. These results reinforced the importance of these subtypes biomarkers. Despite the inability to capture the full breadth of experimental data distributions, our model demonstrates that scRNA-seq data can, to a significant degree, be characterized by our theoretical construct, a premise we plan to expand upon in subsequent sections. We regard this as a progressive step, not only in improving the validity of our hypothesis but also in demonstrating the viability of our modeling approach to elucidate the dynamics underpinning the experimental data.

### GBM GRN dynamics: refining the model and centroid-based parameter tuning

In our modeling approach, we aimed to recreate observed data with stochastic simulations to test the extent to which our hypothesis could account for the data patterns. We initiated our model by establishing a GRN and setting up its dynamics. For our GRN, we focused on marker genes and employed the MetaCore platform for establishing their interconnections. Although this GRN is a simplified model of the complete biological system, we hypothesized that the dynamics of these marker genes would sufficiently encapsulate the subtypes’ dynamics. To better reflect these dynamics, we proposed modifications to the Hill functions, which are often used to represent gene regulation. A key enhancement was the integration of average values from gene expression clusters into the regulatory model. This strategy helped to enclose a range of conditions that are not explicitly included in the model, such as epigenetic differences in patient samples, genetic mutations, missing GRN interactions, or influences from the tumor environment. Our method was designed to provide a representation of the dynamics that aligns more closely with experimental data while also attenuating the network structure’s sensitivity. By adapting the Hill functions, we overcame some of their inherent limitations and successfully captured the centroids observed among different patients. This method proved to be an efficient and effective way to simplify the network’s complexity by incorporating these variations through the data itself.

Building upon the improved Hill functions, linear programming was employed to find the parameters that interpolate the dynamics passing through the centroids in the context of biomarker dimensions. As observed in figure S1, the patients’ data tend to follow similar distribution patterns. Based on this observation, we proposed that a more accurate characterization of the disease landscape could be achieved by taking into account all patient data to improve the sampling around each cancer attractor. Additionally, tuning the model parameters to simultaneously fit the centroids of all clusters was instrumental in preventing the model from being overfitted to any single cluster. It also allowed us to explore whether a single set of parameters could effectively capture intersecting dynamics between all patients and subtypes. In other words, if different subtypes could coexist within the same biological landscape. Indeed, our findings demonstrated that we successfully identified a set of parameters that not only captured the centroids of the data but also delineated a possible dynamic of the disease subtypes. This aligns with findings by ***Neftel et al***. (***2019***), which demonstrate the plasticity of GBM cells and their potential for a single cell to give rise to multiple subtypes. These insights suggest a disease landscape where such cellular states are not only present but also dynamic, with the potential for transitions between subtypes that may contribute to the progression of the disease under certain conditions. Additionally, the selected parameter values leading to these conditions may point to specific, potentially latent configurational states of epigenetic regulation that permit such transitions. In this sense, the identification of a singular parameter set operating as an effective representation of biological phenomena signifies a level of underlying uniformity within the biological dynamics, an attribute that could exist beyond interpatient differences. Lastly, the use of a simplified GRN in our hypothesis does not limit these findings. More complex networks would likely present even more parameter possibilities, of which our model represents a mere subset.

Building upon these deterministic parameters, stochastic features were introduced aiming to match the centroids and distributions observed in the experimental dataset. The challenge lies in the precise calibration of this noise. We sought a noise that neither dominates the dynamics nor is negligible. By achieving this, we assert that the noise model, albeit simplistic, serves as an effective representation. Recognizing the complex factors contributing to this noise, ranging from intrinsic cellular mechanisms to varied patient-specific influences, an effective noise encapsulation arises as a pragmatic choice. Furthermore, the study’s aim wasn’t to trace individual cellular trajectories but to discern broader statistical patterns that emerge from the collective trajectories. Since we are trying to observe the ‘subtypes envelope’ a prior noise was used to have a first idea of results before its improvement. This iterative process allows for a progressive refinement of the noise model in future investigations.

### Parameters variability: heterogeneity and genomic instability

As discussed, we have expanded upon traditional cell-type clustering by incorporating dynamic profiling, which highlights the transient nature of cellular states. Our results accentuated the importance of biomarkers, which proved instrumental in delineating stable states and enabled a more subtle understanding of subtype dynamics. But, while our model has successfully replicated the centroids of these classifications, it did not achieve the same level of accuracy for the dispersions. This section focuses on the challenges faced due to the variability of parameters and the interpretation of the underlying causes, including heterogeneity and genomic instability. Also, their impact on modeling heterogeneity in cancer, including genomic instability. These features, intimately associated with cancer, are directly reflected in scRNA-seq data. They can be seen incorporated in the gene expression variability, both inter and intra-clusters.

These mechanisms introduce variability that makes difficult dynamical modeling, messing with the parameters and hyperparameters in a way that its unique determination becomes difficult. As pointed out in (***Coomer et al., 2022***; ***Weinreb et al., 2018***), multiple parameter combinations could yield results closely mirroring those observed experimentally. To address this multiplicity, we adopted a statistical criteria to analyze and select the most meaningful parameters. Additionally, by changing the hyperparameters we constructed an extensive parameter analysis that gave us insights into the distribution of these parameters. In this way, the heterogeneity both complicates and helps parameter estimation. Complicates in the sense that the complete variability probably does not reflect a single parameter set, but helps in a way that it might end up defining the region of the gene expression space that could contain the centroid of the cancer attractor.

By using different hyperparameters to find the best regulation parameters, we obtained parameter distributions for each gene. These distributions may be more than just a reflection of a methodological change, telling about intrinsic biological characteristics associated with the genes and their GRN topology. Genes showing wide variability in parameter values across different hyperparameter settings might indicate that the behavior of these genes is highly sensitive to changes in their regulatory environment. Multiple sets of parameters that lead to a driven force near zero reflect the nature of heterogeneity, showing that different cells would display different but near expression values but still could lie around the same attractor. Genes with parameter values that remain relatively consistent across different hyperparameters might be considered more robust or stable in their behavior. This robustness might be due to GRN’s built-in compensatory mechanisms. In the genomic context, it suggests that these gene regulatory networks have evolved to maintain their function despite external perturbations. When such a gene does mutate in cancer, the mutation might have profound effects, given that the gene’s behavior is typically so consistent. In these cases, deviations from this narrow window can destabilize the system, potentially leading the cell into aberrant behaviors. This fact might be intimately related to genomic instability.

Additionally, such genes, and the GRN regions they are part of, might be less resilient to perturbations and more susceptible to disruptions. This highlights where certain genes can act as points of vulnerability within the network, predisposing the system to disequilibrium and chaotic behaviors when they are altered. This could be an important feature of this modeling approach, revealing potential genes and regulatory mechanisms to study, new diagnostic pathways, and targeted therapies. But this fact still needs further verifications, increasing sampling size and variability, using a validated network, and comparing the results with biological experiments.

Our parameter modeling offers a robust enhancement to biological interpretation, addressing the limitations of prior studies that often rely on arbitrary parameters values (***Ferrell, 2012***; ***Wang et al., 2011***; ***Li and Wang, 2013b***). Further, we demonstrated that residuals, traditionally utilized as indicators for estimation quality, hold potential as measures of network and model goodness. This proposition needs comprehensive exploration in subsequent research efforts. Notably, the presence of non-zero residuals was anticipated, asserting that an exact equilibrium characterization would necessitate the incorporation of multiple complex elements, rendering a precise depiction implausible.

### Landscape and dynamics inside basins

Central to our investigation is the complexity involved in modeling dispersions, the varied number of basins, and the emergence of distinct phenotypes. An in-depth comparison of experimental and simulated data enabled us to estimate our model’s performance and its potential predictive capability. The observed deviation from expected dispersion of experimental data could underscore the need to explore multiple parameter sets following the stochastic fixed-point dynamics. Moreover, technical noise in experimental data — unrelated to core biological processes — might have masked genuine dispersion that our simulated landscape struggled to reproduce. However, an important feature is that both the experimental and simulated landscapes identified multiple attractors, reinforcing the hypothesis of multiple stable states. Another point to note is that the GRN and parameter values might undergo changes due to factors such as mutations and epigenetic modifications. In that case, it would still support the investigation of short term dynamics characterization and introduce a possible avenue for investigating tumor progression through longitudinal analysis.

In the final stage of the investigation, we chose suitable parameters and proceeded to investigate the in silico dynamics inside each basin aiming to gain new insights by induction from simulated data. This analysis involved examining the stability of each cluster, comparing time and sample averages, and evaluating the time spent within each cluster, which can serve as a measure of stability. By investigating transitions between attractors, we aimed to understand their stability and the interplay between sample averages and time averages, especially since frequent transitions could challenge the equivalence of these two metrics. To ascertain this, we assessed the frequency of transitions and the timescales within each attractor to better understand the system’s dynamics. It also helped to verify the extent to which the stochastic fixed point could achieve sufficient stability to accommodate the wide experimental data distributions.

In conclusion, we stress the value of studying these aspects in more detail by identifying the factors influencing the system’s properties. If cancer aggressiveness indeed causes a shift leading to a statistical dynamics regime as hypothesized, investigating the detailed causality behind this could provide critical insights into cancer progression and potential interventions. To the best of our knowledge, the in silico verification of stochastic fixed point dynamics and transitions between GBM attractors using this integrated approach has not yet been reported.

## Conclusion

Our study introduces a novel approach to the analysis of scRNA-seq data in the context of GBM dynamics. While the dynamic systems perspective on scRNA-seq data is not new, our method of using cluster centroids as fixed point coordinates for parameter estimation is distinctive. Building on the stochastic fixed point concept, we understand that the aggressive nature of GBM not only creates a noisy environment but also profoundly influences the dynamics by determining how extensively the cancer attractor is sampled. In this context, the sample average (used to identify the cluster centroid) is equivalent to the time average of a representative sample traversing the cancer attractor for an extended duration.

Through this methodology, we have established a direct bridge between theoretical dynamical systems and experimental data. Our findings demonstrate a high degree of compatibility between centroids of experimental and simulated data. Our analysis also delves into the stability of cellular trajectories and transitions between attractors. While our study has limitations, this exploration offers deeper insights into the dynamics of GBM subtypes. The transitions between the basins revealed a possible interplay between subtypes, potentially uncovering factors that drive cancer recurrence and progression.

By connecting molecular mechanisms related to cancer heterogeneity with statistical properties of gene expression dynamics, we expect to set the stage for the development of potential diagnostic tools and pave the way for personalized therapeutic interventions. Distinguishing these unique molecular imprints would allow us to forge stronger connections between the ‘geometry of heterogeneity’ seen in single-cell data and distinct instability mechanisms, offering a more enriched understanding of tumor biology. Also, future research can further explore the dynamics within the (epi)genetic landscape of GBM subtypes to understand better the pathways that lead the system before the differentiation of each subtype. Such studies could provide even more powerful insights into subtype development and contribute to developing targeted therapies for GBM patients. Additionally, this approach could be used to assess and monitor the evolution of malignant states, serving as an instrument for diagnosis and treatment.

## Supporting information

Supplementary Table rede_GenesList.xlsx

Supplementary Table rede_BT_ALL_seurat.xlsx

Supplementary table BeforeSim_combinedClusteringResultsNoTidyDataset.xlsx

Supplementary table parAuxData.xlsx

Supplementary table parResidualsTidyData.xlsx

Supplementary table parResidualsStatisticsData.xlsx

Supplementary table parsRegressionData.xlsx

Supplementary table AfterSim_Method1_combinedHistogramNoTidy.xlsx

Supplementary table AfterSim_Method2_combinedHistogramNoTidy.xlsx

## Author contributions

MGVJ designed the analysis, developed the codes, analyzed/interpreted the data, and wrote the manuscript. AMAC revised the mathematical model and its implementation. NC and FRGC ensured biological accuracy. FABS provided structural critiques and improvements. All authors participated in text revision and approved the final manuscript.

## Acknowledgments

I would like to thank Mariana Boroni for reading the manuscript and for her suggestions concerning patient-specific analyses in this report.

## Appendix 1 Basins of attraction: deterministic vs stochastic modeling

Gene regulation can be modeled as a dynamical system, represented by a set of equations 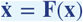, with **x** ∈ ℝ^*n*^ characterizing the state vector and **F**(**x**) a vector field defining the dynamics within the gene regulatory network (GRN) model (***Wang et al., 2010***). We consider an attractor, denoted as ***A***, as a set of points in the phase space that the system converges to under a specific set of initial conditions (***Perko, 2001***; ***Wiggins, 2003***; ***Strogatz, 2018***; ***Guckenheimer and Holmes, 1983***; ***Alligood et al., 2000***; ***Pitzer et al., 2010***).

The basin of attraction, ℬ (***A***), is associated with the attractor ***A*** and is defined as the set of points **x**_0_ ∈ ℝ^*n*^ such that the trajectory of the system starting from **x**_0_ converges to ***A*** as time goes to infinity (***Perko, 2001***; ***Wiggins, 2003***; ***Strogatz, 2018***; ***Guckenheimer and Holmes, 1983***; ***Alligood et al., 2000***; ***Pitzer et al., 2010***). In this context, a basin of attraction represents a set of points in the phase space that lead to similar long-term behavior. We considered that each basin of attraction relates to a group of genes associated with specific conditions, such as cancer or cancer subtypes. A more formal definition could be:

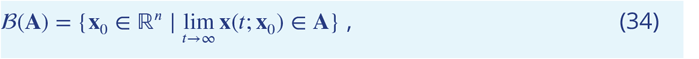

where **x**(*t*; **x**_0_) denotes the trajectory starting from the initial condition **x**_0_ at time *t* = 0.

In the outlined deterministic framework, trajectories converge to attractors given a set of initial conditions. However, biological systems are subject to fluctuations due to inherent stochasticity in gene expression. This randomness leads expression values to be described by probability distributions (***Chen and Wang, 2016***). The temporal evolution of these probability distributions *p*(**x**, *t*) is described by the Fokker-Planck equation (***Kampen, 2007***):

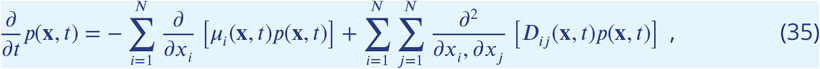

where the temporal change of the probability distribution *p*(**x**, *t*) is determined by two main components: the drift term, represented by ***μ*** = (*μ*_1_, …, *μ*_*N*_), and the diffusion term, associated with the diffusion tensor 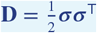. The diffusion tensor **D** is defined as follows:

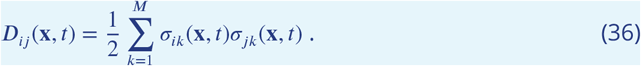

where *D*_*ij*_ (**x**, *t*) is the diffusion tensors’ elements.

In summary, the Fokker-Planck equation (35) describes the evolution of probability distributions subjected to deterministic and stochastic dynamics. It models gene expression fluctuation’s influence on biological systems over time. Studies have shown that in the low noise regime, the solution is a Gaussian and that the first-order moment corresponds to the deterministic model (***Bover, 1978***). Therefore, one alternative is to characterize its first and second moments relationships and solve the corresponding system of differential equations.

To analyze the long-term behavior of gene expression levels in GBM GRN dynamics within the context of our stochastic system, we recognized the inherent assumption that individual trajectories converge to the same behavior when averaged over time. Given the possible complex ways that noise could shape the basin of attraction size/geometry, we introduce a custom distance function *d*(**x, *A***) (***Perko, 2001***) to adjust its definition. It allows for being generic and still captures the complexity of the problem. It becomes:

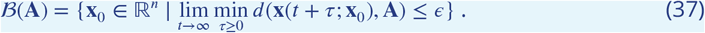

where **x**(*t*; **x**_0_) denotes the trajectory of the system starting from the initial condition **x**_0_ at time *t* = 0, *ϵ* > 0 is a small positive number representing the tolerance level within which the system oscillates around the attractor ***A***, and *τ* ≥ 0 is a time shift that accounts for the oscillation around the attractor. This way, instead of converging to a fixed point, the noise drives the system to remain within the neighborhood of the attractor.

In practice, the choice of the distance function and the tolerance level might depend on the characteristics of the system and the specific dynamics around the attractor. For simplicity, we assumed symmetrical geometries with Euclidean distances and a tolerance level *ϵ* proportional to the distribution standard deviation. Given our Gaussian distribution assumption, we expect it should deal with in silico model fluctuations.

Analyzing the stability of basins and their inter-transitions, we utilized clustering algorithms and inherent assumptions about system behavior over extended periods. By doing so, we sought to understand the possible relations between GRN dynamics and expression patterns observed in experimental data. Essentially, for such systems, it implies that its time average over a long period is equivalent to its ensemble average over all possible system states. Considering the quantification of gene expression for gene *i* at time *t* as *x*_*i*_(*t*), the time average ⟨*x*_*i*_⟩_*t*_ is given by (***Gardiner, 2004***):

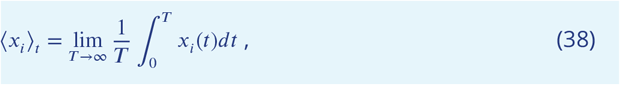

where *T* is the time interval over which *x*_*i*_(*t*) is observed. The ensemble average ⟨*x*_*i*_ ⟩ _*ss*_ for the gene *i* of the gene expression quantification *x*_*i*_ over the stationary state, given the stationary probability distribution *P* (*x*_*i*_), can be calculated as (***Gardiner, 2004***):

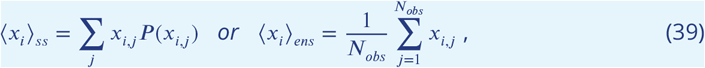

where *x*_*i,j*_ represents the expression quantification of gene *i* in the *j*-th realization of the system, *N*_*obs*_ is the total number of realizations or observations, and the summation is taken over all possible quantification levels of gene *i*. The equivalence of the ensemble average with the time average is given by ⟨*x*_*i*_⟩_*ss*_ = ⟨*x*_*i*_⟩_*t*_.

## Appendix 2

### Glioblastoma Multiforme

Among the different types of cancer, Glioblastoma Multiforme is the most common and aggressive type of brain tumor. It has an average survival of around 15 months with only approximately 10% chances of 5-year overall survival and peculiarities due to its anatomical location and physiological characteristics. In general, without presenting symptoms before the malignant state, it has its highest incidence in individuals of more advanced ages, with approximately 50% cases related to patients over 65 years old.

GBM can occur as both a primary tumor and a secondary tumor. Primary tumors account for more than 90% of GBM cases, having a worse prognosis. Secondary tumors represent approximately 5%, rarer in younger individuals (***Sasmita et al., 2017***). Both the primary and the secondary tumors have different characteristics. Primary tumors present greater expression of the epidermal growth factor receptor (*EGFR*) and mutations in the homologous phosphatase to tensin (*PTEN*) gene. In contrast, secondary tumors show mutations in the tumor suppressor gene *TP53*, not observed in primary tumors (***Sasmita et al., 2017***). GBM can still divide into subtypes according to various criteria and markers, which is challenging. However, it is essential in the development of treatments. The main classification proposals are from Verhaak, Philips, and Jiao. Verhaak et al. proposed four subtypes of GBM: Classical, Mesenchymal, Neural, and Proneural. Philips included a proliferative, marker-enriched subtype of NSCs. Jiao focused on differences in mutations in Isocitrate Dehydrogenase (*IDH*) (***Sasmita et al., 2017***; ***Jiao et al., 2012***; ***Phillips et al., 2006***; ***Verhaak et al., 2010***).

Verhaak et al., in addition to the classification of subtypes, observed similarities in the markers of each subtype and different cell types. For example, the Classic subtype is closer to neurons and astrocytes; the Neural presents markers typical of neurons; the Mesenchymal with both mesenchymal and astrocyte markers, and the Proneural with high expression of oligodendrocyte genes (***Verhaak et al., 2010***). In general, the Classic subtype has high levels of expression for *EGFR*, the Mesenchymal has low levels of the *NF1* gene (Neurophymatosis type 1) and high *MET*, the Proneural subtype features altered Receptor Type Alpha for Platelet-Derived Growth Factor (*PDGFRA*) and mutations in *IDH1*, and the Neural subtype exhibiting mutations in *TP53*, high expressions of *EGFR*, and deletions in inhibitor cyclin-dependent kinase 2A (*CDKN2A*). Additionally, they found a relationship between the subventricular zone (SVZ) proximity and subtypes characteristics: the Neural and Proneural types are closer to the SVZ, displaying faster progression, lower survival rates, and similarity with NSCs, whereas the Classic and Mesenchymal types are more distant, showing a diffuse aggregation pattern and a better response to more aggressive treatments. This association implies that the proximity to the SVZ is a prognostic factor related to pathogenesis (***Sasmita et al., 2017***).

Standard treatment consists of surgical approaches, radiotherapy, and chemotherapy treatments to minimize damage and avoid severe consequences to the nervous system. However, as a complicating factor, the blood-brain barrier still considerably limits the chemotherapy options, which requests for studies capable of proposing viable alternatives (***Gallego, 2015***).

## Supplementary results

**Figure S1:**
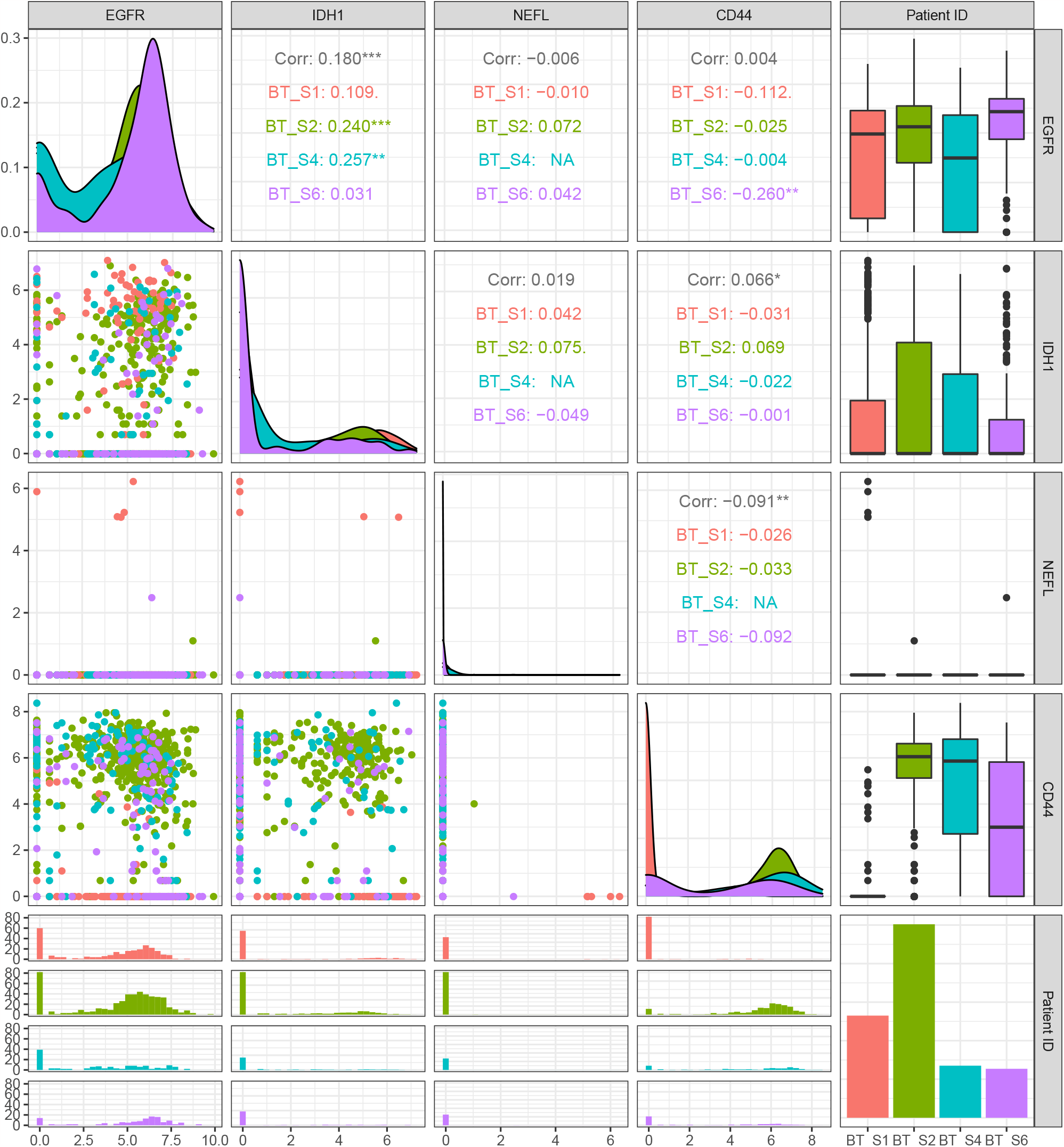
The figure presents a multi-faceted exploration of the relationship between four selected genes—EGFR, IDH1, NEFL, and CD44—for the present scRNA-seq dataset. Each off-diagonal scatter plot showcases the pairwise relationship between two different genes, enabling an assessment of gene-gene relations. These scatter plots are color-coded based on ‘Patient ID,’ providing an additional layer of information concerning inter-patient variability. The diagonal histograms present the distributions for each gene, highlighting their individual expression profiles. Above diagonal histograms, the Pearson correlation coefficients are displayed, offering a quantified measure of the linear relationship between each pair of genes. This comprehensive view allows for a multi-dimensional understanding of gene relations and their distributions across different patients. The arrangement of markers in the scatter plots underscores a notable consistency in the spatial localization of these gene expression markers, irrespective of patient-to-patient variability. In the far-right column, boxplots for each gene across different patients provide an overview of the gene expression variability and central tendency within the patient cohort. The last row of this column features a bar chart that aggregates this data.

**Figure S2:**
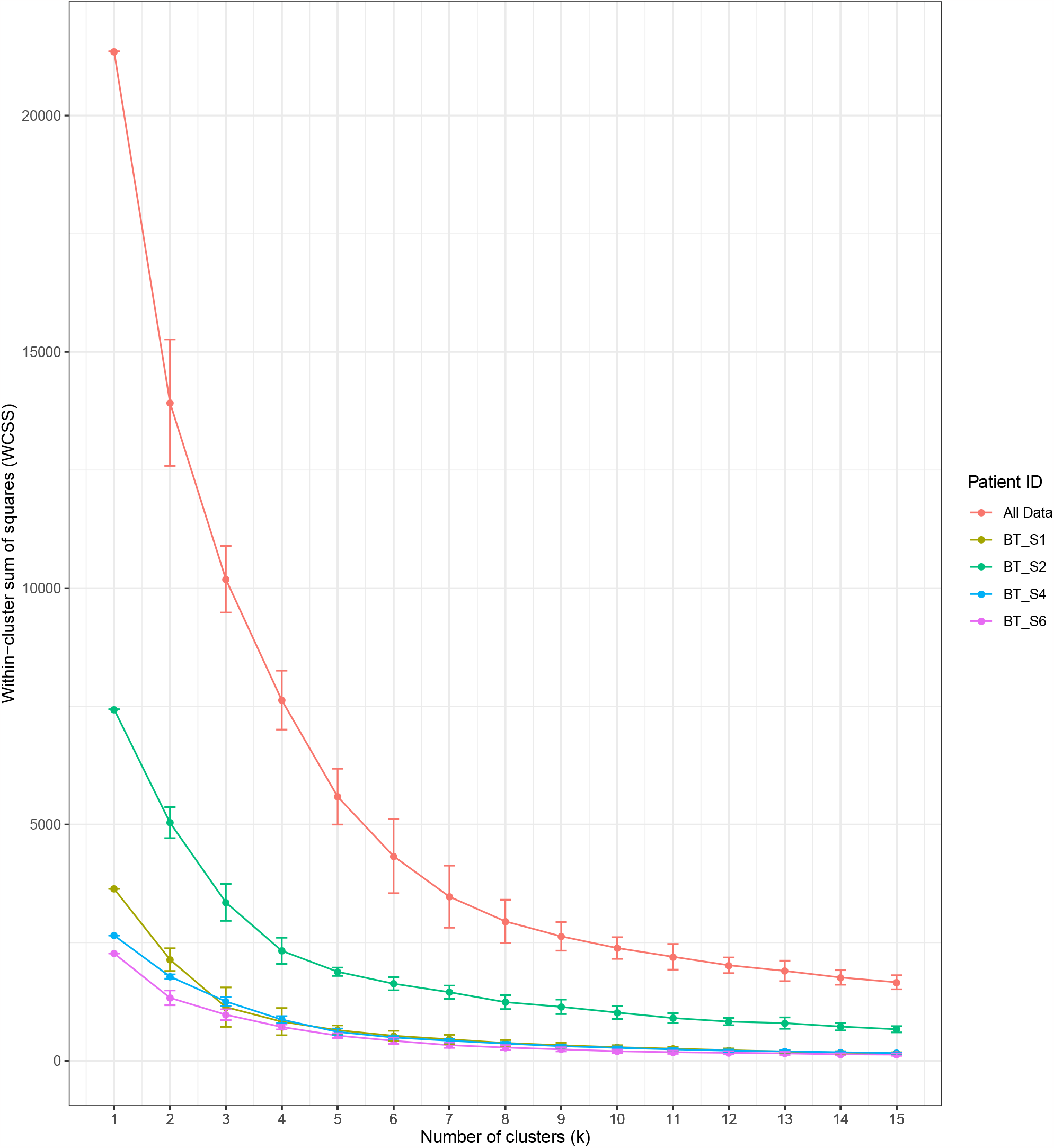
The figure presents an assessment of the Within-Cluster Sum of Squares (WCSS) to determine the optimal number of clusters (k) for the gene expression data, specifically focusing on four key genes—EGFR, IDH1, NEFL, and CD44. It encapsulates results derived from a repeated (100 runs) K-means clustering algorithm, both for individual patients and the aggregated dataset. Each data point represents the mean WCSS value computed across the 100 runs for each k value ranging from 1 to 15. Lines connect these mean points, providing a visual trajectory of WCSS as a function of the number of clusters. The error bars, extending from each mean point, represent the standard deviation of the WCSS values, thereby offering a glimpse into the algorithmic stability across different runs. This approach leverages both a global analysis (aggregated data) and local examinations (patient-specific data) to furnish a robust statistical landscape of the clustering behavior. The plot is color-coded based on ‘Patient ID’.

**Figure S3:**
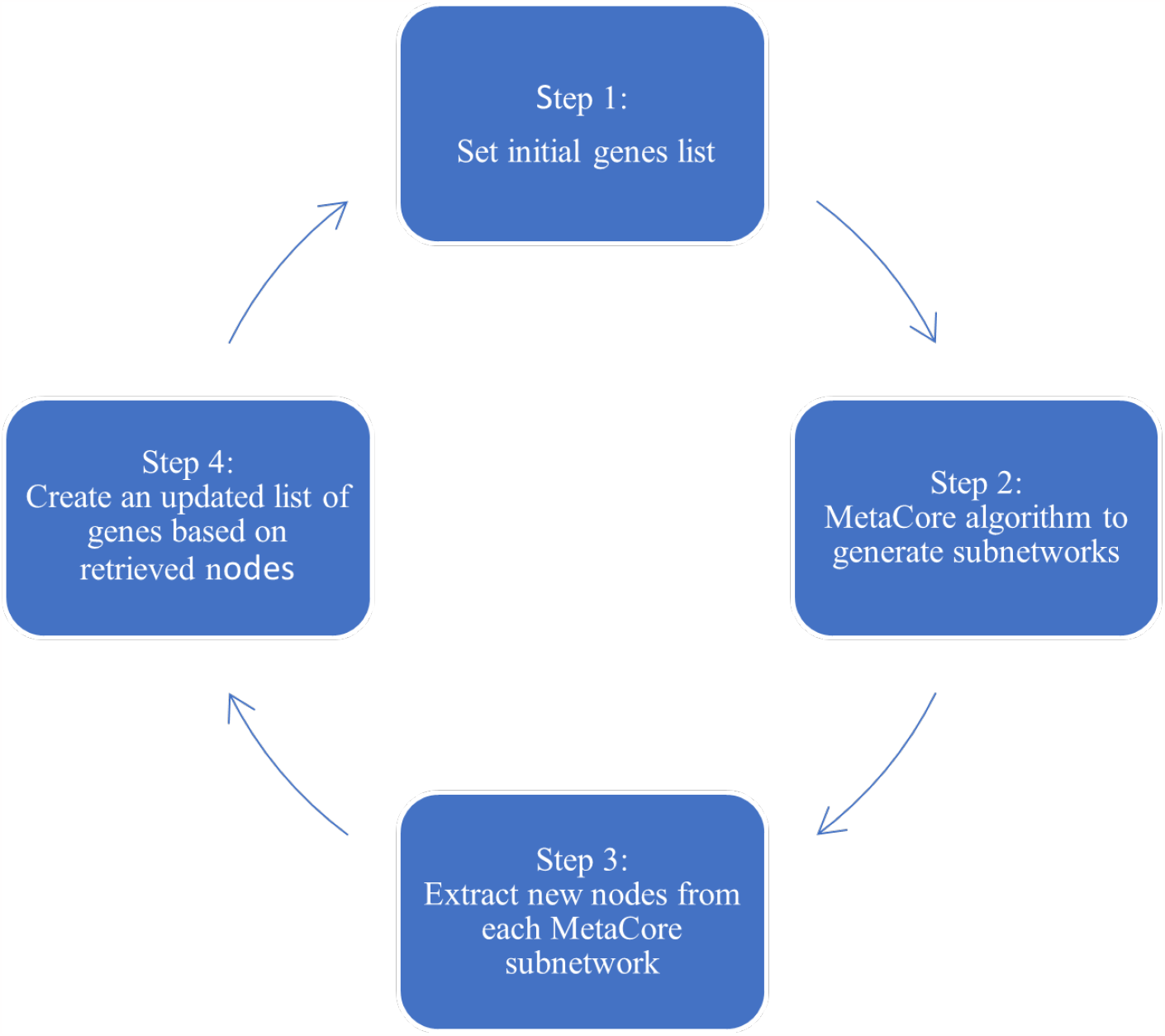
GRN building procedure: (i) Select a group of initial (marker) genes. (ii) Input the list into the MetaCore GRN generator. (iii) Retrieve additional vertices. (iv) Create a new list with the initial vertices and those added by the MetaCore algorithm. (v) Use the updated list as the initial group of genes. This cycle can be repeated until the network is interconnected or until no new vertices are added. The process was performed twice in this case, but the number of iterations can be adjusted according to specific requirements.

**Figure S4:**
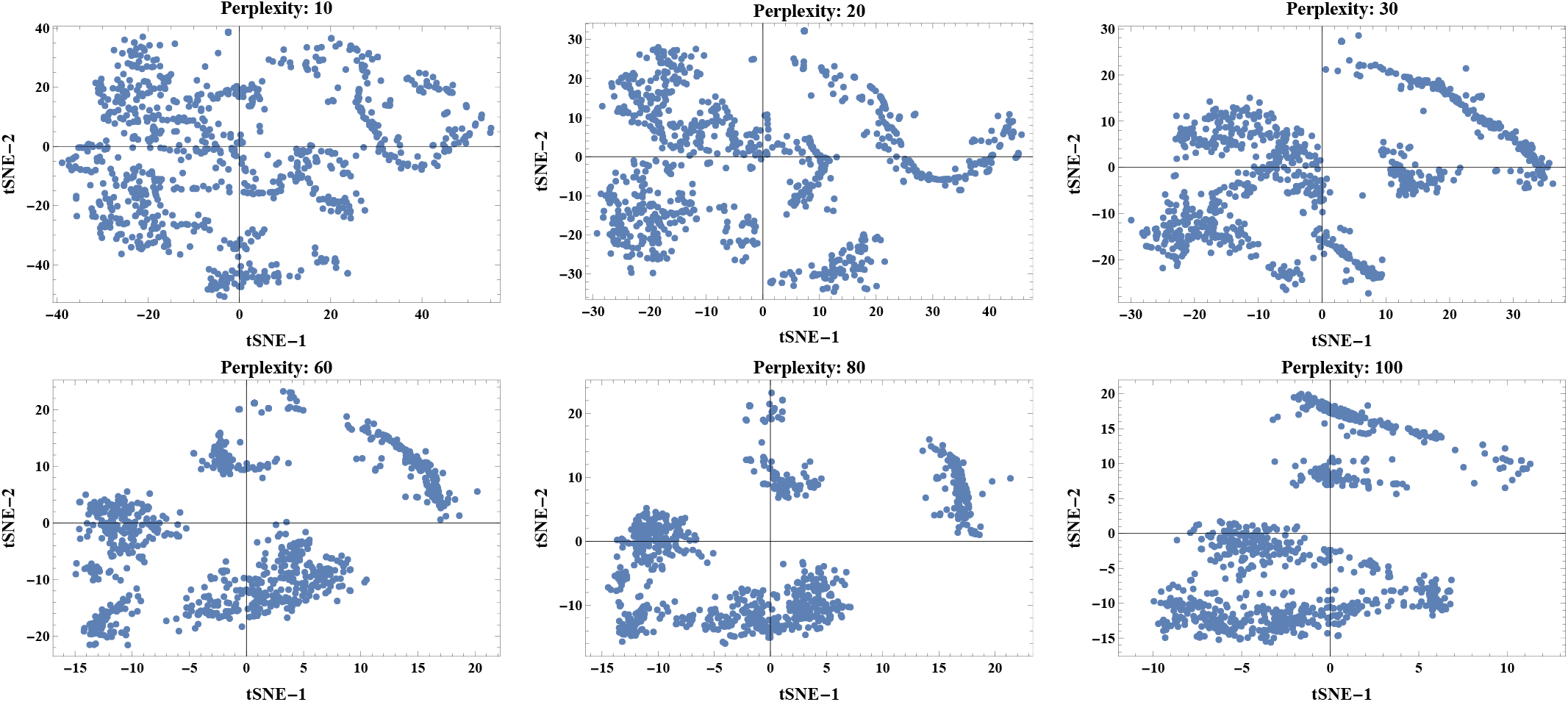
tSNE-based dimensional reduction visualizations with varying perplexity values (10, 20, 30, 60, 80, and 100). These plots demonstrate the effect of different perplexity values on the clustering and structure of the data, helping to identify the most suitable parameter for further analysis. The axes represent the two reduced dimensions obtained from tSNE.

**Figure S5:**
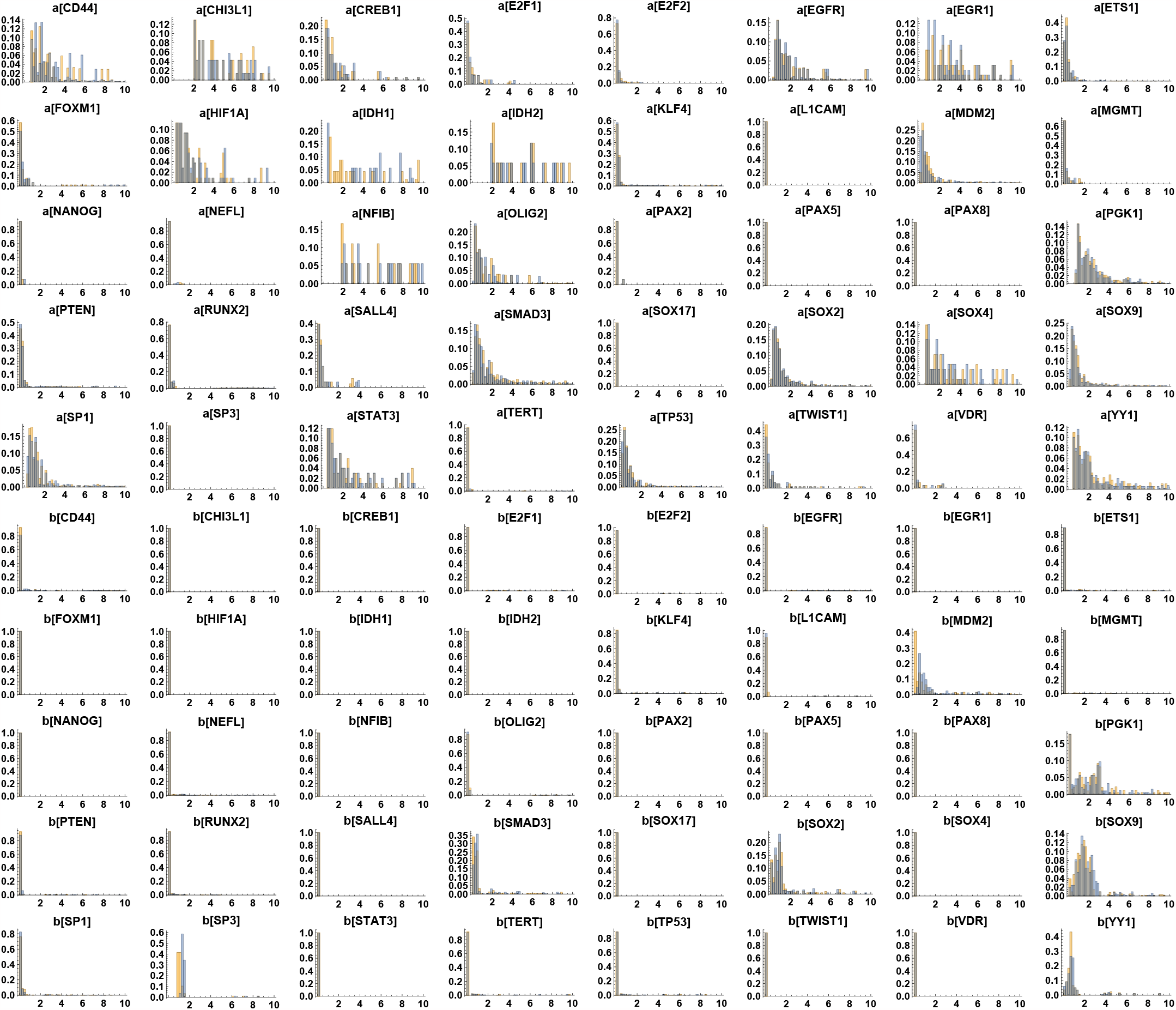
Distributions of the initial parameter estimation after clustering using k-means and NbC methods. The a[] and b[] notation refers to each gene’s activation and inhibition parameters. The orange values represent k-means, and the blue values represent NbC. These parameters multiply all interactions in each gene equation, with the same values applied across all considered clusters. The horizontal axis shows the parameter values, while the vertical axis represents normalized counts or frequencies to indicate probabilities.

**Figure S6:**
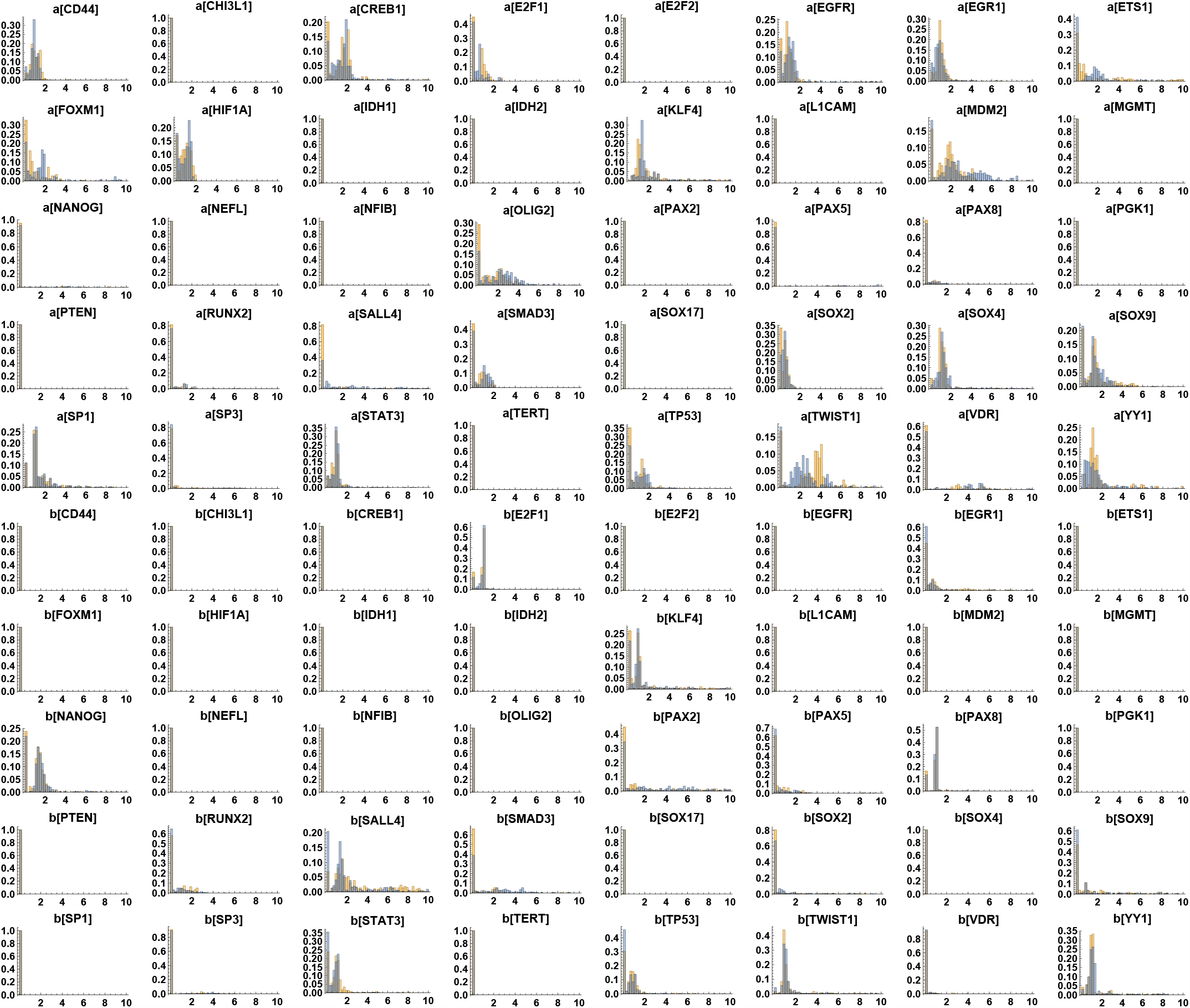
Distributions of the second parameter estimation after clustering using k-means and NbC methods. The a[] and b[] notation refers to each gene’s activation and inhibition parameters. The orange values represent k-means, and the blue values represent NbC. These second parameters multiply all interactions in all gene equations regulated by these edges. It multiplies the activation or inhibition parameters previously computed to balance the dynamics better. The horizontal axis shows the parameter values, while the vertical axis represents normalized counts or frequencies to indicate probabilities.

**Figure S7:**
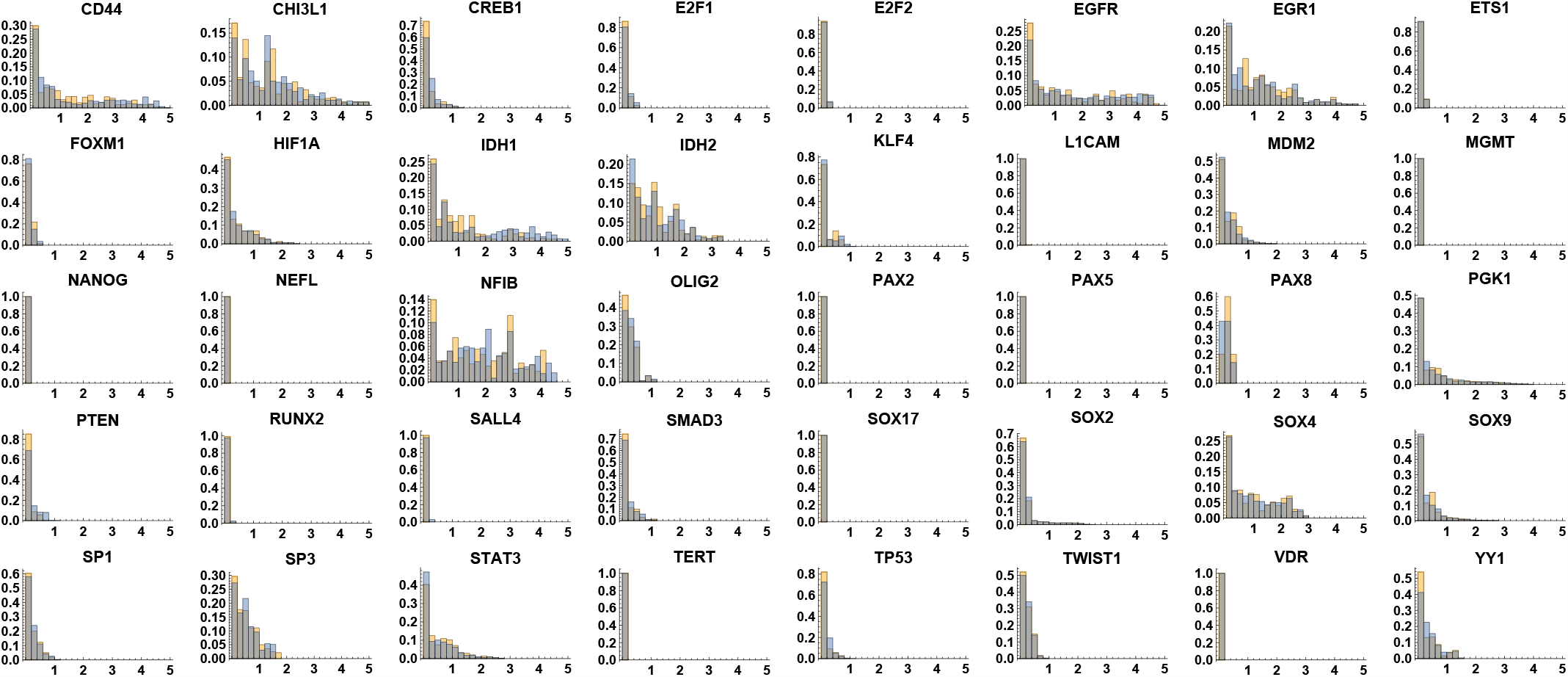
Superimposed residual distributions for the first parameters estimation after clustering using k-means and NbC methods. The orange values represent k-means, and the blue values represent NbC. For each estimation, the number of residuals for each gene was equal to the number of clusters (due to considering all clustering simultaneously). The horizontal axis shows the residual values, while the vertical axis represents normalized counts or frequencies to indicate probabilities.

**Figure S8:**
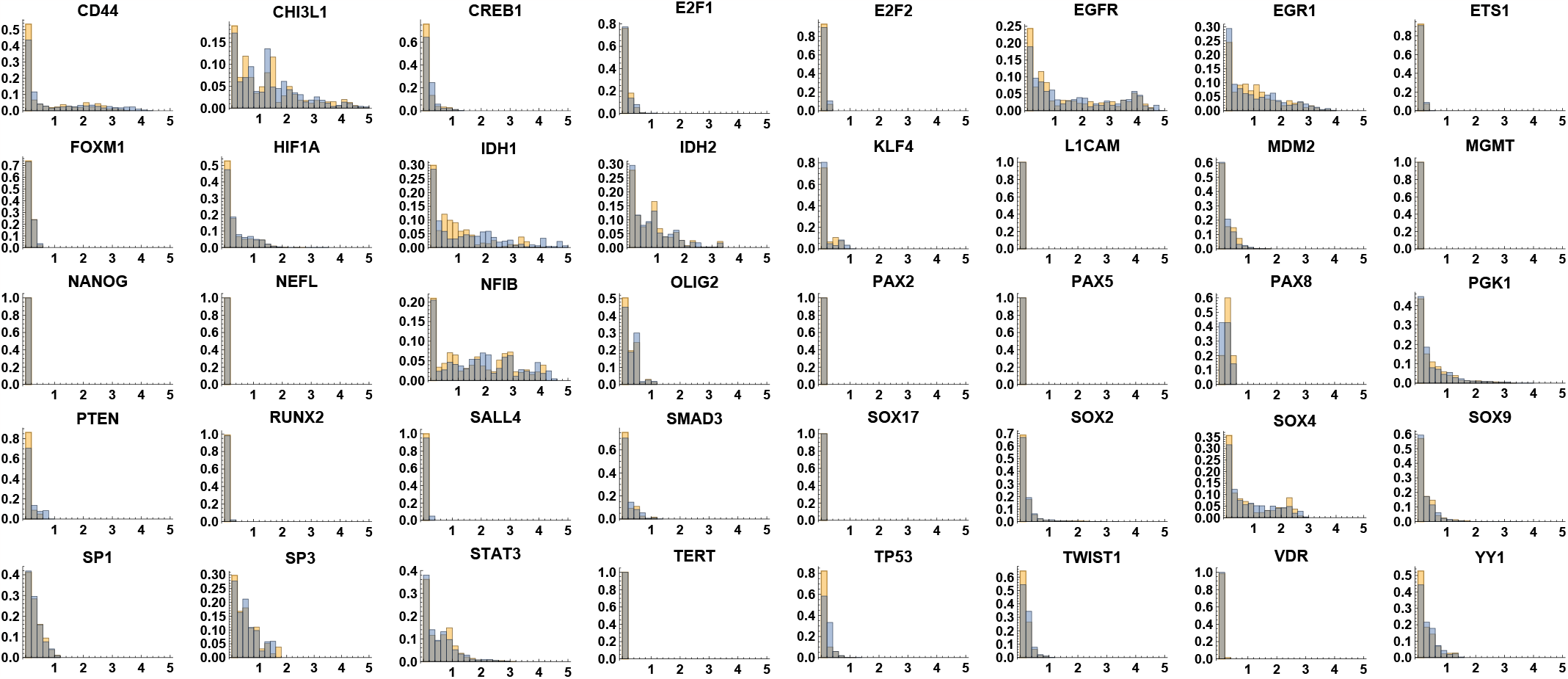
Superimposed residual distributions for the second parameters estimation after clustering using k-means and NbC methods. The orange values represent k-means, and the blue values represent NbC. For each estimation, we had the number of residuals for each gene equal to the number of clusters (due to considering all clustering simultaneously). The horizontal axis shows the residual values, while the vertical axis represents normalized counts or frequencies to indicate probabilities.

**Figure S9:**
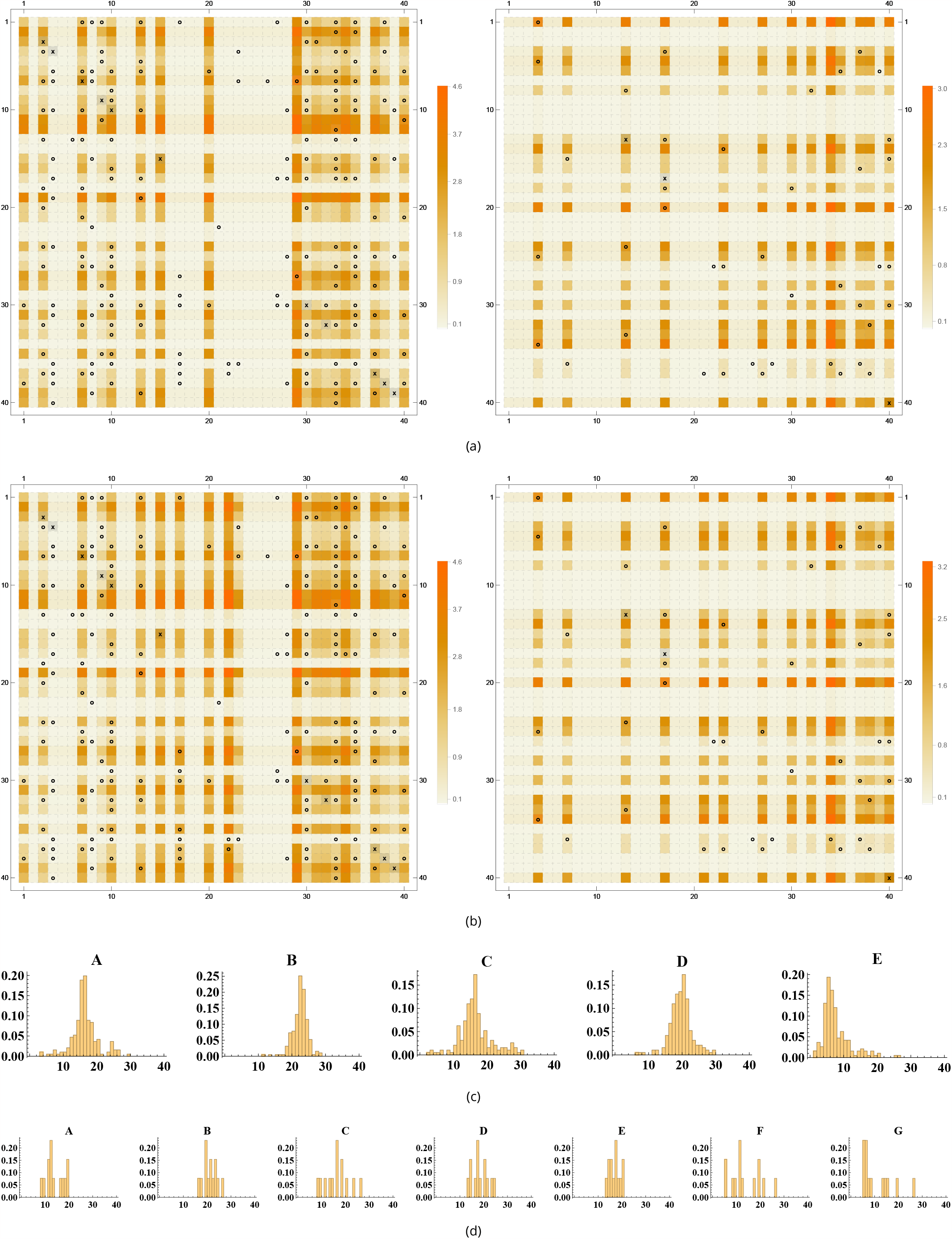
Figures (a) and (b) show the heatmap with the color gradient for the logarithm of the parameter values considering k-means and NbC, respectively. The ‘o’ represents the regulations, and the ‘x’ is the self-regulation edges in the network. Left: activation parameters. Right: inhibition parameters. The matrix indices in (a) and (b) represent each variable/vertex of the model. Figures (c) and (d) represent the distribution of the number of genes that match each cluster, considering all tested combinations of the multiplicative factors for *a, sa, b*, and *sb*. In (c) and (d), the horizontal axis represents the count of compatibility cases, and the vertical axis represents normalized counts or frequencies to indicate probabilities.

**Figure S10:**
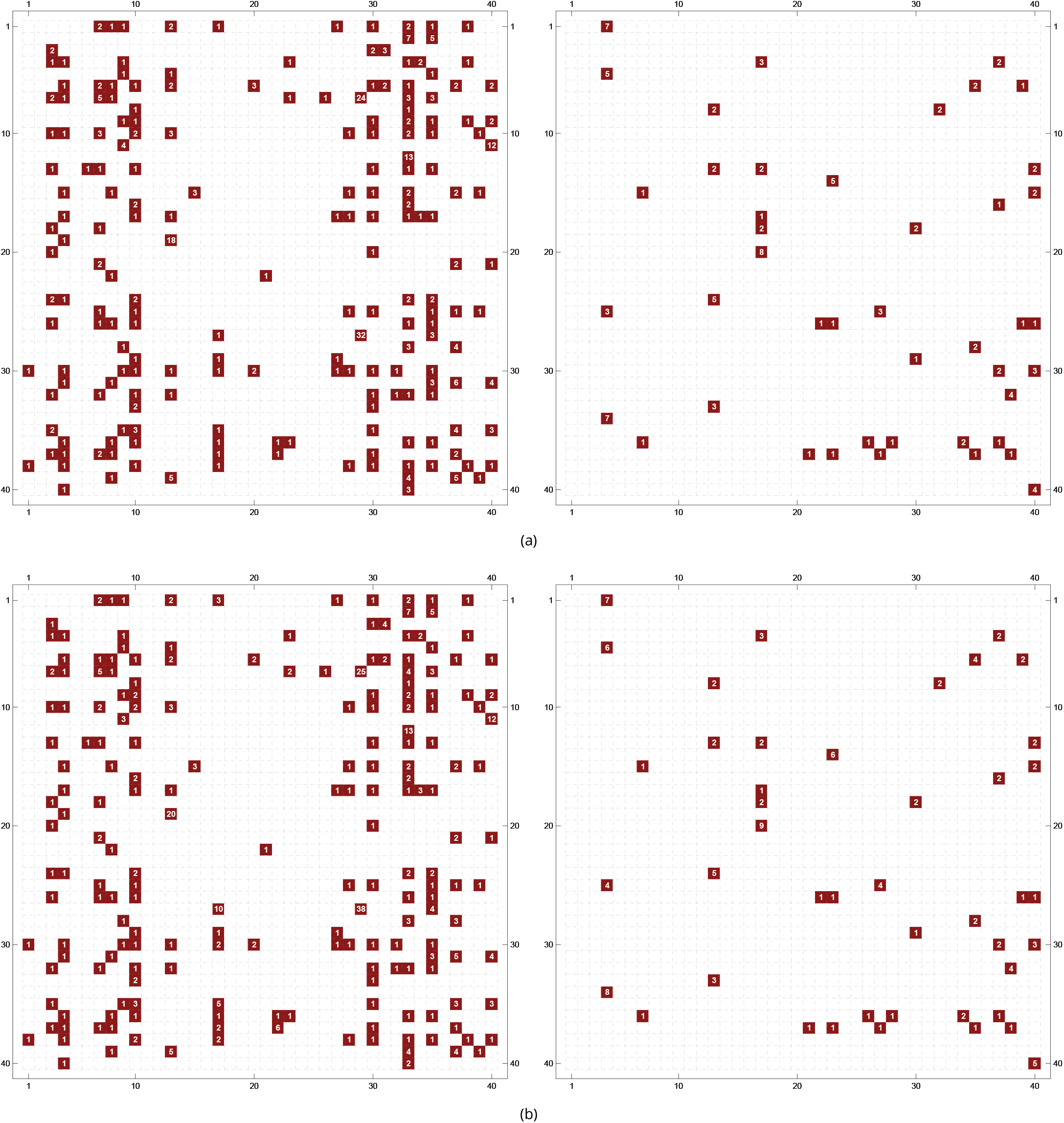
Figures (a) e (b) show the ceil() of the values of all parameters for the k-means clustering and NbC clustering, respectively. The left images correspond to activation parameters, and the right images to inhibition parameters. The ceil() was used to help the visualization of the values. The matrix indices in (a) and (b) represent each variable/vertex of the model.

**Figure S11:**
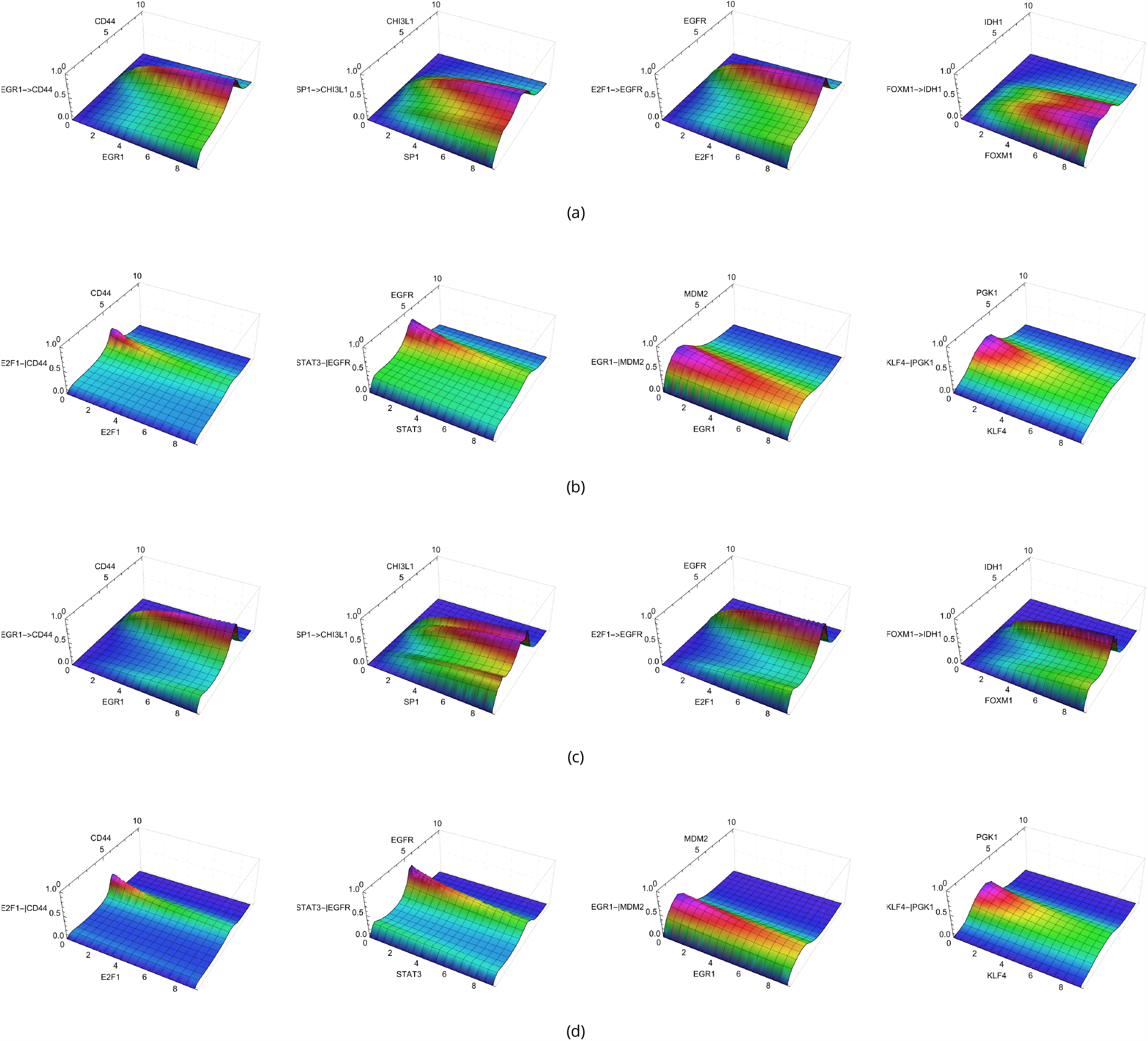
New regulation function *V* with *n* = 1 and *h*_2_(*x*) for different combinations of genes and/or transcription factors. The horizontal axis represents the transcription factor and gene quantification using the normalized amount of single-cell RNA sequencing of experimental data. The vertical axis represents the quantification of the interaction regulations. (a) Activation values using the five k-means clusters and *f*_*a*_ = 0.1; (b) Inhibitory interactions using the five k-means clusters and *f*_*b*_ = 1.3. (c) Activation values using the seven Neighborhood Contraction clusters and *f*_*a*_ = 0.1; (d) Inhibitory interactions using the seven Neighborhood Contraction clusters and *f*_*b*_ = 1.1.

**Figure S12:**
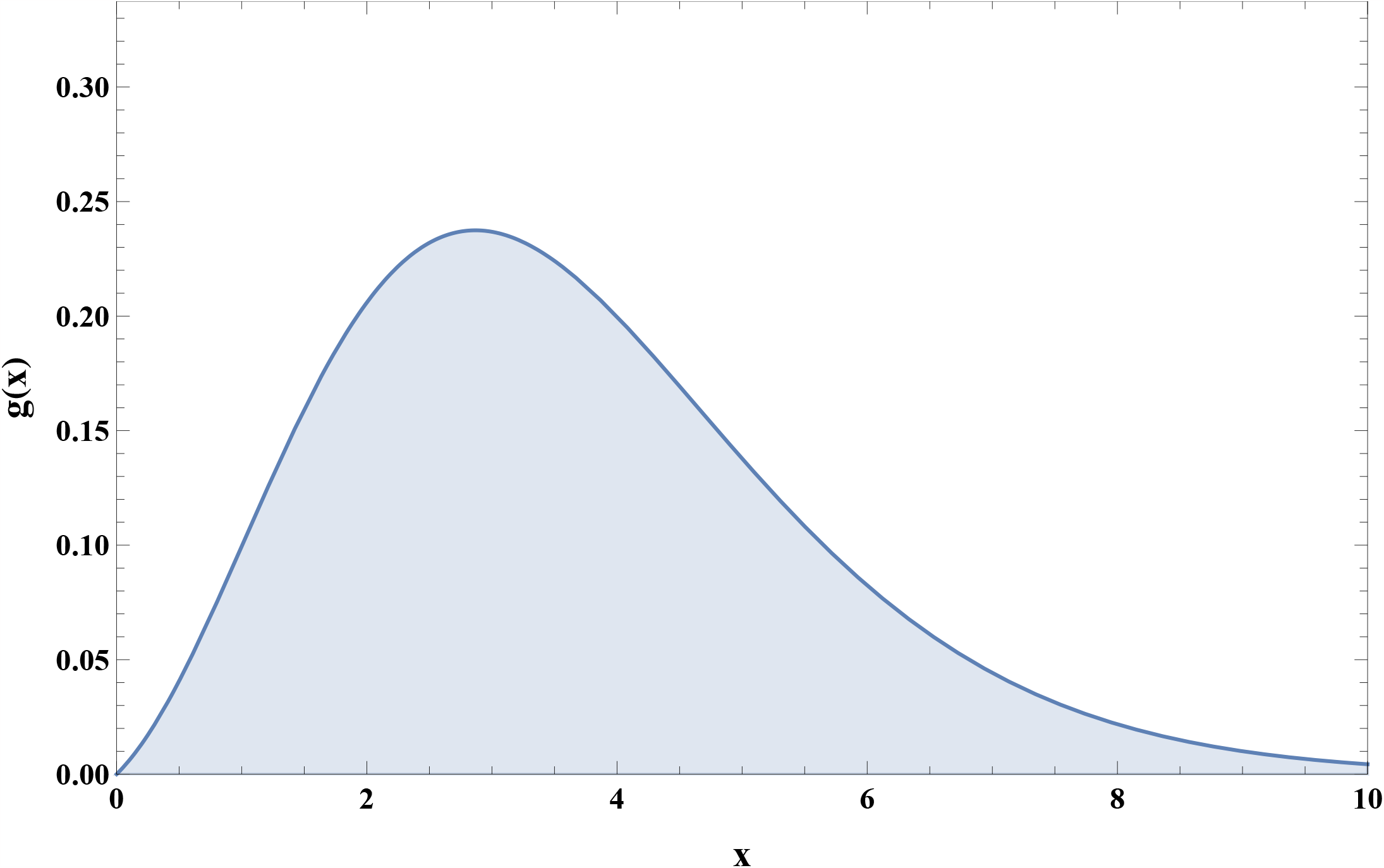
Function used for the multiplicative noise after manual verification of compatibility between the experimental data and those obtained by the simulation The horizontal axis represents the expression level quantification, and the vertical axis represents the noise amplitude before multiplication by the amplitude that was defined in the parameter estimation.

**Figure S13:**
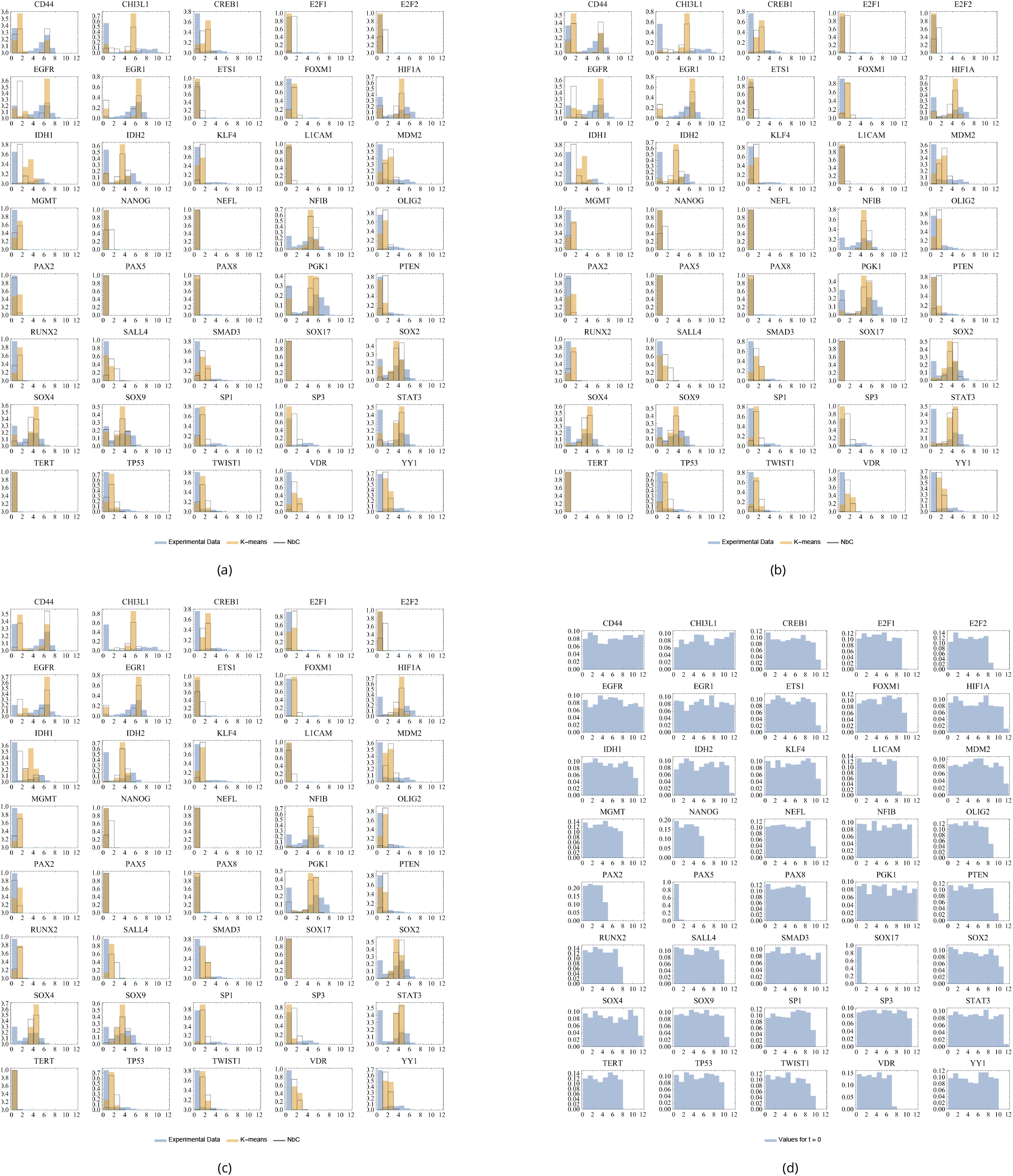
(a) to (c): Superposition of experimental (blue) and simulated (orange: K-means; black line: NbC) data for each gene of the GRN; (a) t = 50 (500 time steps); (b) t = 25 (250 time steps); (c) t = 5 (50 time steps); (d) Distribution of simulated initial conditions. The expression values are on the x-axis, and the relative frequency in the y-axis. The interval goes from 0 to 12 in unity steps. The different times were used to analyze the probability distribution evolution over time. The horizontal axis shows the expression values, while the vertical axis represents normalized counts or frequencies to indicate probabilities.

**Figure S14:**
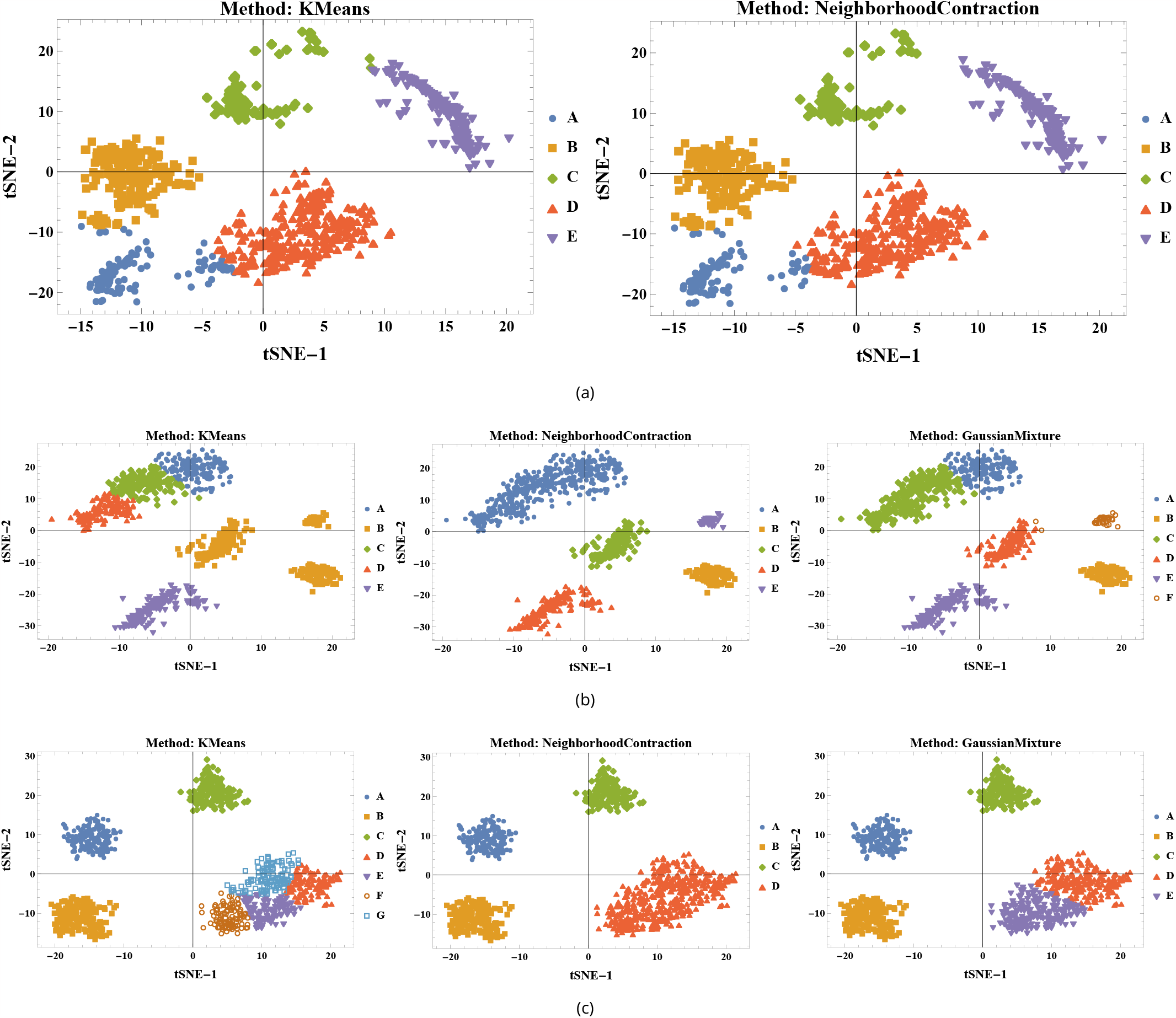
Clustering results using different methods. The axes represent the two reduced dimensions obtained for each tSNE. (a) Experimental data; (b) simulated data after k-means clustering; (c) simulated data after Nbc clustering. The letters represent the identified clusters. These clusters were defined in this 2-dimensional space and were used solely to assess the clustering quality. The actual clustering was performed in the 4-marker dimensional space. All methods used the built-in Mathematica functions with PerformanceGoal set to “Quality”, CriterionFunction set to “StandardDeviation”, and DistanceFunction set to EuclideanDistance.

**Figure S15:**
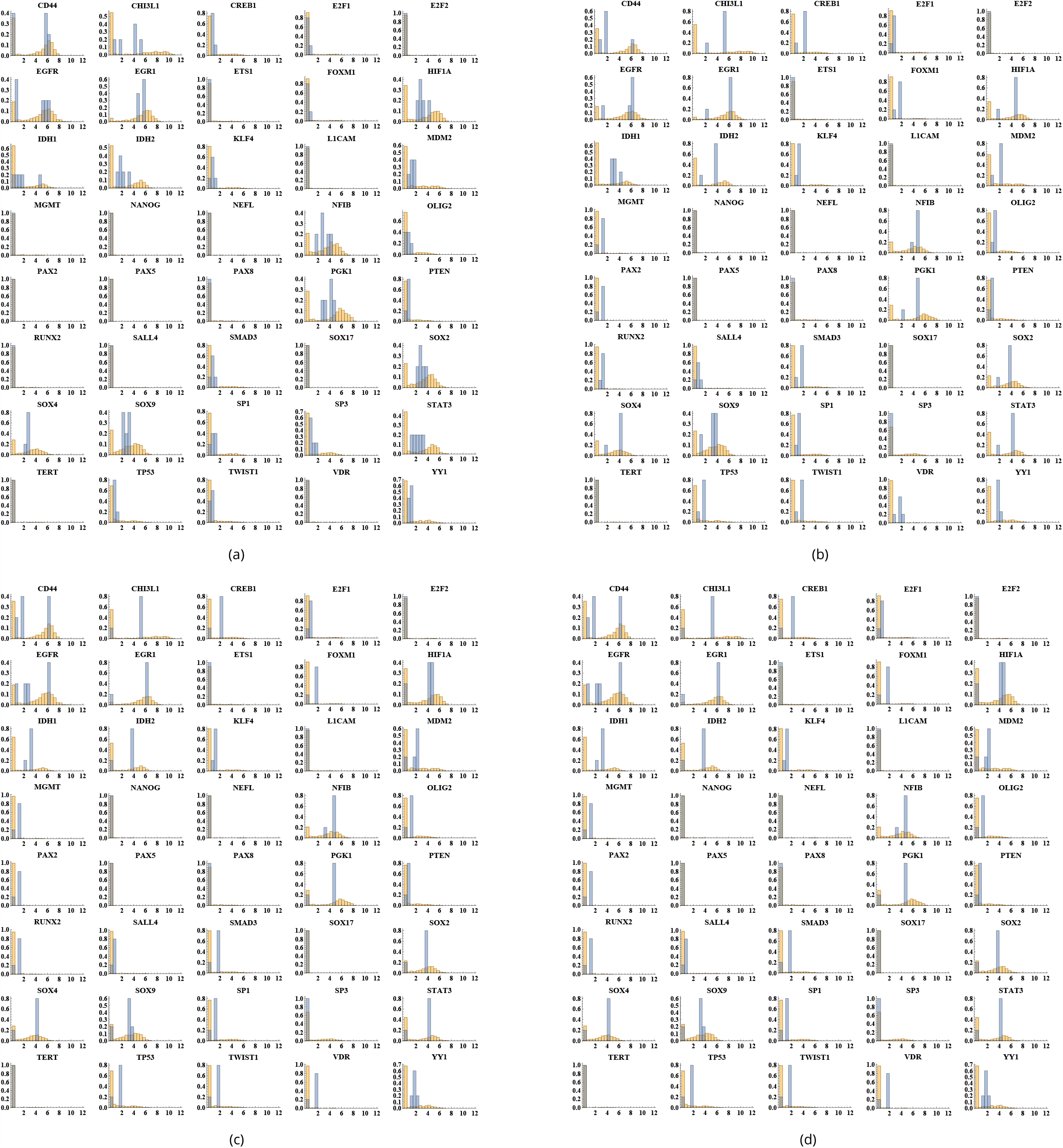
Cluster centroids superimposed on experimental data distributions for each gene. (a) Centroids found using k-means with experimental data; (b) Centroids found using k-means with simulated data after parameter estimation using k-means clusters; (c) Centroids found using NbC; (d) Centroids found using Gaussian Mixture. The horizontal axis shows the expression values, while the vertical axis represents normalized counts or frequencies to indicate probabilities.

**Figure S16:**
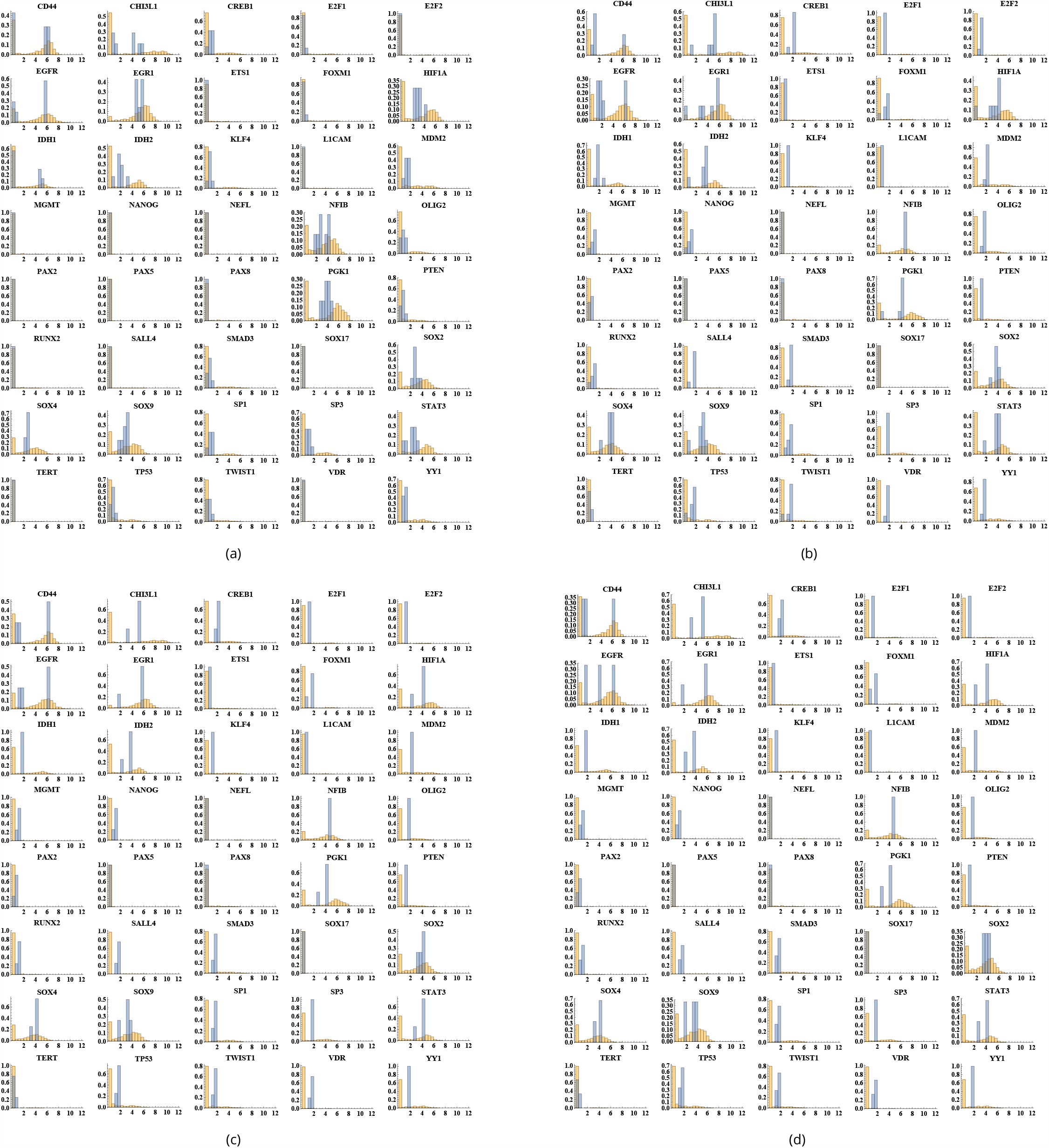
Cluster centroids superimposed on experimental data distributions for each gene. (a) Centroids found using NbC with experimental data; (b) Centroids found using k-means with simulated data after parameter estimation using NbC clusters; (c) Centroids found using NbC; (d) Centroids found using Gaussian Mixture. The horizontal axis shows the expression values, while the vertical axis represents normalized counts or frequencies to indicate probabilities.

**Figure S17:**
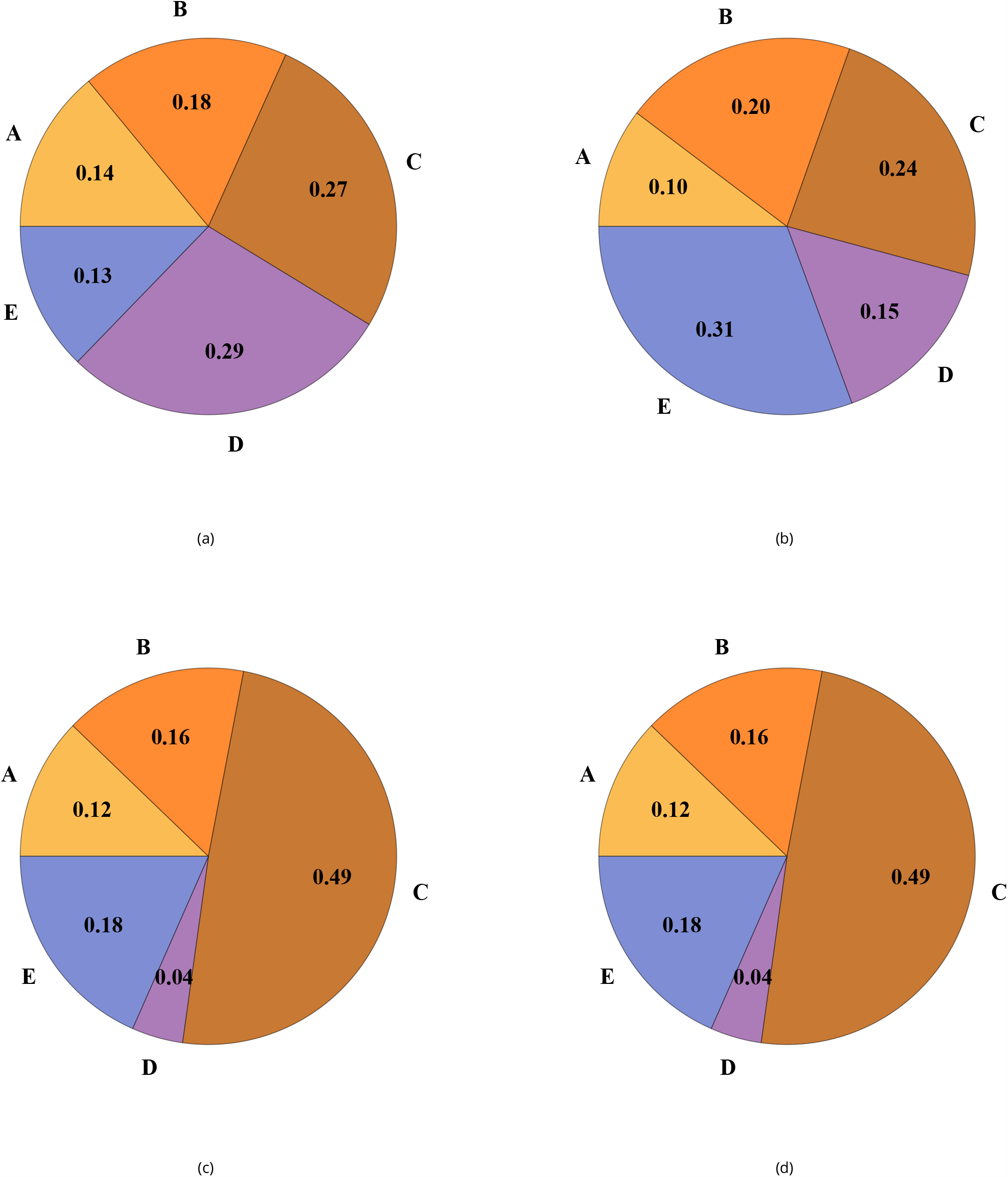
Pie charts representing the number of elements within each cluster. (a) Experimental data clusters using k-means; (b) to (d) Clusters of simulated data after parameter estimation using k-means clusters. (b) k-means; (c) NbC; (d) Gaussian Mixture.

**Figure S18:**
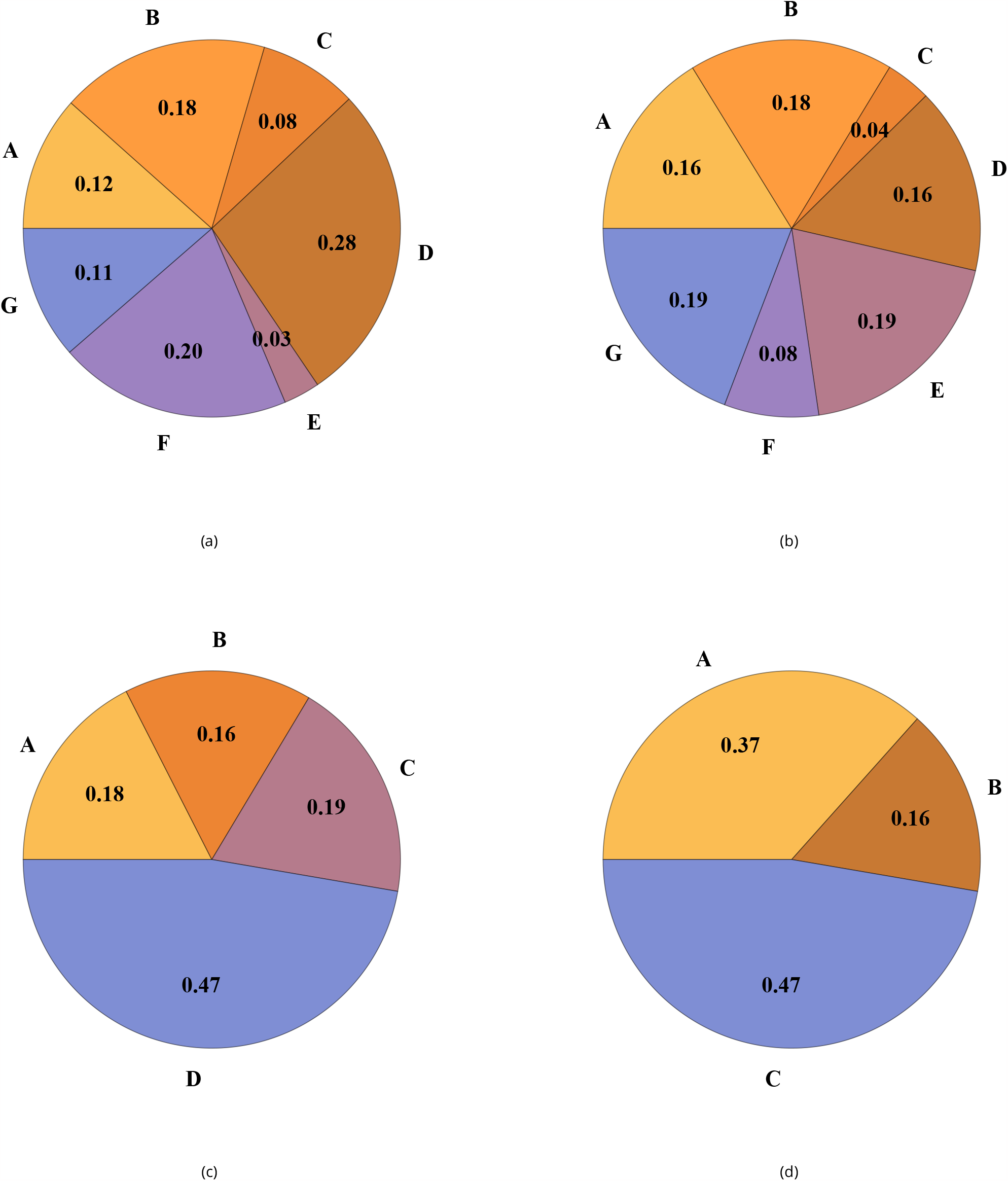
Pie charts representing the number of elements within each cluster. (a) Experimental data clusters using NbC; (b) to (d) Clusters of of simulated data after parameter estimation using NbC clusters. (b) k-means; (c) NbC; (d) Gaussian Mixture.

**Figure S19:**
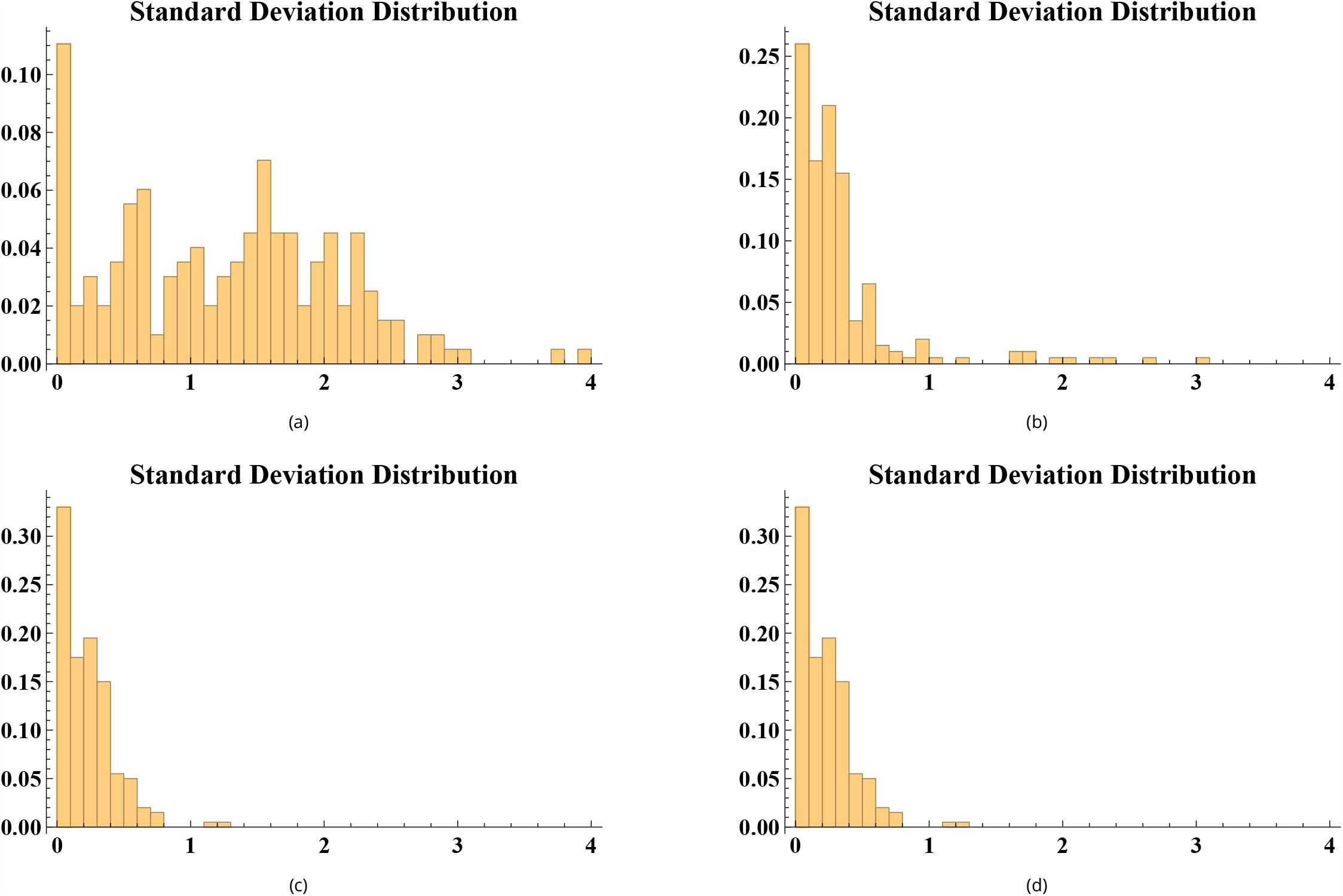
Standard deviation distribution of all genes within all clusters for each clustering method. (a) k-means applied to experimental data; (b) to (d) Clustering of simulated data obtained after parameter estimation using k-means clusters. (b) k-means; (c) NbC; (d) Gaussian Mixture. The horizontal axis shows the standard deviation values, while the vertical axis represents normalized counts or frequencies to indicate probabilities.

**Figure S20:**
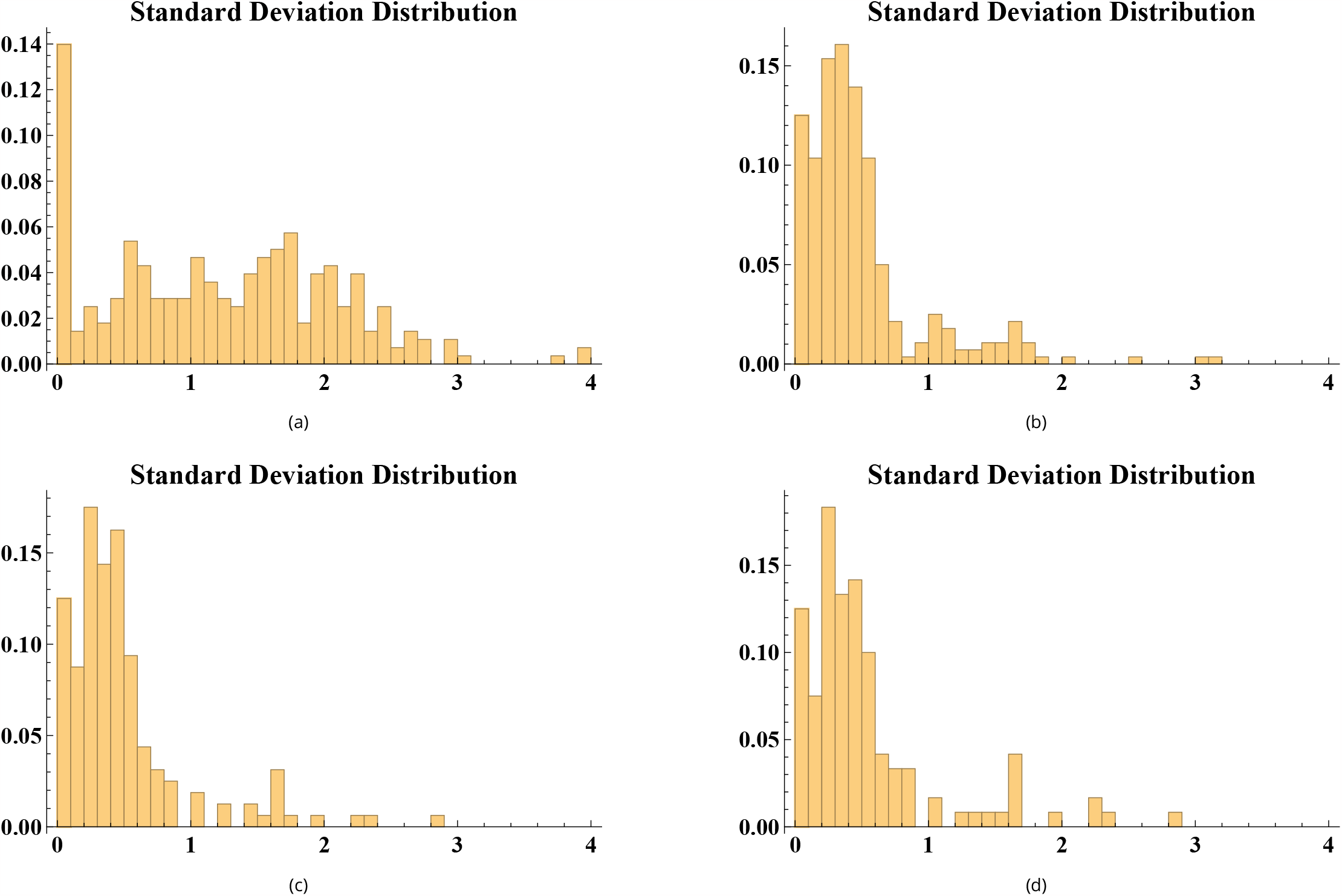
Standard deviation distribution of all genes within all clusters for each clustering method. (a) NbC applied to experimental data; (b) to (d) Clustering of simulated data obtained after parameter estimation using NbC clusters. (b) k-means; (c) NbC; (d) Gaussian Mixture. The horizontal axis shows the standard deviation values, while the vertical axis represents normalized counts or frequencies to indicate probabilities.

**Figure S21:**
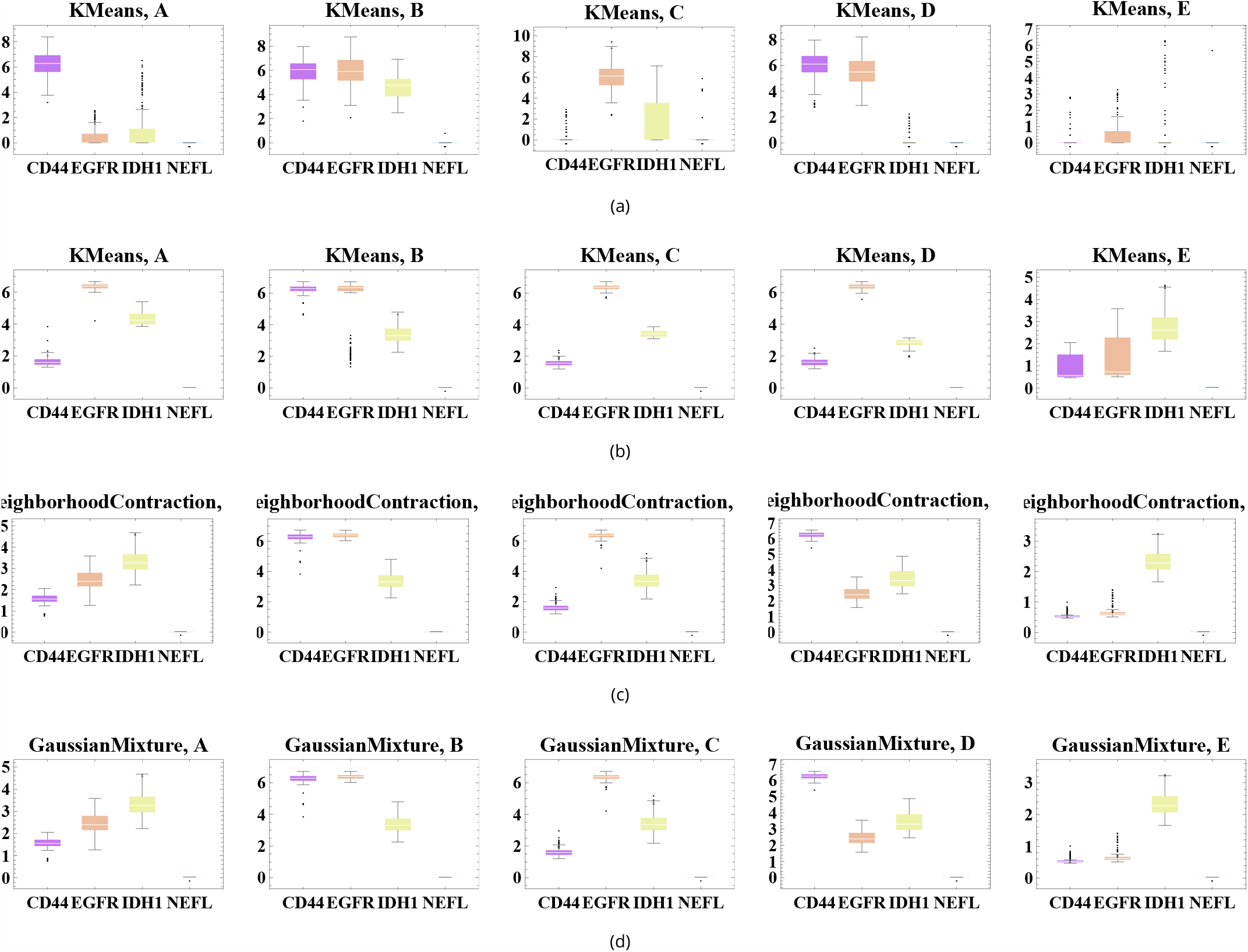
Boxplots of gene expression distributions for different clustering methods. (a) k-means applied to experimental data; (b) k-means applied to simulated data after parameter estimation using k-means clusters; (c) NbC applied to simulated data after parameter estimation using k-means clusters; (d) Gaussian Mixture applied to simulated data after parameter estimation using k-means clusters. The horizontal axis displays each considered marker gene, while the vertical axis represents the respective expression values.

**Figure S22:**
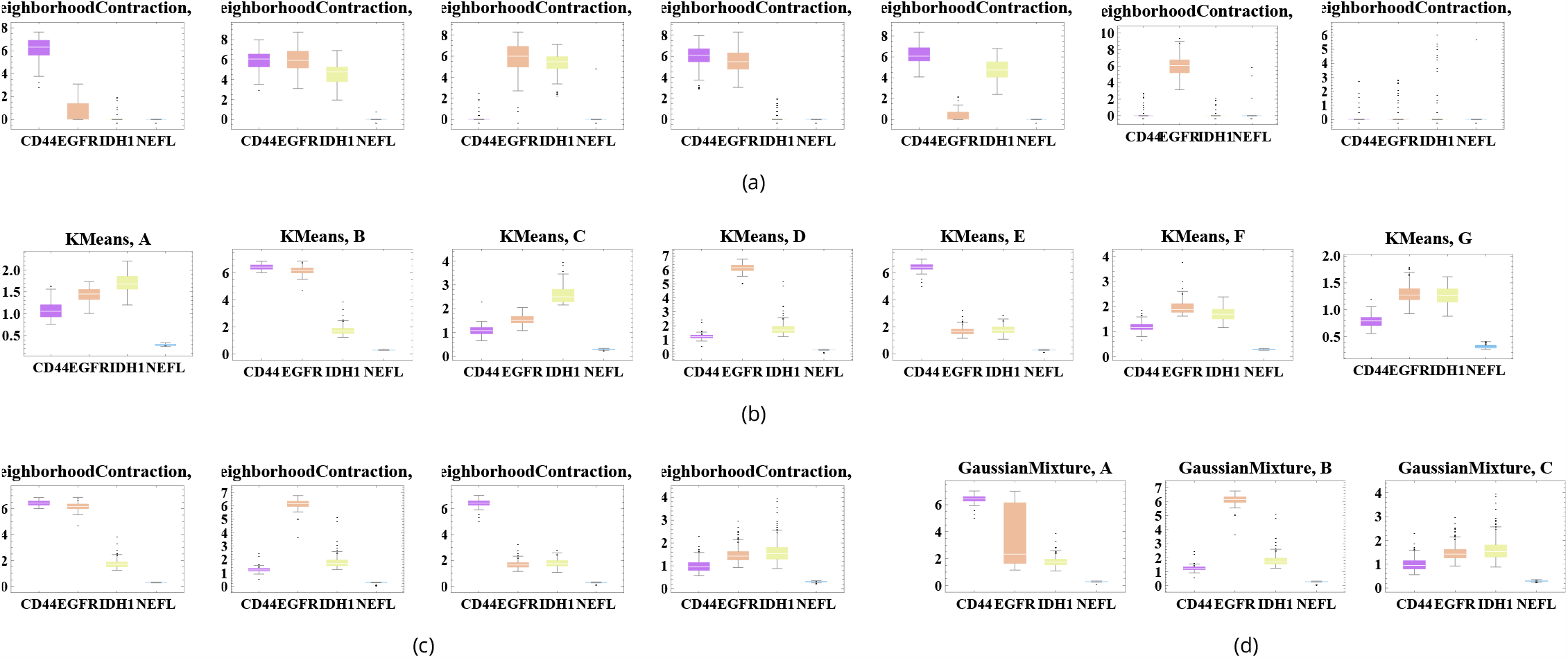
Boxplots of gene expression distributions for different clustering methods. (a) NbC applied to experimental data; (b) k-means applied to simulated data after parameter estimation using NbC clusters; (c) NbC applied to simulated data after parameter estimation using NbC clusters; (d) Gaussian Mixture applied to simulated data after parameter estimation using NbC clusters. The horizontal axis displays each considered marker gene, while the vertical axis represents the respective expression values.

**Figure S23:**
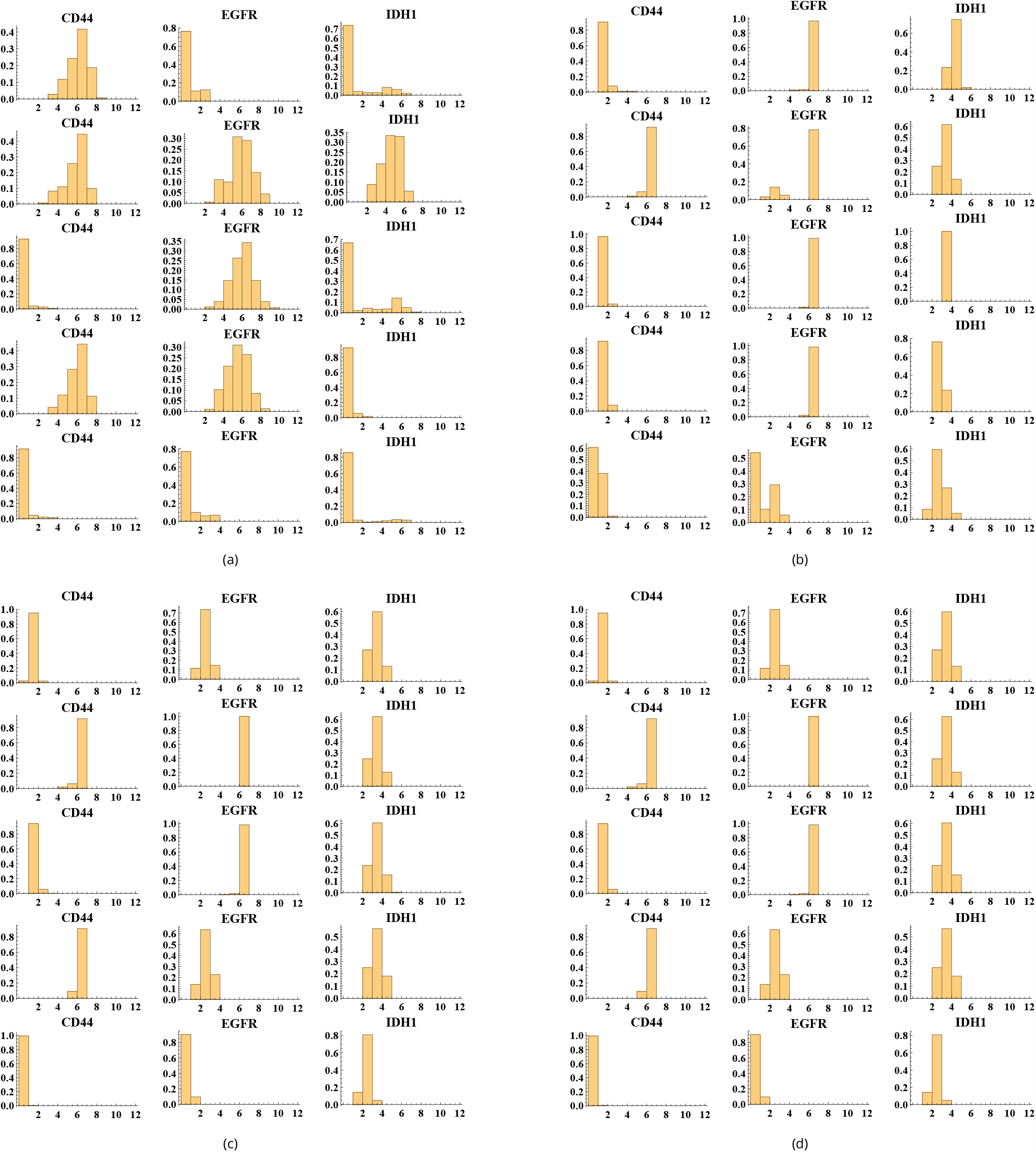
Gene expression distribution of markers for each cluster in different clustering methods. (a) k-means applied to experimental data; (b) k-means applied to simulated data after parameter estimation using k-means clusters; (c) NbC applied to simulated data after parameter estimation using k-means clusters; (d) Gaussian Mixture applied to simulated data after parameter estimation using k-means clusters. Each horizontal line shows the marker distribution for each respective cluster. For each case, the cluster identification ranges from A to E. The horizontal axis shows the expression values, while the vertical axis represents normalized counts or frequencies to indicate probabilities.

**Figure S24:**
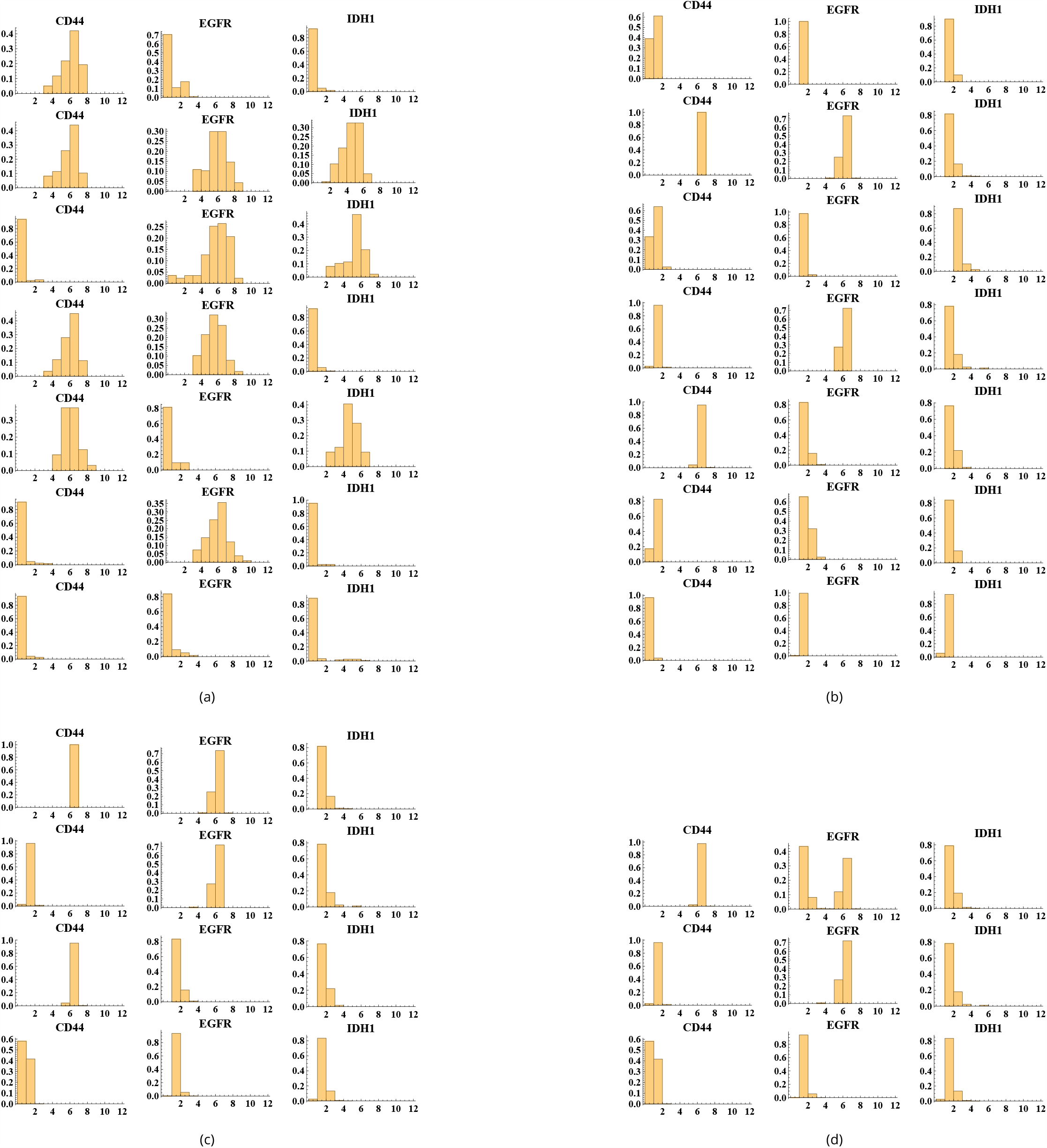
Gene expression distribution of markers for each cluster in different clustering methods. (a) NbC applied to experimental data; (b) k-means applied to simulated data after parameter estimation using NbC clusters; (c) NbC applied to simulated data after parameter estimation using NbC clusters; (d) Gaussian Mixture applied to simulated data after parameter estimation using NbC clusters. Each horizontal line shows the marker distribution for each respective cluster. For each case, the identification of the cluster ranges from A to its respective letter. The horizontal axis shows the expression values, while the vertical axis represents normalized counts or frequencies to indicate probabilities.

**Figure S25:**
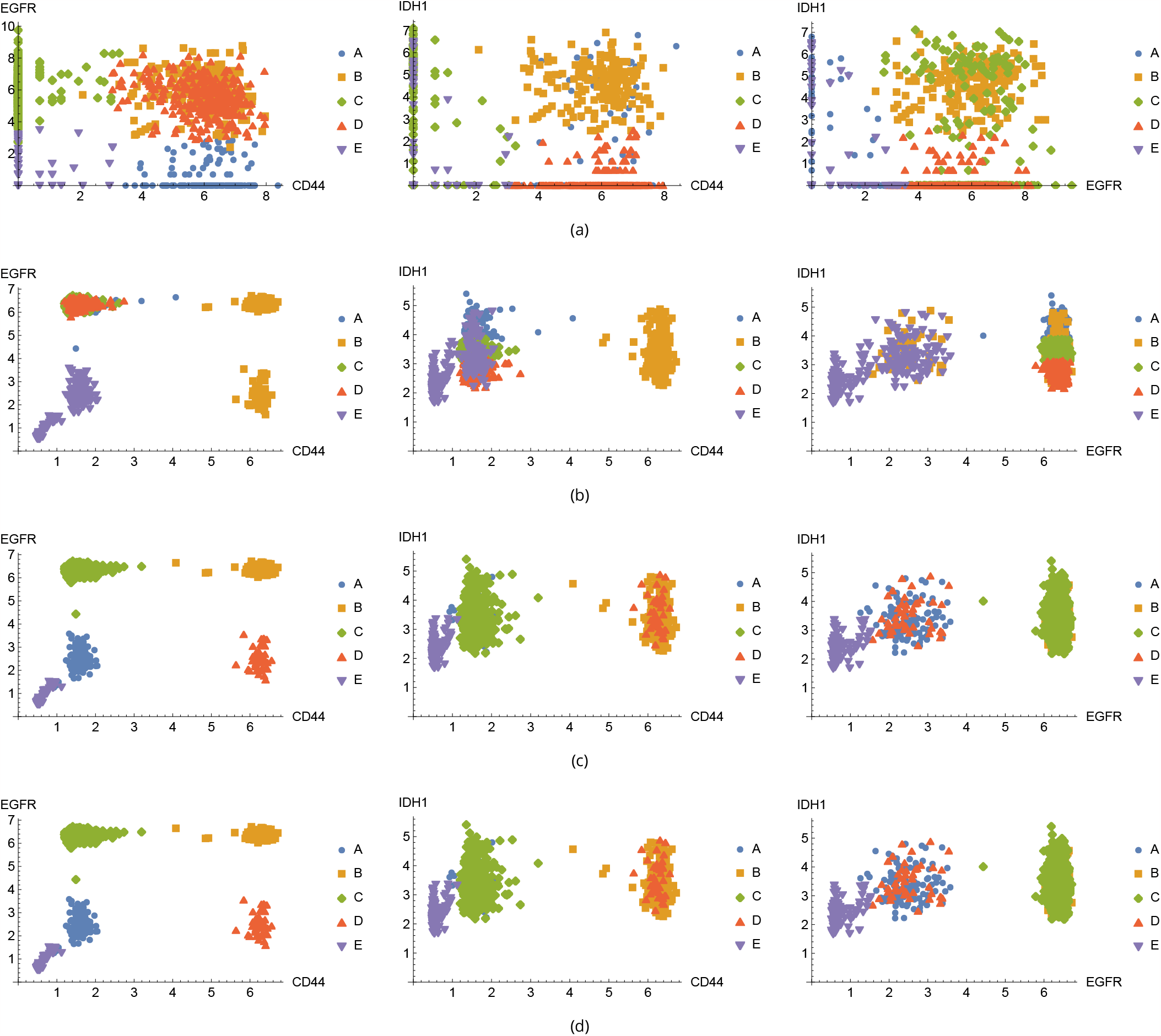
Scatter plots for combinations of marker gene expressions, excluding NEFL due to predominantly zero values. (a) Experimental data with five k-means clusters; (b) to (d) Simulated data after parameter estimation using k-means clusters. (b) Clustering with k-means; (c) Clustering with NbC; (d) Clustering with Gaussian Mixture. Each scatter plot represents the relationship between the expressions of different marker genes in each clustering method. The horizontal and the vertical axes show the expression values of each marker gene, while the different colors and shapes represent their corresponding cluster.

**Figure S26:**
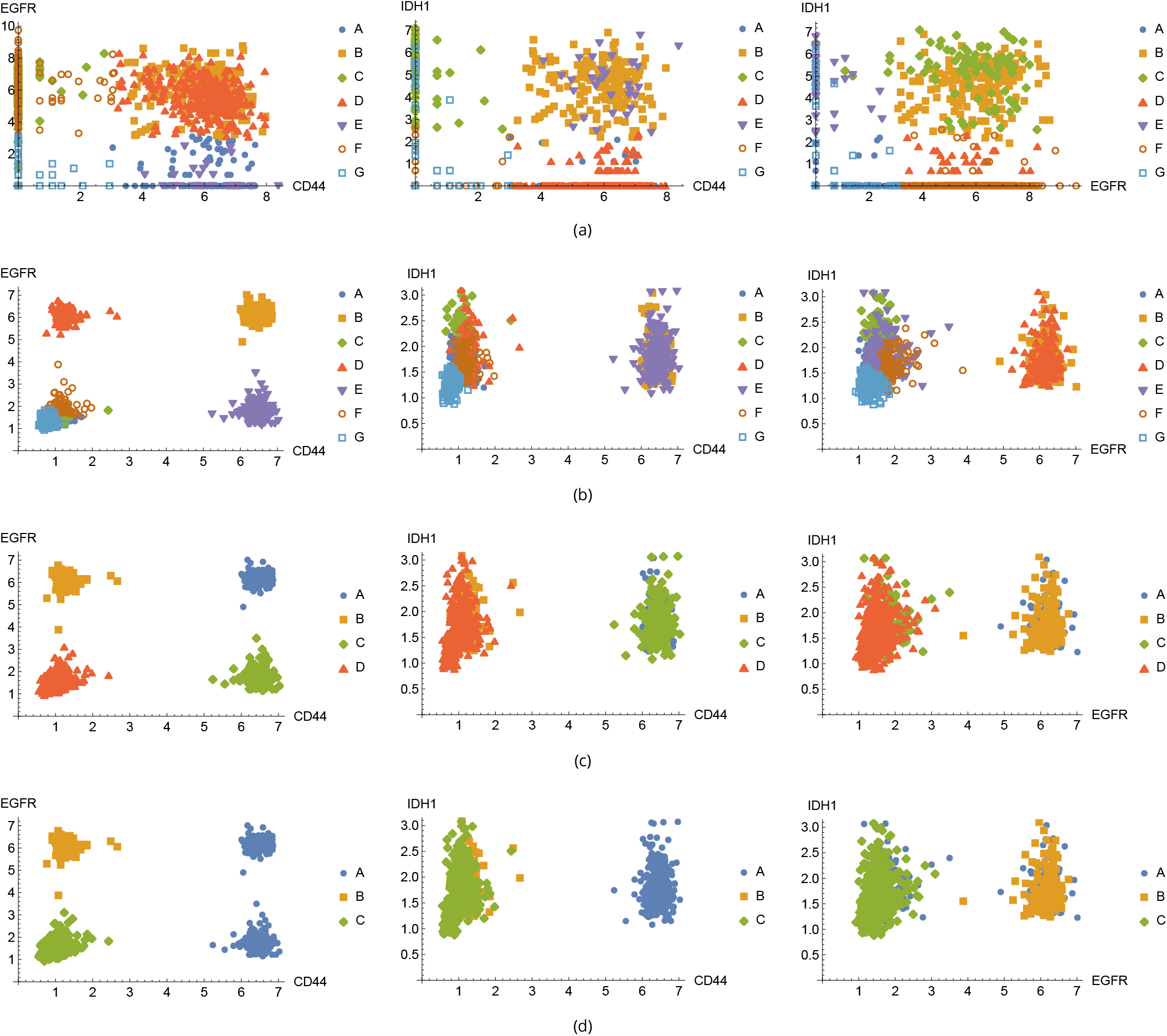
Scatter plots for combinations of marker gene expressions, excluding NEFL due to predominantly zero values. (a) Experimental data with seven NbC clusters; (b) to (d) Simulated data after parameter estimation using NbC clusters. (b) Clustering with k-means; (c) Clustering with NbC; (d) Clustering with Gaussian Mixture. Each scatter plot represents the relationship between the expressions of different marker genes in each clustering method. The horizontal and the vertical axes show the expression values of each marker gene, while the different colors and shapes represent their corresponding cluster.

**Figure S27:**
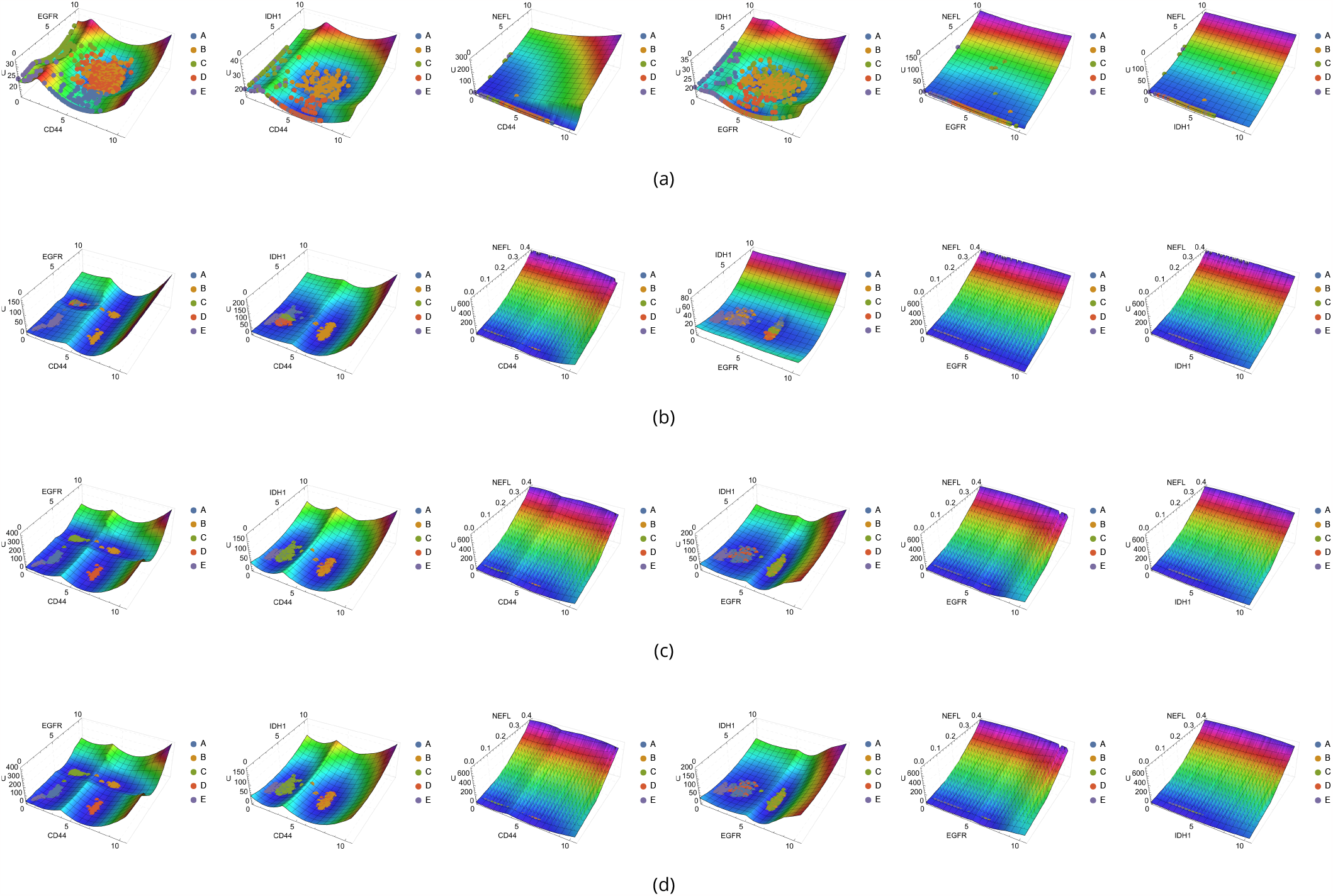
(Epi)genetic landscapes for experimental and simulated data, with experimental and simulated points overlaid for compatibility visualization. (a) Landscape for experimental data with k-means clusters; (b) Landscape for simulated data after parameter estimation and clustering with k-means; (c) Landscape for simulated data after parameter estimation and clustering with NbC; (d) Landscape for simulated data after parameter estimation and clustering with Gaussian Mixture. These landscapes visually represent the compatibility between experimental and simulated data in each clustering method. The horizontal axes show the expression values of each marker gene, while the vertical axis represents the values of the landscape. The different colors and shapes correspond to the respective clusters.

**Figure S28:**
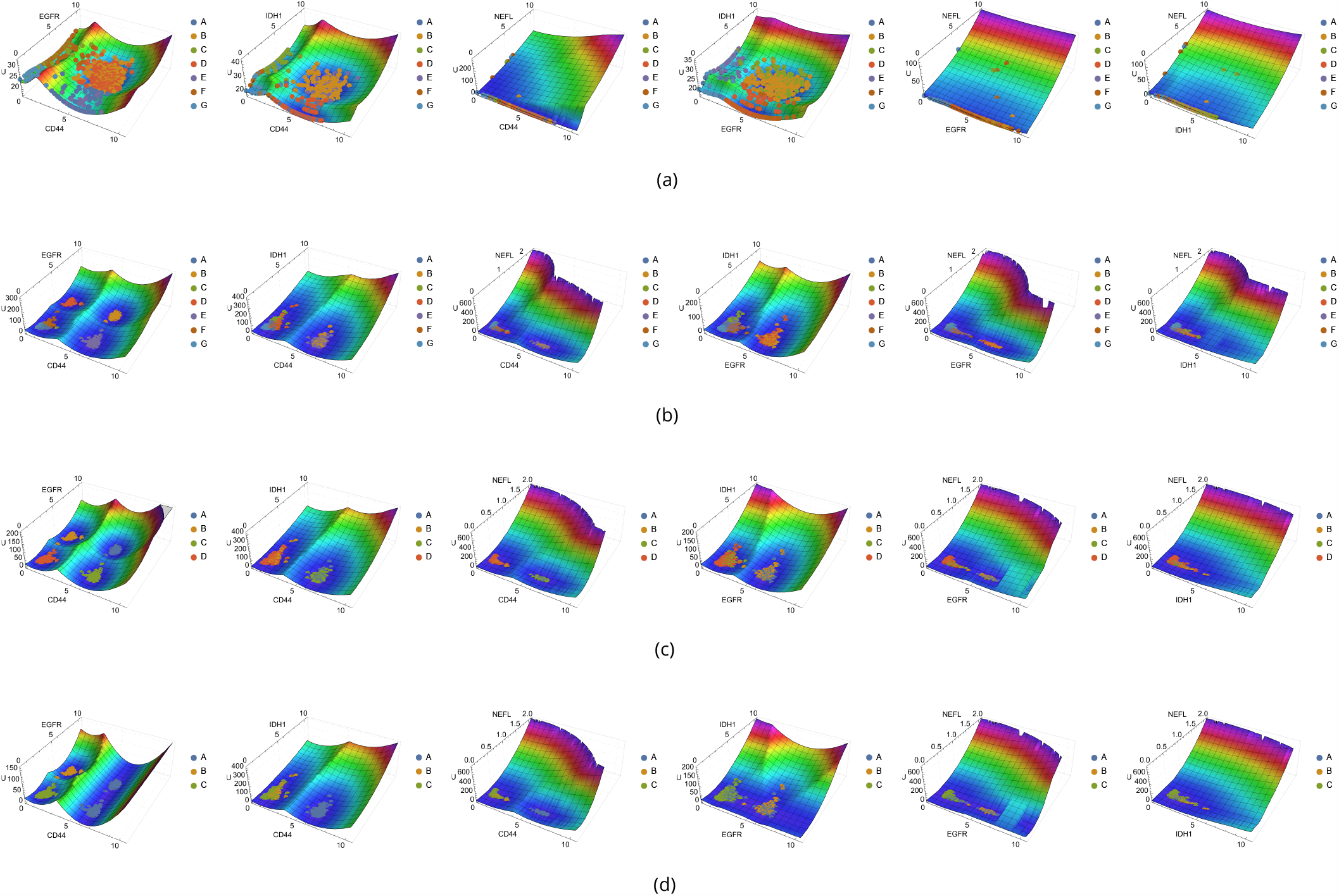
(Epi)genetic landscapes for experimental and simulated data, with experimental and simulated points overlaid for compatibility visualization. (a) Landscape for experimental data with NbC clusters; (b) Landscape for simulated data after parameter estimation and clustering with k-means; (c) Landscape for simulated data after parameter estimation and clustering with NbC; (d) Landscape for simulated data after parameter estimation and clustering with Gaussian Mixture. These landscapes visually represent the compatibility between experimental and simulated data in each clustering method. The horizontal axes show the expression values of each marker gene, while the vertical axis represents the values of the landscape. The different colors and shapes correspond to the respective clusters.

**Figure S29:**
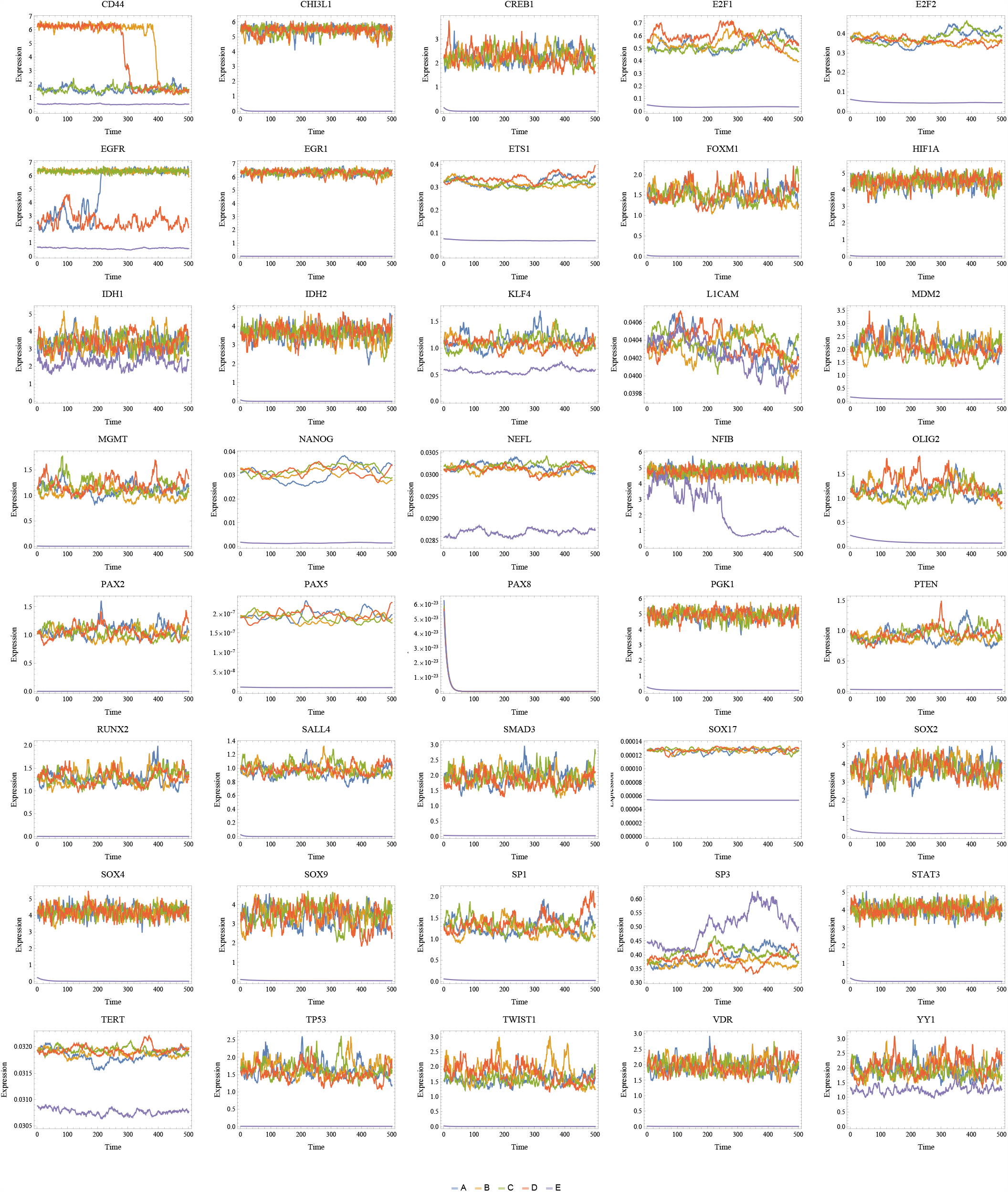
Trajectory plots for each basin, showcasing the system’s dynamics. Each panel represents a different gene, with five trajectories, one for each cluster. The horizontal axis represents time, and the vertical axis shows the expression level of the gene. The trajectories illustrate the time evolution from initial conditions as the centroid of each cluster. These visualizations provide insights into the internal dynamics of each basin and help identify potential transitions between clusters.

**Figure S30:**
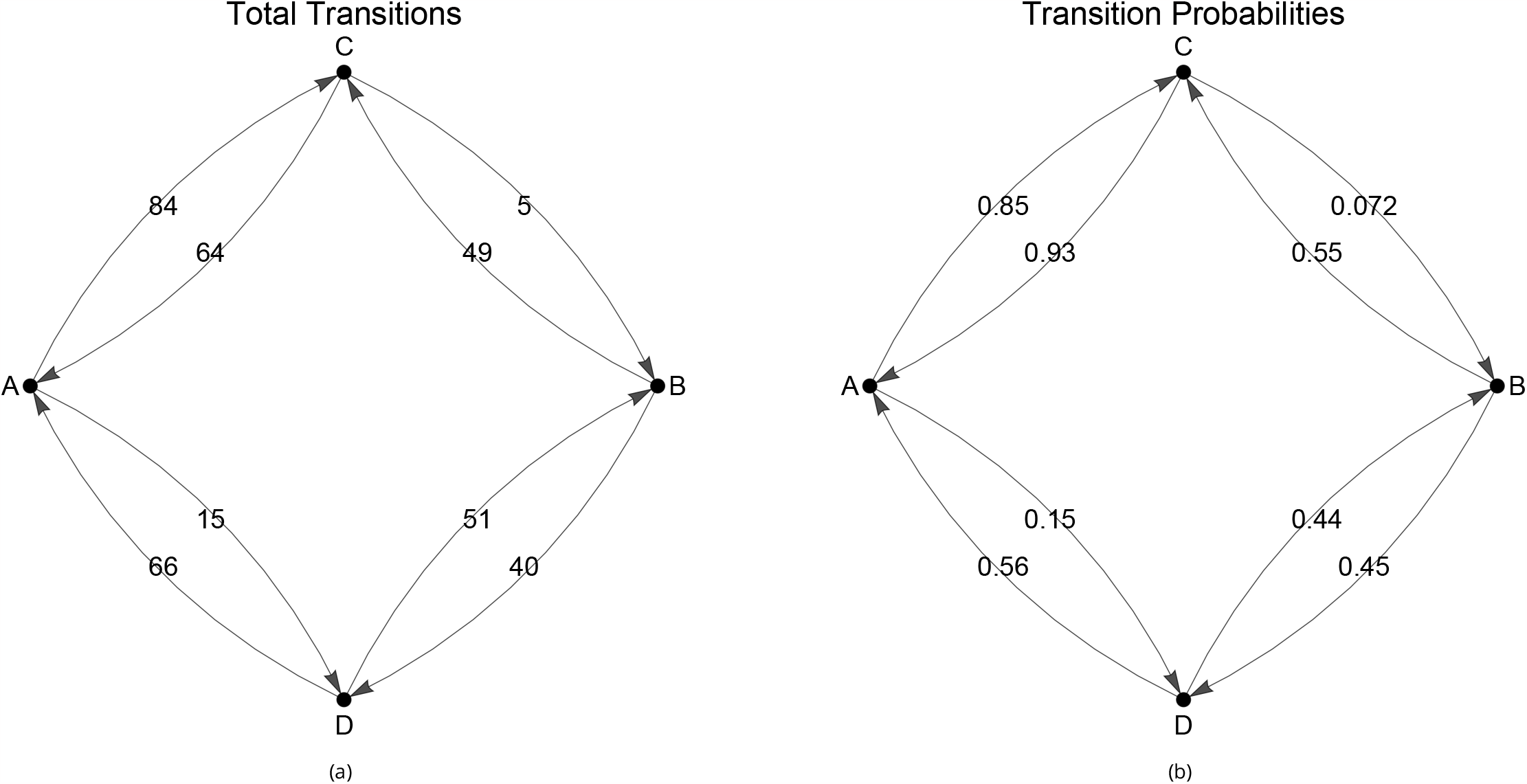
Transition graphs illustrating the connections between different basins. (a) Graph representation of transitions between basins, with vertices representing basins and edges representing observed transitions. (b) The same transition graph as in (a), but with edge weights representing the probabilities of transitions between basins. These visualizations provide insights into the potential transition pathways and their relative likelihood within the system.

**Figure S31:**
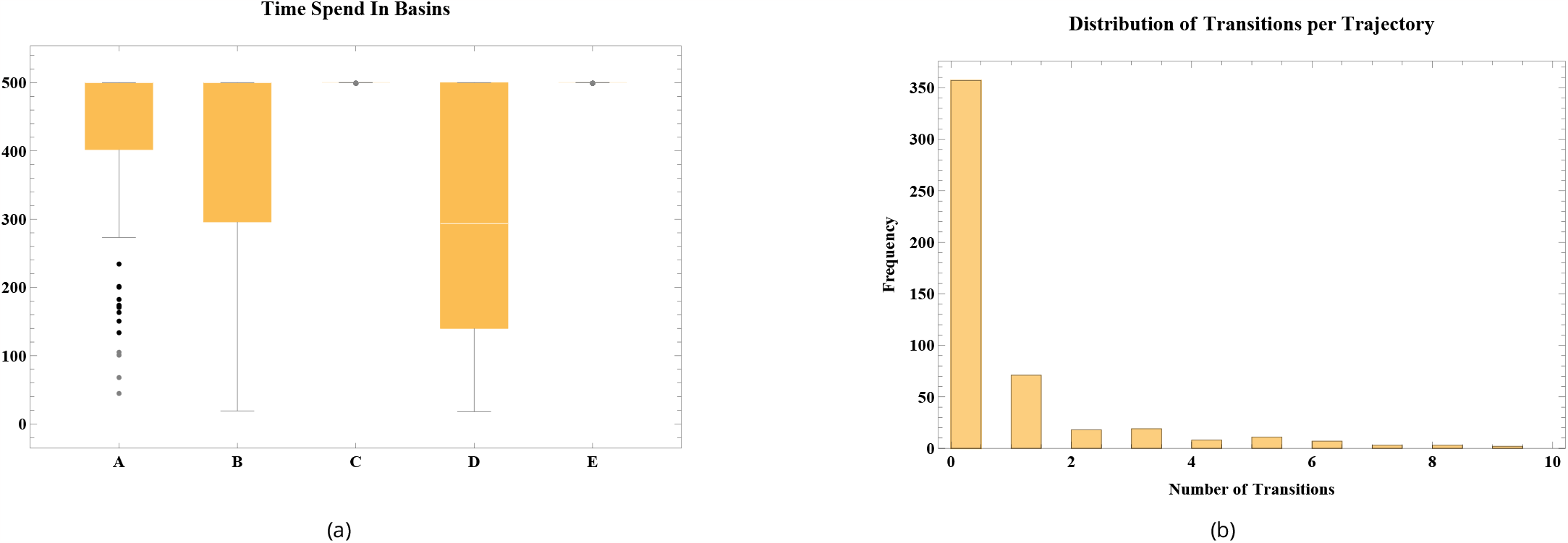
Analysis of time spent in basins before a transition and the frequency of transitions per trajectory. (a) Box plots of the distribution of time spent in each basin across all trajectories before they present a transition, providing insights into the relative stability of different basins. The vertical axis represents the time spent in the basin, while the horizontal axis the correspondent basins. (b) Histogram showing the frequency of transitions between basins in each trajectory, highlighting that most trajectories do not present any transition, and those that do tend to have a small number of transitions. The vertical axis shows the frequency of each number of transitions per trajectory, while the horizontal axis the number of transitions per trajectory. These plots help to assess the inter-basin dynamics.

**Figure S32:**
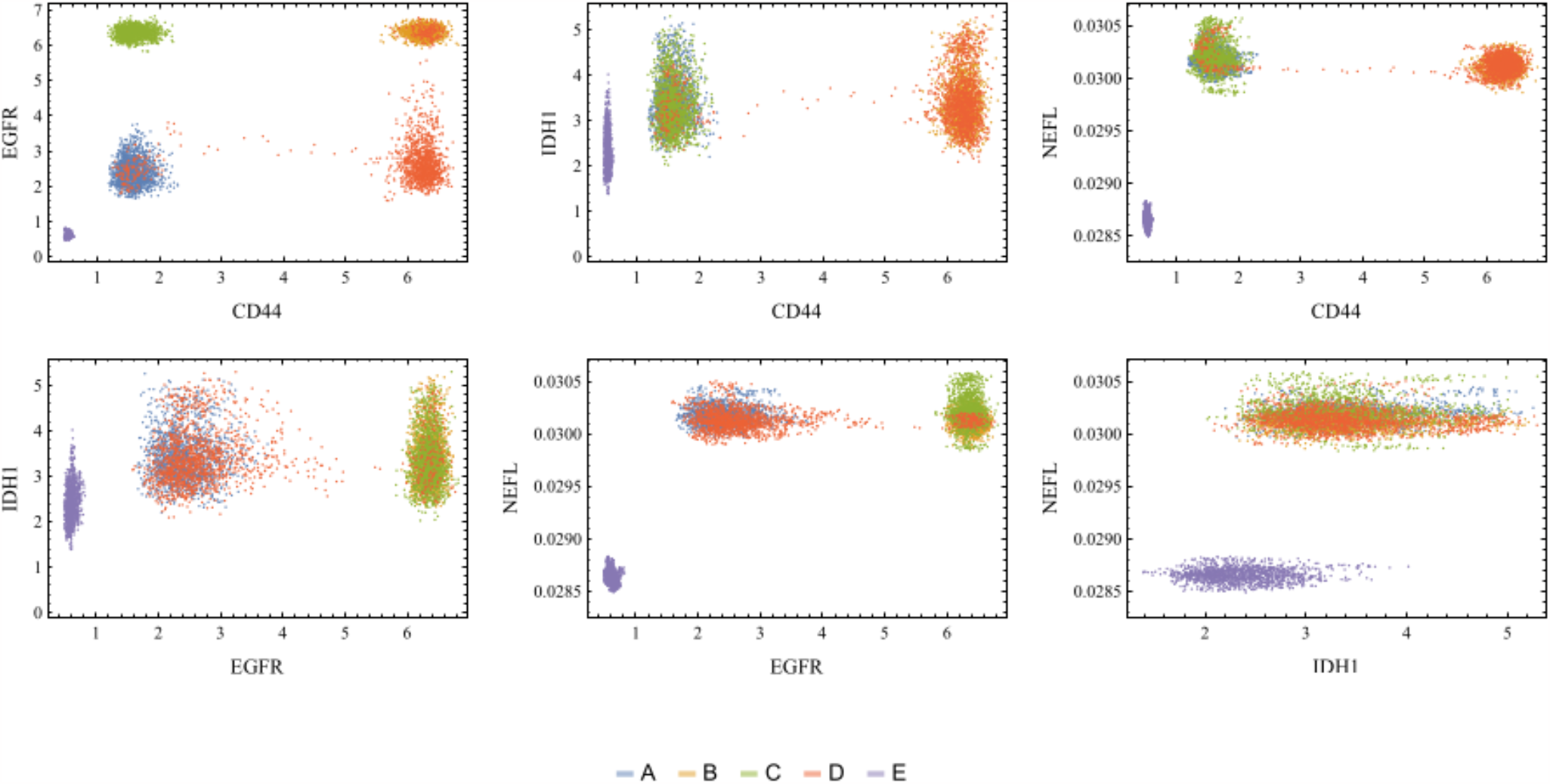
Two-dimensional visualization of the trajectories in the markers combination space. The horizontal and vertical axes display the expression values of each marker gene. Each point represents a single time step of three considered trajectories. Each color/letter indicates its respective basin. This figure helps to illustrate the trajectories’ paths and the system’s dynamics in a simplified 2D space.

**Figure S33:**
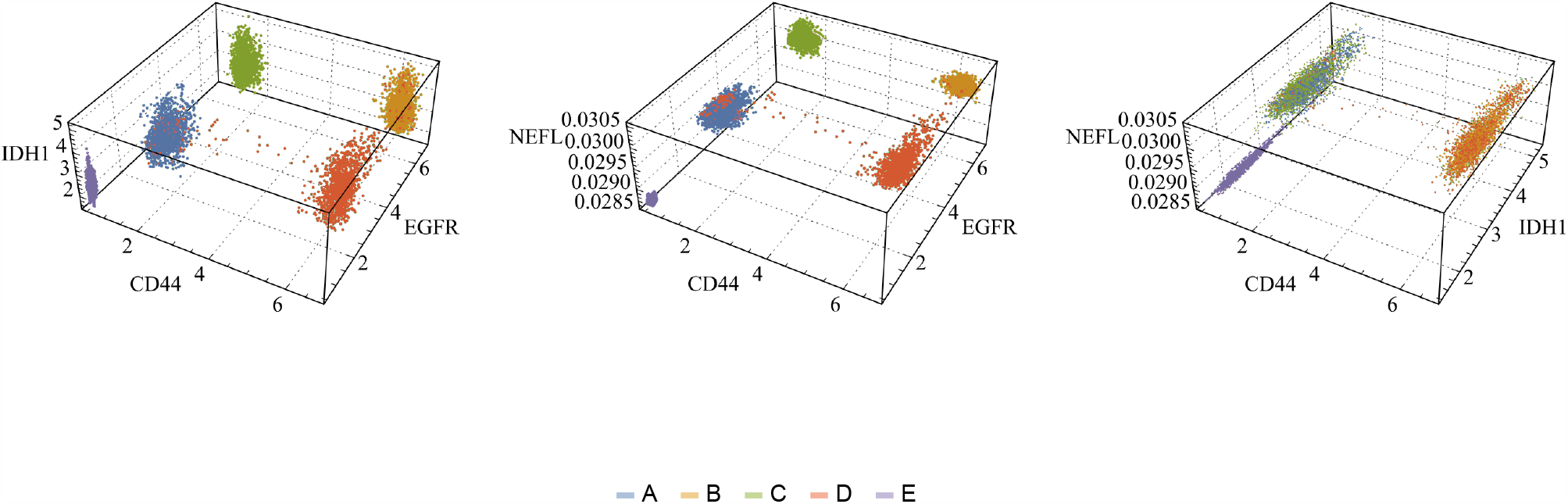
Three-dimensional visualization of the trajectories in the markers combination space. All axes display the expression values of each marker gene. Each point represents a single time step of three considered trajectories. Each color/letter indicates its respective basin. This figure offers a more detailed view of the system’s dynamics and trajectories in a 3D space.

**Figure S34:**
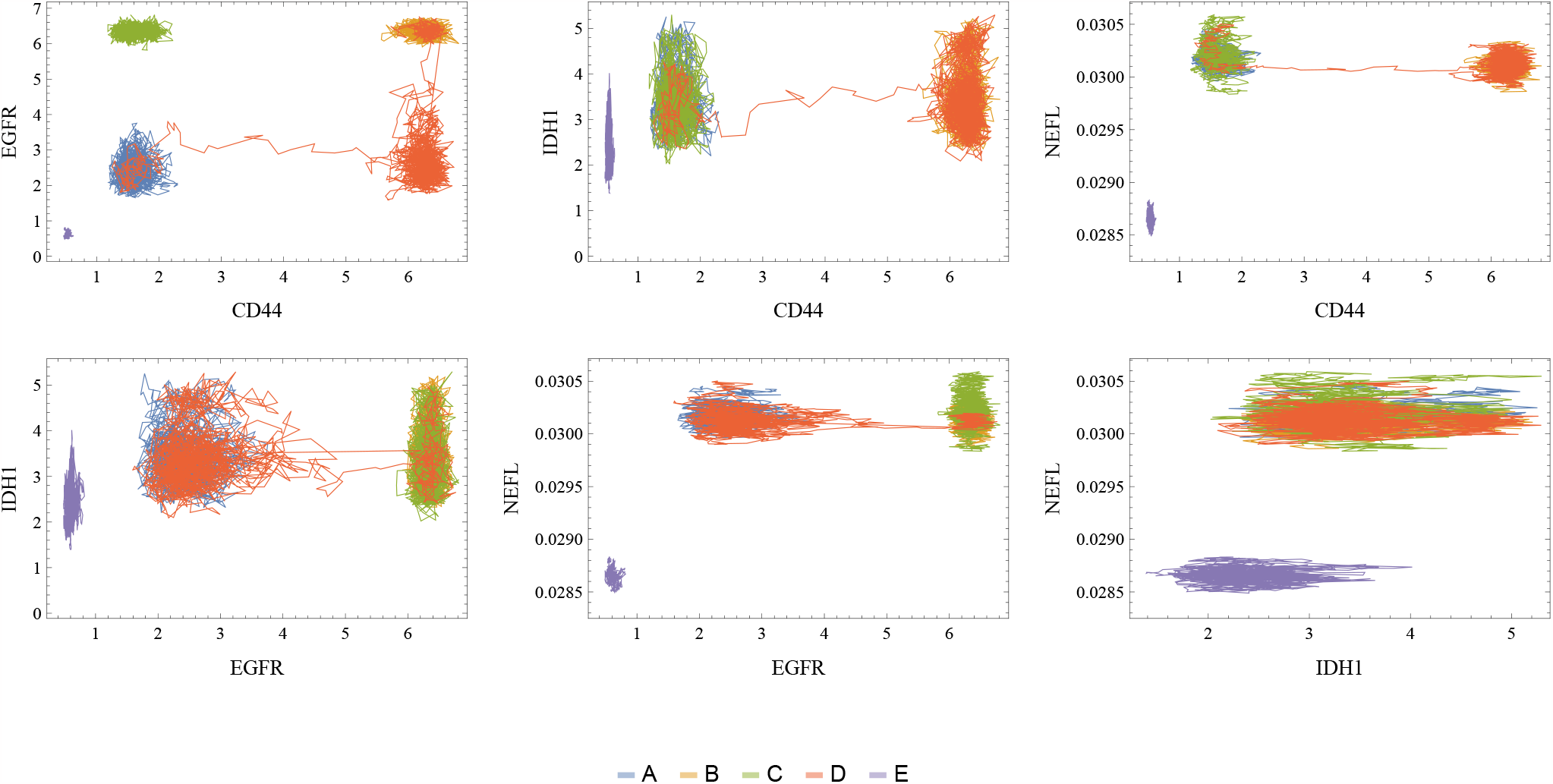
Two-dimensional visualization of full trajectories in the markers combination space. The horizontal and vertical axes display the expression values of each marker gene. Each line represents an entire time of the three considered trajectories. Each color/letter indicates its respective basin. This figure provides an overview of the paths and dynamics of the system’s trajectories in a 2D space.

**Figure S35:**
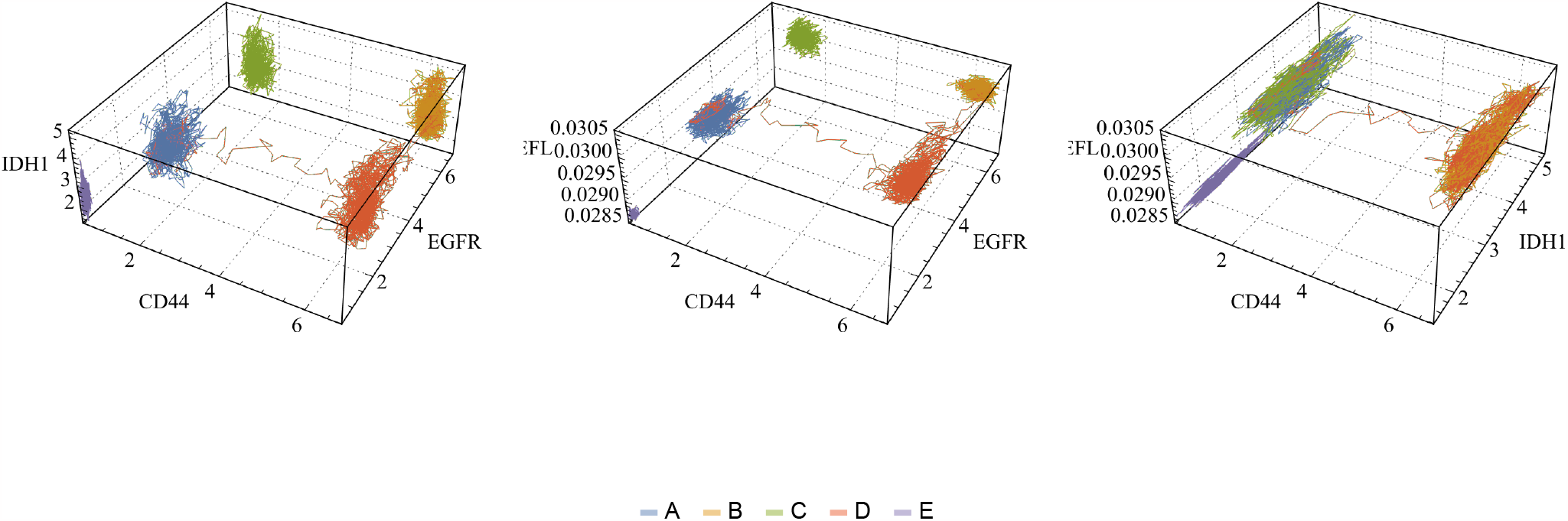
Three-dimensional visualization of full trajectories in the markers combination space. All axes display the expression values of each marker gene. Each line represents an entire time of the three considered trajectories. Each color/letter indicates its respective basin. This figure offers a comprehensive view of the system’s dynamics and trajectories in a 3D space.

**Figure S36:**
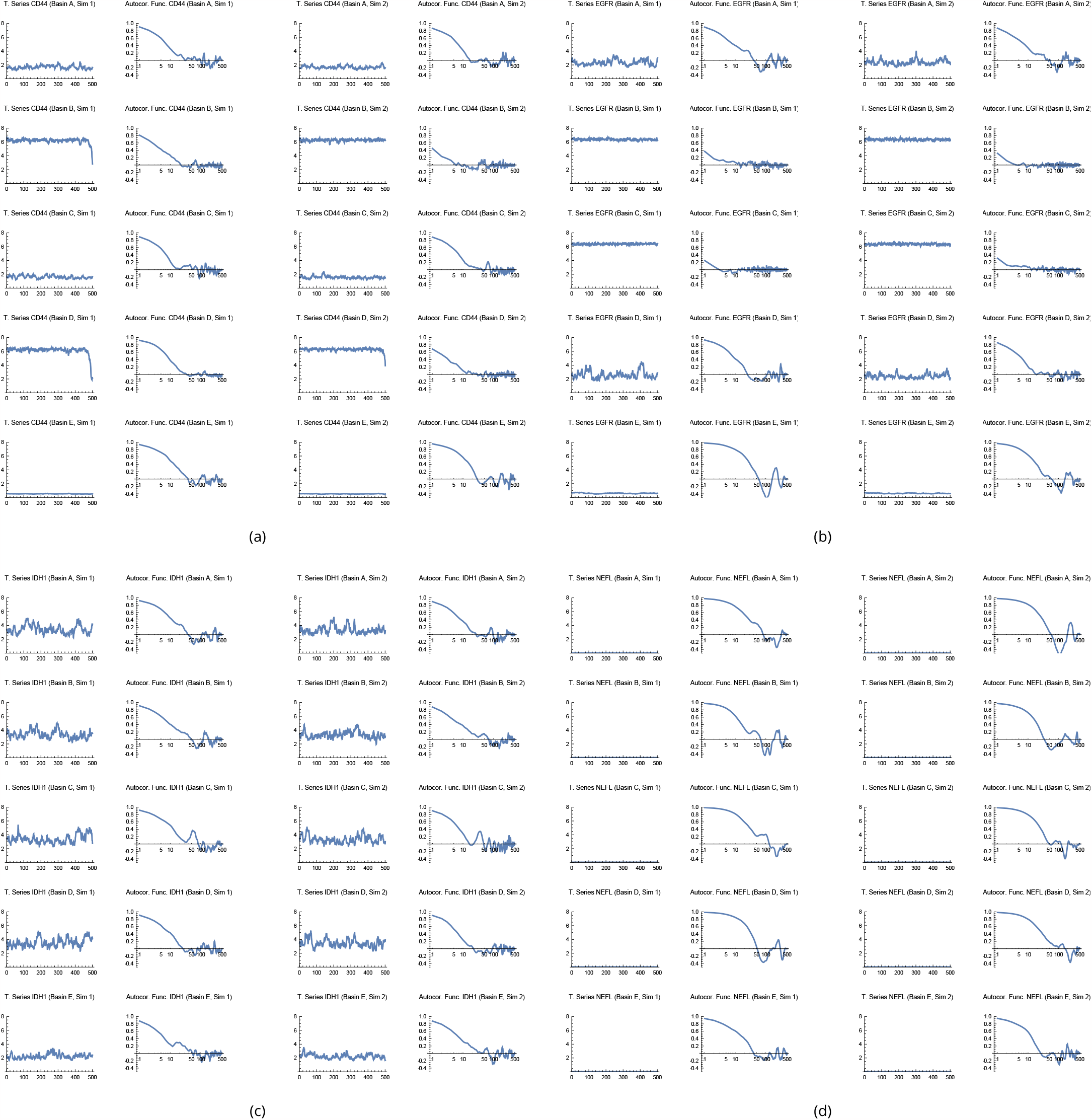
Autocorrelation analysis of time series data for different genes, basins, and repetitions. (a) to (d) Autocorrelation plots for four representative genes, illustrating the dependence structure of the time series data. Each pair of plots within (a) to (d) includes a time series plot (left) with the horizontal axis representing time and the vertical axis representing expression values and an autocorrelation plot (right) with the horizontal axis representing time lags and the vertical axis representing autocorrelation values. This figure helps to assess the temporal dependence of gene expressions and the potential impact of time series structure on the autocorrelation analysis.

**Figure S37:**
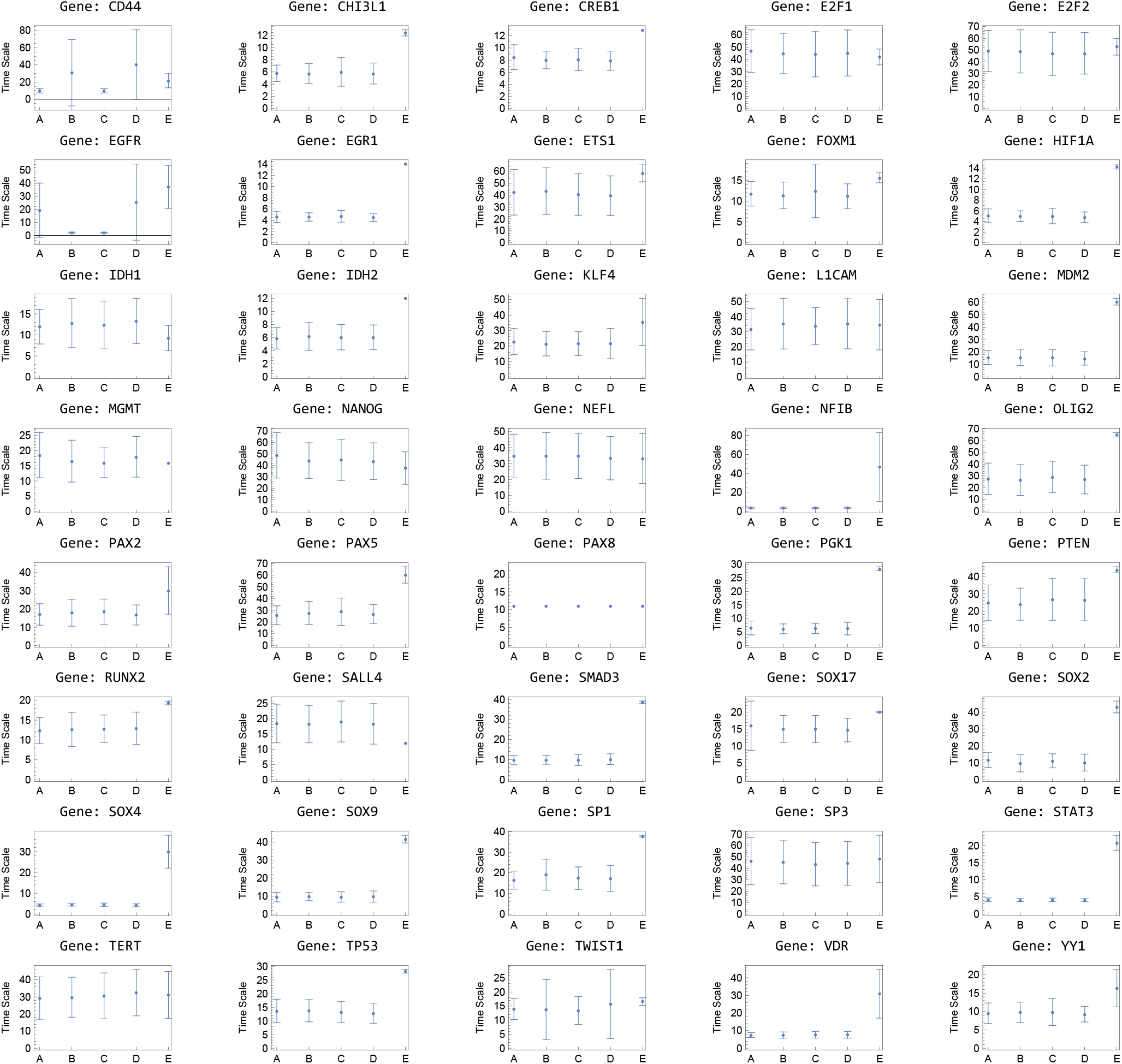
Distribution of timescales for different genes and their basins. The horizontal axis represents the cluster labels, and the vertical axis represents the timescale values, defined as the minimum time lag at which the autocorrelation falls below *e*^−1^. The figure presents the distribution of timescales for each gene, with error bars indicating the standard deviation within all repetitions. This visualization helps to understand the characteristic timescales within and across different genes, providing insights into the internal dynamics and possible transition behaviors of the system.

**Figure S38:**
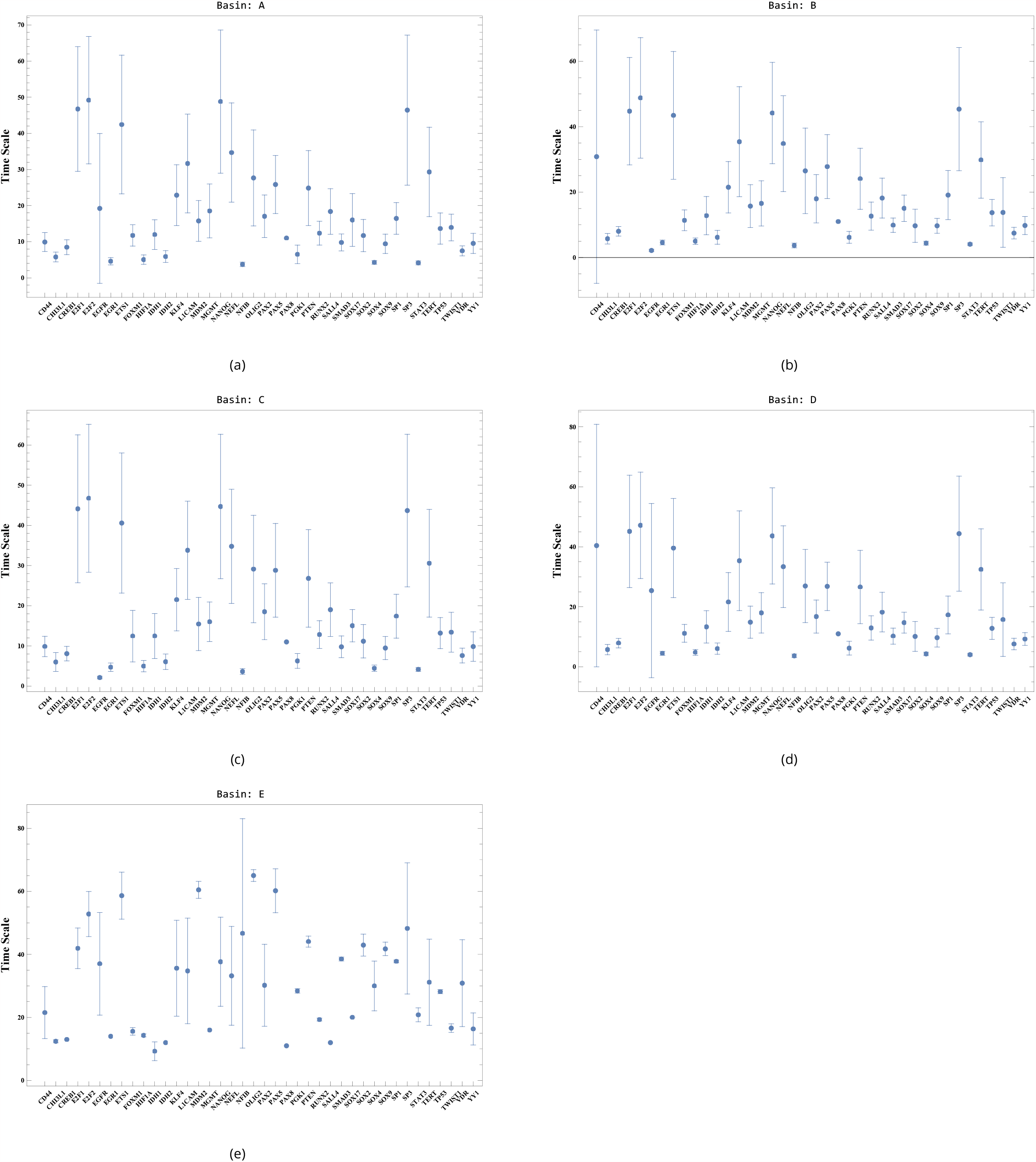
Timescale distributions for different genes within their basins. The horizontal axis represents the gene labels, and the vertical axis represents the timescale values, defined as the minimum time lag at which the autocorrelation falls below *e*^−1^. The figure emphasizes the differences within each basin, showcasing the distinct characteristics of each gene’s time-scale distribution across different basins.

**Figure S39:**
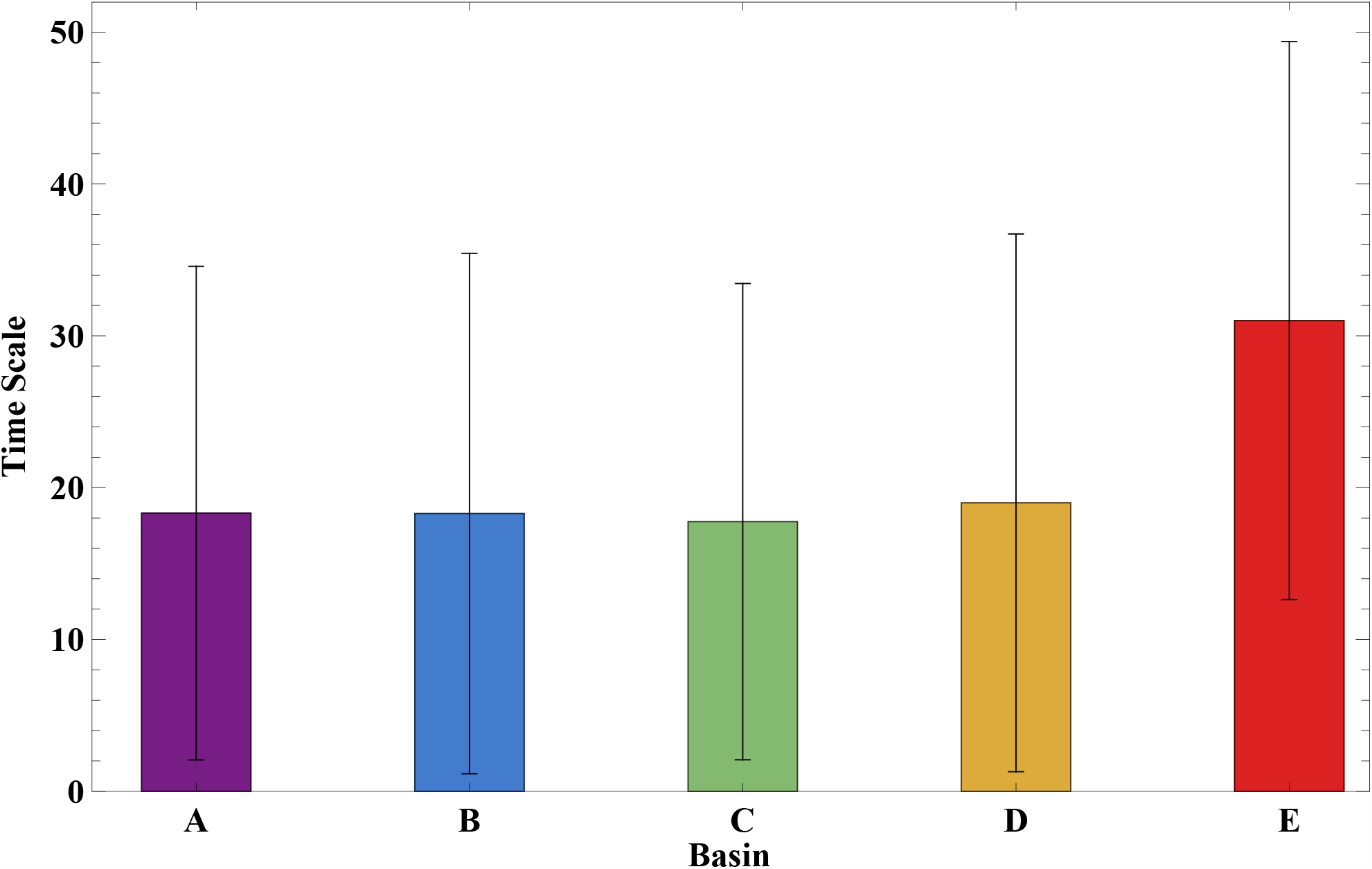
Final average timescales of each basin across all genes and repetitions. The horizontal axis represents the cluster labels, and the vertical axis represents each basin’s final average timescale values.

**Figure S40:**
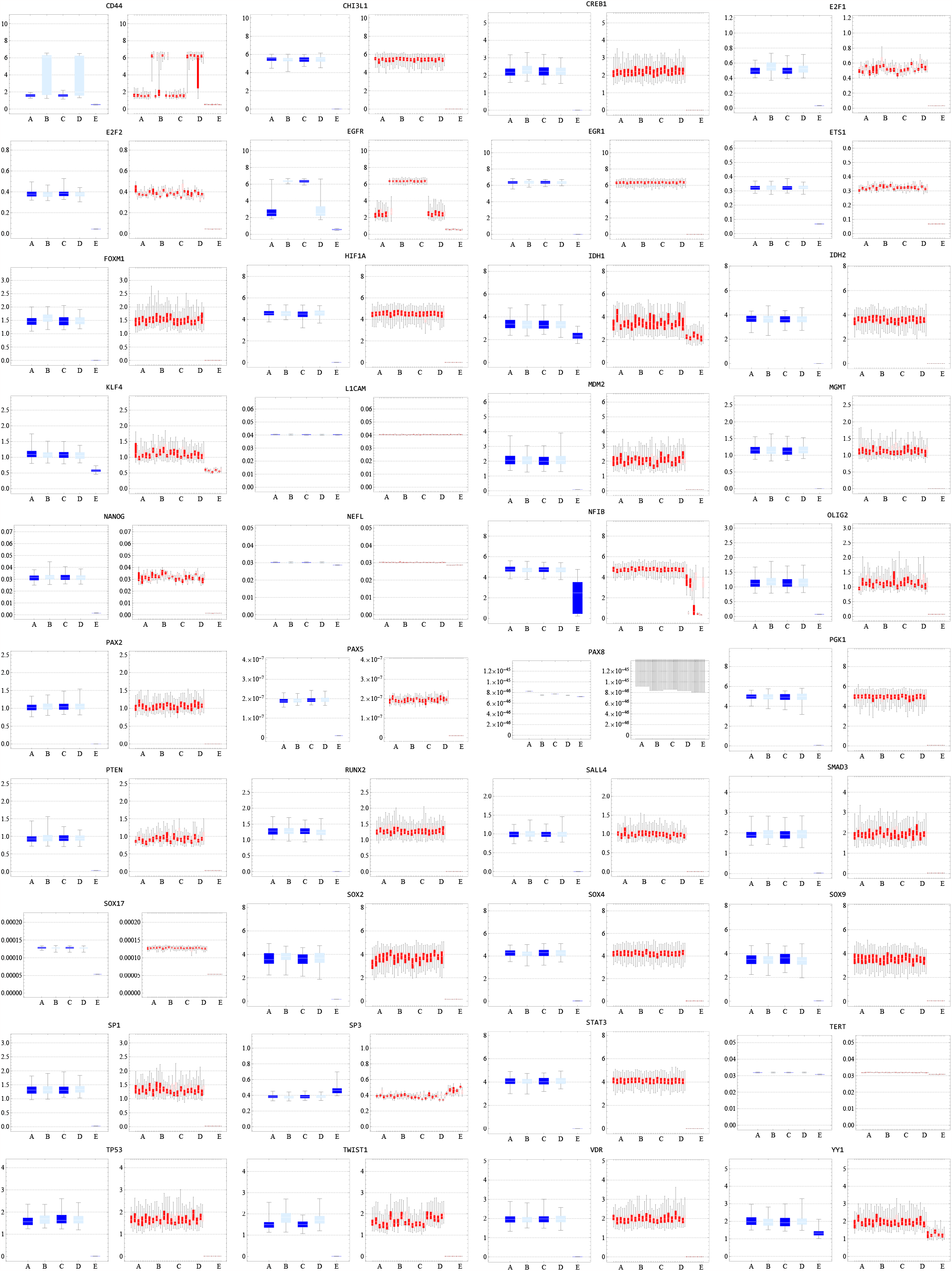
Comparison of time and sample averages for various clusters and genes using boxplots. The horizontal axis represents the cluster labels, and the vertical axis shows the expression values. The figure displays two sets of plots: the left set shows the sample average of 100 samples at the final time interval, while the right set represents the time average considering 10 trajectories from time 30 to 50 (steps 300 to 500). The boxplots illustrate the distribution of expression values within each cluster, with the box representing the interquartile range (IQR), the line inside the box showing the median, and the whiskers extending to the minimum and maximum data points within 1.5 times the IQR. This comparison helps to assess the compatibility between time and sample averages, supporting our hypothesis and providing insights into potential discrepancies between different basins and genes.

## Footnotes

1 Since the RNA sequencing data provide an estimate of expression after genetic and epigenetic regulations, and it is not possible to verify the contributions of each explicitly, we chose to use parentheses in (epi)genetics.

2 Differences in the interval resulting in approximately 1% changes in the indicator values were disregarded.

